# A TWAS method calibrated for the uncertainty of predicted expression

**DOI:** 10.1101/2022.11.06.515378

**Authors:** Arunabha Majumdar, Tanushree Haldar

**Affiliations:** Department of Mathematics, Indian Institute of Technology Hyderabad, Kandi, Telangana 502285, India; Institute for Human Genetics, University of California San Francisco, San Francisco, California 94143, USA

**Keywords:** Adaptive Lasso, Gene expression, Gene-phenotype associations, Penalized regression, Residual bootstrap, Trimmed mean

## Abstract

The transcriptome-wide association study (TWAS) is a powerful approach to identifying novel genes associated with complex phenotypes. Standard TWAS approaches build a prediction model for the genetic component of expression based on reference transcriptome data. Next, an outcome is regressed on the predicted expression in separate GWAS data. The traditional TWAS approach disregards the uncertainty of predicted expression, which can lead to unreliable inference on gene-phenotype associations. We propose a novel approach that adjusts for the uncertainty of predicted expression in TWAS. We adapt techniques from measurement error theory and implement bootstrapping algorithms for penalized regression to explicitly obtain an adjustment factor to be incorporated into the unadjusted TWAS. We base the framework on adaptive Lasso. Using extensive simulations, we show that our approach produces more accurate estimates of the gene’s effect size than a traditional TWAS approach. Traditional TWAS marginally inflates the type I error rate, whereas the adjusted TWAS adequately controls it. At the expense of a modestly inflated false positive rate, an unadjusted TWAS offers a limited increase in power compared to the adjusted TWAS. Simulations show that some other unadjusted and unified TWAS approaches can also inflate the type I error rate, particularly when using summary-level data. We demonstrate the merits of the proposed adjusted approach by conducting TWAS for height and lipid phenotypes, LDL and HDL cholesterol, and triglycerides while integrating the Geuvadis transcriptome and UK Biobank GWAS data.

## 1 Introduction

Genome-wide association studies (GWAS) have successfully associated single-nucleotide polymorphism (SNP) loci with various complex human phenotypes. The tremendous success in discovering SNP associations has inspired translational medicine studies to strive for improving precision medicine for many diseases, bringing genetic discoveries into service [1]. However, most of the GWAS signals reside in the non-coding regions of the human genome [2], lacking a comprehensive understanding of molecular mechanisms underlying the associations. The missing link between the susceptibility SNPs and the genes impacting the phenotype inhibits the translation of GWAS findings to clinical settings [1]. Non-coding regions of the human genome, e.g., intergenic regions, can regulate genes’ transcriptional or translational activities, termed regulatory elements of gene expression. An expression quantitative trait locus, eQTL, explains a fraction of the variance of a gene’s expression. Thus, eQTLs comprise various categories of regulatory elements, e.g., enhancers, promoters, transcription factor binding sites, etc. Various studies [3] demonstrated that GWAS-identified SNPs are more likely to be eQTLs. Therefore, a genetic variant identified by GWAS can influence the phenotype by modulating the expression level of its neighboring gene. This crucial speculation motivated the development of the transcriptome-wide association study (TWAS).

TWAS [4, 5] aggregates regulatory effects of multiple eQTLs and tests for association between a gene and an outcome phenotype. It combines the genotype and outcome phenotype data from GWAS with reference transcriptome data. First, we build a prediction model for the genetic component of expression for a gene based on reference transcriptome data. Next, we predict the genetic component of expression in separate GWAS data using the prediction model. Then, an outcome phenotype is regressed on the predicted expression in the GWAS data to test for gene-phenotype association (see supplementary materials for an example). TWAS has discovered many novel gene-phenotype associations and reconfirmed associations between numerous genetic loci and phenotypes initially identified by GWAS [6,7].

A significant criticism of the traditional TWAS approach is that it disregards the uncertainty of the predicted expression in the second-step regression. The uncertainty arises due to the variance of the prediction model for the expression estimated in the reference transcriptome data. A recent study [8] demonstrated that the TWAS inference on gene-phenotype association, more specifically, on the relationship between the genetic component of expression and the outcome, may not be reliable due to the non-adjustment of the uncertainty in the predicted expression. The false positive rate can be inflated, thereby reducing the accuracy of detected gene-phenotype associations. Therefore, extensive effort and valuable resources to reveal the biological mechanisms underlying the TWAS associations can be unsuccessful in follow-up functional studies. Despite considerable benefits and advantages over GWAS, TWAS needs to overcome this limitation. We term the commonly used methods for TWAS that ignore the uncertainty of predicted expression as unadjusted TWAS.

We develop a frequentist framework to fine-tune the uncertainty of predicted expression in TWAS explicitly. We adapt measurement error theory [9] to derive the adjustment factor that has to be multiplied by the test statistic obtained from an unadjusted TWAS to calibrate the uncertainty of imputed expression. We apply bootstrapping techniques for penalized regression [10] to estimate the adjustment factor and devise the procedure for testing gene-phenotype association. We base the approach on the adaptive Lasso [11] and refer to it as PenaTWAS (**P**redicted **e**xpression **n**oise **a**djusted TWAS). We develop the method using both individual-level and summary-statistics data from a GWAS [4]. Therefore, it can easily be applied to complex phenotypes using publicly available summary-level association data from GWAS. We modify an adjusted variance approach proposed by Xue et al. [12] to adjust for the uncertainty of predicted expression, which requires individual-level data.

Using simulations for individual-level data, we compare PenaTWAS with the adjusted variance approach concerning type I error rate and power. Next, we compare PenaTWAS with the unadjusted TWAS approaches regarding type I error rate and power. We perform the comparison using both individual and summary-level GWAS data. In PenaTWAS, we obtain the adjustment of the gene-outcome effect size estimate. We evaluate the estimation accuracy of the true effect sizes by PenaTWAS compared to the unadjusted TWAS approaches. Even though we developed the approach for a continuous phenotype, we also applied the same adjustment factor in PenaTWAS for a binary case-control phenotype.

A few recent approaches proposed joint modeling of the transcriptome and GWAS data to calibrate the uncertainty of predicted expression: CoMM [13] and PMR-Egger [14]. CoMM is based on a collaborative mixed effects model, and PMR-Egger uses a combined model. Both methods make a polygenicity assumption regarding the effects of local SNPs on gene expression: all local SNPs are eQTLs and have non-zero effects on the expression. On the contrary, PenaTWAS models the sparsity in the local SNPs’ effects on gene expression: not all but a subset of local SNPs are eQTLs. We compare PenaTWAS with CoMM and PMR-Egger for individual-level data, and PMR-Egger for summary-level data. We then compare PenaTWAS with another class of methods: HMAT [15] and Otters [16], which combine the results obtained by TWASs based on different expression prediction models. While HMAT uses harmonic mean aggregation [17] to combine the TWAS p-values, Otters employs Cauchy aggregation [18]. In real data applications, we implement the summary PenaTWAS, unadjusted summary-TWAS using elastic net regression and Otters for height, HDL and LDL cholesterol, and triglycerides, integrating Geuvadis transcriptome and UK Biobank GWAS data.

## 2 Materials and methods

We present a schematic diagram outlining the main steps of the proposed method in Figure 1. We provide a summary of notations and details on the mathematical derivations in the supplementary materials.

**Figure 1:**
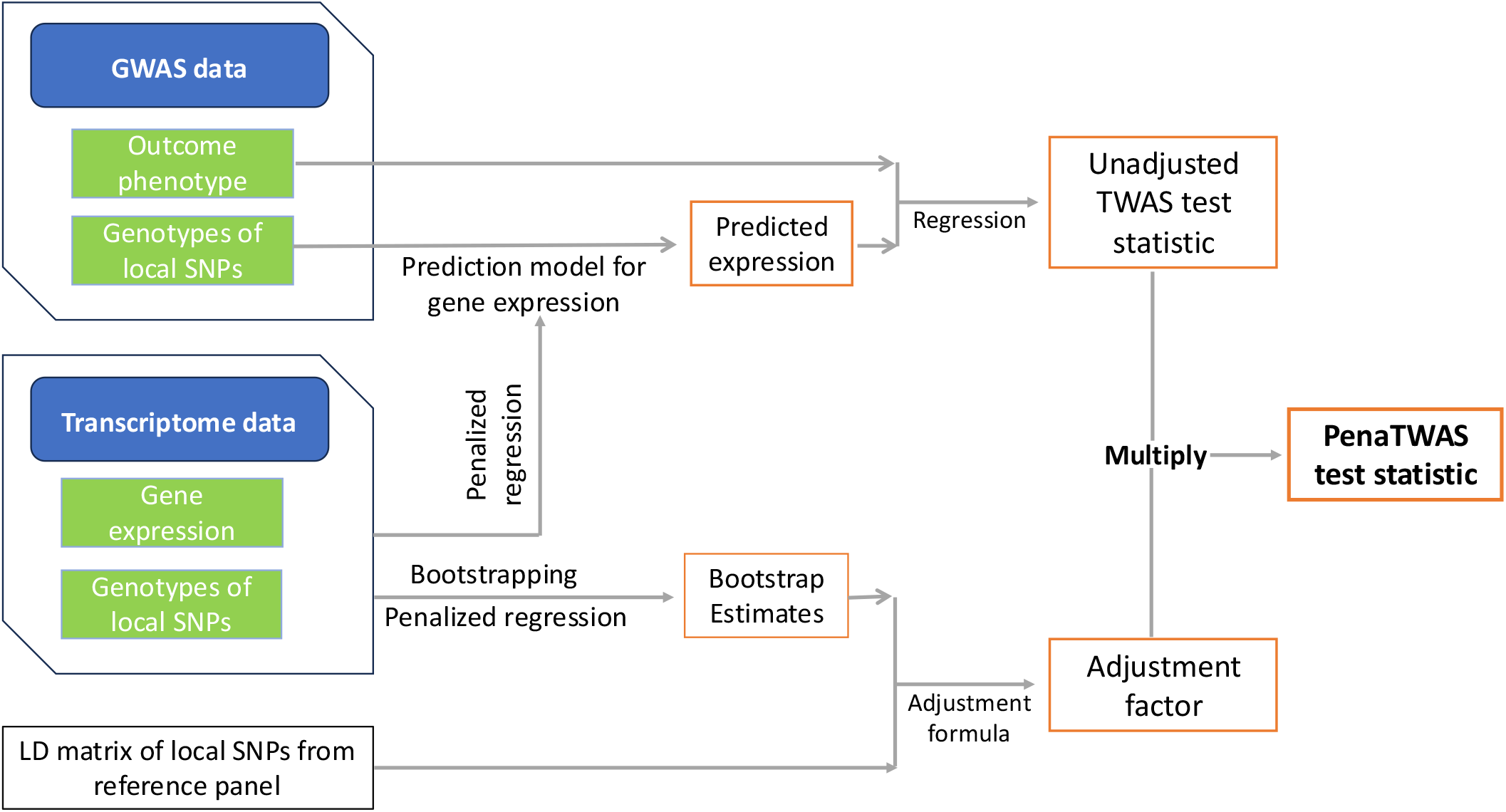
Flow chart of different steps in the adjusted method PenaTWAS. The left side of the diagram shows the three main components of the input data. First, a penalized regression of gene expression on local SNP genotypes is fitted to obtain a prediction model for gene expression (middle part of the chart). We use this prediction model to obtain predicted expression from the GWAS data (upper-middle). Next, we obtain the unadjusted TWAS test statistic by regressing the outcome phenotype on imputed expression in the GWAS data. We perform bootstrapping for penalized regression based on the transcriptome data to generate bootstrap estimates of the local SNPs’ effects on expression (lower panel). We combine these with the reference panel LD matrix to get the adjustment factor, which accounts for uncertainty in predicted expression (lower panel). Finally, we multiply the adjustment factor by the unadjusted TWAS statistic to obtain the PenaTWAS test statistic (right panel). The orange-outlined boxes represent the outputs generated during model fitting. When using GWAS summary statistics, we do not need individual-level GWAS data.

### 2.1 Regression of gene expression on local SNPs

Consider a given gene and its expression in a tissue of interest in the reference transcriptome data. Select a set of local SNPs around the gene. We regress the gene’s expression on the genotypes of the local SNPs. Consider the following linear model:

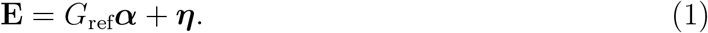

**E** denotes the vector of expression values for a gene for *n*_1_ individuals. In *G*_ref_, the genotype data of local SNPs in the reference data, genotypes for each SNP in a column are centered for the mean and then divided by the standard deviation. ***α*** = (*α*_1_, …, *α*_*p*_)′ denotes the vector of effect sizes of *p* local SNPs on the gene expression, and ***η*** denotes the vector of residuals. The residuals are independent and normally distributed with mean zero and variance 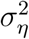. Since the number of local SNPs can be larger than the sample size, a penalized regression is implemented to estimate ***α***, e.g., Lasso [19] and Elastic net [20]. In Lasso, 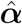 is obtained by minimizing the penalized loss function:

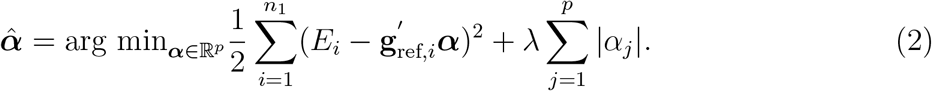

Here, *E*_*i*_ and **g**_ref,*i*_ denote the gene expression and genotype vector for *i*^*th*^ individual, respectively, *i* = 1, …, *n*_1_. *λ* ≥ 0 denotes the weight associated with the penalty term, which determines the amount of shrinkage. In Elastic net, 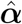 is given by:

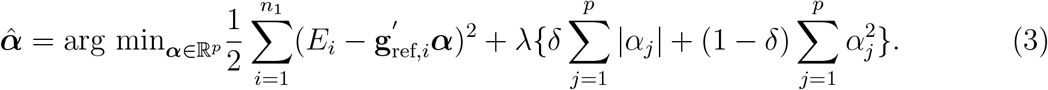

Here, *δ* ∈ (0, 1) determines the relative contribution of the Lasso and ridge regression penalty and is chosen as 1*/*2 in practice. An optimal *λ* is chosen by cross-validation. Let 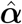 denote the estimate of ***α*** obtained by the preferred penalized regression. Let 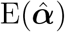 and 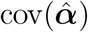 denote the expectation vector and covariance matrix of 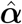.

### 2.2 Regression of phenotype on genetic component of expression

Consider GWAS data; the subjects are non-overlapping with the transcriptome data. In general, expression measurements are not available in GWAS data. Assume that the data has the genotypes for the set of *p* local SNPs considered for the gene in the transcriptome data. We assume that the transcriptome and GWAS data are from the same population, such that the effect of local SNPs on the expression is the same between the two datasets. If **g** denotes the genotype vector for the local SNPs in GWAS, the true genetic component of expression is given by *x* = **g**′ ***α***. Since ***α*** is not known, we predict *x* by 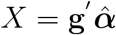, where 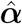 was obtained in the first-step regression (Equations 2, 3). The main goal of TWAS is to test for an association between the outcome and the true genetic component of expression. Let *Y* denote the outcome phenotype of interest and **y** = (*y*_1_, …, *y*_*n*_) be the phenotype values for the GWAS individuals. For the *i*^*th*^ GWAS individual, let **g**_*i*_ denote the genotype vector for the *p* local SNPs. Hence,

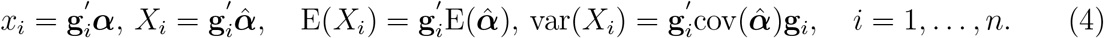

Thus, *X*_*i*_ estimates *x*_*i*_ with some measurement error or uncertainty.

#### 2.2.1 Ideal and working regressions

We consider two regressions, one considering *x* as the regressor and the other taking *X* as the regressor. Here, *x* is unobserved, and *X* is observed. We specify the ideal regression as:

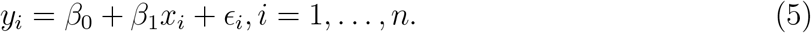

*β*_1_ is the effect of the true genetic component of expression on the phenotype. Assume that {*ϵ*_*i*_,*i* = 1, …, *n*} are i.i.d. 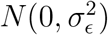. Next, the working regression is specified as follows:

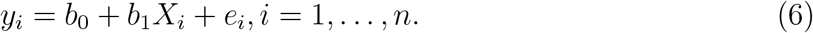

Here, *b*_1_ is the effect of the noisy predicted expression on the phenotype, and {*e*_*i*_,*i* = 1, …, *n*} are i.i.d. 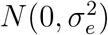. The definitions of the two regressions go along similar lines of linear regression with measurement error in the regressors [9]. TWAS is a two-stage, two-sample framework where the two samples from the transcriptome and GWAS data are non-overlapping and independent. Hence, *η*s in the Equation 1 and *ϵ*s in the Equation 5 are independent. In an unadjusted TWAS, we implement the working regression (Equation 6), ignoring the uncertainty or measurement error in *X* observations. Our main goal is to test whether the true genetic component of expression affects the phenotype, i.e., *H*_0_ : *β*_1_ = 0 vs. *H*_1_ : *β*_1_ ≠ 0. However, in unadjusted TWAS, we test for *H*_0_ : *b*_1_ = 0 vs *b*_1_ ≠ 0 using the least squares estimator (LSE) of *b*_1_, based on working regression (Equation 6):

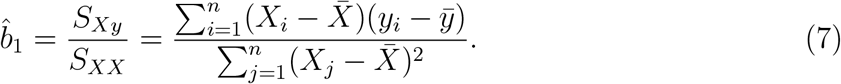

#### 2.2.2 An adjusted estimator and related test

Our main goal is to obtain an estimator of *β*_1_ in the ideal regression (Equation 5) and test for *H*_0_ : *β*_1_ = 0 vs. *H*_1_ : *β*_1_ ≠ 0. We obtain an estimator of *β*_1_ based on 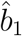. We obtain that:

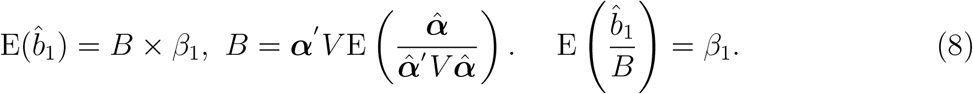

*V* denotes the GWAS linkage disequilibrium (LD) matrix of the *p* local SNPs. We regard 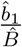 as an estimator of *β*_1_, where 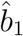 is an LSE of *b*_1_ in the working regression. The details of the above and following derivations are provided in the supplementary materials. Next, we obtain the variance of 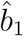 as:

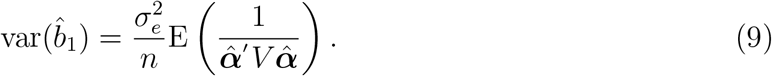

Therefore, the variance of 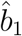 involves the error variance in working regression and the expectation of the inverse of a quadratic form, which depends on the transcriptome data and GWAS LD pattern. The uncertainty of predicted expression is summarized through this expectation, which is ignored in a standard TWAS while obtaining the variance of 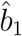. Denote the *Z* (Wald) statistic for the unadjusted TWAS as *Z*_unadj_, and that for PenaTWAS as *Z*_pena_.

Then, we obtain the relation between *Z*_pena_ and *Z*_unadj_ as:

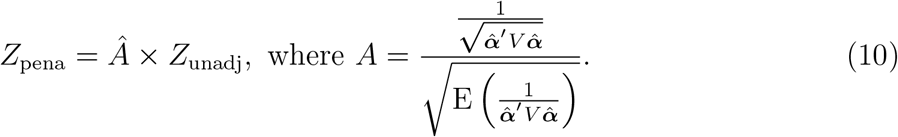

To adjust for the uncertainty of the predicted expression, we need to multiply the *Z* statistic used in the unadjusted TWAS by an estimate of the adjustment factor *A*. Intuitively, the unadjusted TWAS ignores the expectation of the quantity 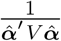 in the denominator of *A*, in which case, *A* becomes one. In the following, we discuss an approach to estimating the adjustment factor *A*. To obtain an estimate of *β*_1_, we use Equation 8:

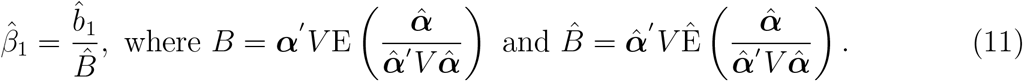

We plug in the estimate of ***α*** based on complete data: 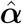. Here, 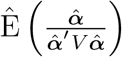 is an estimate of 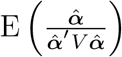.

#### 2.2.3 Estimation of the adjustment factor

We estimate the adjustment factor since it is unknown for a given gene. In the form of the adjustment factor above, 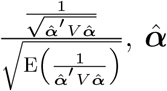 is the estimated effects of local SNPs on expression in the transcriptome data, and *V* is the LD-matrix of the local SNPs in the GWAS data. The numerator is already calculated from complete data. Deriving a closed-form formula for 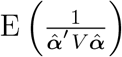 and then estimating it is very challenging. Hence, we propose an estimation approach based on bootstrapping. Note that 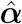 but not *V* involves variability. We perform bootstrapping for the penalized regression of expression on local SNPs in the transcriptome data (Equations 1, 2, 3). We employ the bootstrap expectation based on the bootstrapped estimates of ***α*** to estimate 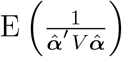. There is a considerable amount of literature on bootstrapping for penalized regressions. We adapt the residual bootstrap technique instead of a pair-wise bootstrap due to the former’s better performance, given that the linear model assumption in Equation is correct.

The bootstrapped estimates of ***α*** should correctly approximate the distribution of 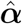, the full-sample estimate. Chatterjee and Lahiri [21] showed that the estimates of ***α*** based on the residual bootstrap for Lasso regression may not converge in distribution to 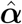. Chatterjee and Lahiri [10] proved a significant result: the residual bootstrap for adaptive Lasso [11] converges in distribution to the full sample estimate. The authors [10] proposed a version of residual bootstrap for Lasso, which converges. However, the algorithm is computationally intensive, rendering it infeasible for many genes (in the order of thousands). Hence, we develop the proposed PenaTWAS approach under the framework of adaptive Lasso [11]. The adaptive Lasso (ALasso) estimator for the local SNPs’ effects on the expression is given by:

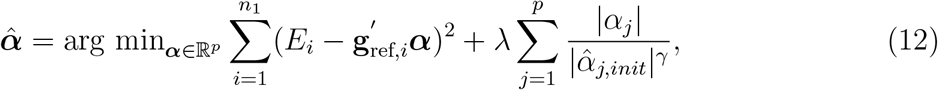

where ***α*** = (*α*_1_, …, *α*_*p*_)′, and 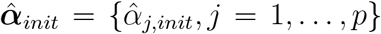 denotes the initial estimate of ***α*** based on the full sample. Here, *p* denotes the number of local SNPs, and *λ* denotes the weight associated with the penalty. The initial estimator of ***α*** must be 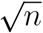 consistent [11]. The ordinary least squares (OLS) estimator satisfies this condition and is commonly used to obtain the initial estimate. In the context of TWAS, the number of local SNPs for a gene can be larger than the sample size of the transcriptome data. Some local SNPs can also be strongly correlated, i.e., in strong LD. Consequently, the OLS estimator may not be stable for many genes. On the other hand, the ridge regression estimator has the consistency property and can perform robustly in these scenarios. Hence, we implement the ridge regression to obtain the initial estimates of ***α*** instead of the OLS approach, i.e., 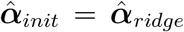. A positive *γ* allows for varying the influence of the initial estimate on the penalty term. We consider *γ* = 1 for simplicity in this work. The adaptive Lasso has some attractive statistical properties [11] and is a reliable choice for penalized regression. The ALasso allows differential shrinkage in the effects of the local SNPs on the expression. We use ten-fold cross-validation to choose the optimal *λ*. In ridge regression, we use ten-fold cross-validation, too. Next, we outline the residual bootstrap for adaptive Lasso to estimate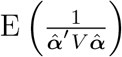, where 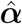 is the E full-sample ALasso estimate.

#### 2.2.4 Residual bootstrap for adaptive Lasso

As defined above, let 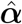 denote the full sample ALasso estimate of ***α***. Define:

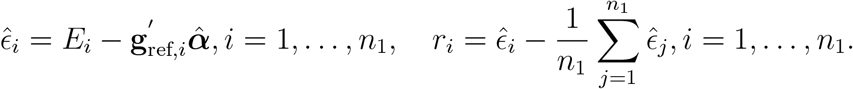

Here, 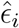 denotes the ALasso residual for the *i*^*th*^ individual in the transcriptome data, and *r*_*i*_ represents the mean-centered residual. Let 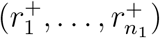 denote a random sample of size *n*_1_ drawn from 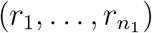 with replacement. Define:

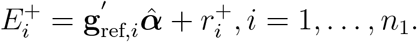

Implement ALasso for the residual bootstrap sample, 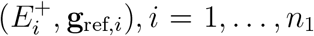, to obtain a bootstrapped estimate of ***α*** as:

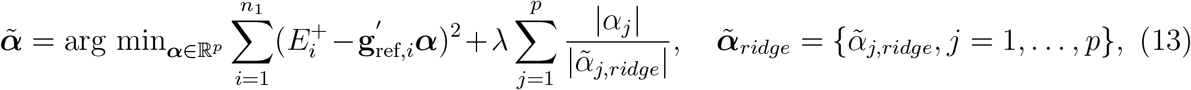

where 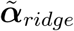 is the ridge regression estimate of ***α*** based on the bootstrap sample. Thus, 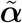 provides the bootstrapped ALasso estimate of ***α*** for a given residual bootstrap sample. We used the same choice of the penalty parameter in the ridge regression for a bootstrap sample, which was obtained from the original complete data. We used five-fold cross-validation to choose the optimal penalty parameter in ALasso for each bootstrap sample. We regard *T* residual bootstrap samples and obtain *T* bootstrap estimates of 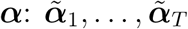. We compute 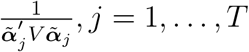, where *V* is the LD matrix of local SNPs estimated based on the GWAS data. We use the sample mean of 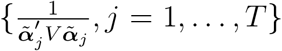 to estimate 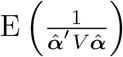. We assume that *V* is positive-definite and 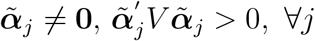.

We consider trimmed mean of 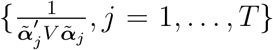 as a robust estimate of 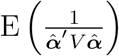. While we use the trimmed mean for the bootstrap sample to estimate the adjustment factor, we choose the option of no trimming, i.e., 0% trimmed mean, 5% trimmed mean, and 10% trimmed mean. We refer to the corresponding versions of PenaTWAS as PenaTWAS0, PenaTWAS5, and PenaTWAS10, respectively. The remaining details of the estimation of the adjustment factor, the estimation of the TWAS effect size, and a Taylor expanded version of the PenaTWAS approach are described in the supplementary materials.

### 2.3 Summary statistics version of PenaTWAS

It may often be challenging to get access to individual-level GWAS data. Gusev et al. [4] proposed a summary statistics version of TWAS (summary-TWAS), which only uses GWAS summary statistics. As a result, it can easily be implemented for various phenotypes based on publicly available GWAS summary statistics. We adjust the *Z* statistic from the unadjusted summary-TWAS. The following outlines the derivation of the unadjusted summary-TWAS *Z* statistic. Consider the estimated vector of local SNPs’ effects on expression, 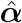, obtained from the ALasso regression (Equation 12) for Equation 1. Next, consider the phenotype regression on the predicted genetic component of expression in GWAS employing the working regression (Equation 6).

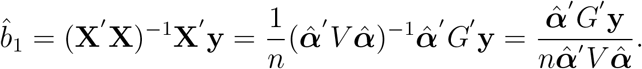

*G* denotes the genotype data matrix of the local SNPs in GWAS. Suppose the outcome *Y* is also individually regressed on the genotype of *j*^*th*^ local SNP in GWAS:

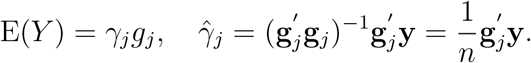

*γ*_*j*_ is the marginal effect of the SNP on the phenotype in GWAS, and **g**_*j*_ denotes the normalized genotype vector of the *j*^*th*^ SNP. Let ***γ*** denote the vector of marginal GWAS effects of the local SNPs on the outcome. We obtain the following:

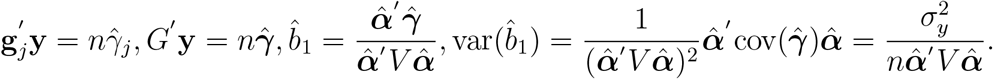

Note that 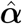 is considered non-random in the unadjusted TWAS. Here, 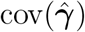 can be simplified as 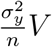, where *V* is the GWAS LD matrix of the SNPs. Thus, the unadjusted TWAS statistic *Z*_unadj_ can be formulated as 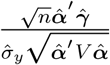. The *Z* statistic for marginally testing *H*_0_ : *γ*_*j*_ = 0 vs *H*_1_ : *γ*_*j*_ ≠ 0 in the GWAS is given by:

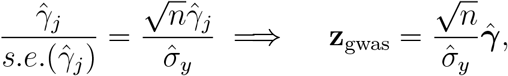

where **z**_gwas_ is the vector of *Z* statistics for testing the marginal GWAS effects. The *Z* statistic for testing *H*_0_ : *b*_1_ = 0 versus *H*_1_ : *b*_1_ = 0 in the unadjusted summary-TWAS is given by:

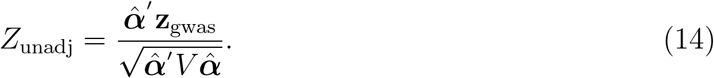

To adjust for the uncertainty of predicted expression in unadjusted summary-TWAS, we consider *Z*_pena_ = *Â*× *Z*_unadj_ (Equation 25), where *Â* is the estimated adjustment factor described in previous sections and *Z*_unadj_ is the unadjusted summary-TWAS *Z* statistic. This summary version of PenaTWAS requires summary-level GWAS association data. The approach can be implemented using publicly available GWAS summary statistics and the prediction model for expression based on ALasso and the adjustment factor. Without individual-level data from GWAS, we can use 1000 Genomes data to estimate the LD matrix, *V*. Due to a relatively lower sample size of reference panel genotype data, regularization of the calculated LD matrix is desirable. We implement a simple regularization strategy following truncated SVD. After the eigenvalue decomposition of the LD matrix, we discard the negative eigenvalues if present and the corresponding eigenvectors. Among the remaining, we retain the top eigenvalues explaining 99% of the total variance (the sum of the eigenvalues) and the corresponding eigenvectors. Finally, we retrieve the LD matrix based on the retained eigenvalues and eigenvectors.

To test for *H*_0_ : *β*_1_ = 0 vs *H*_1_ : *β*_1_ ≠ 0 in the ideal regression (Equation 5), we require the expectation and variance of *Z*_pena_ under *H*_0_. We show that 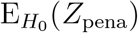 is zero and 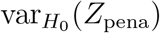 is one (supplementary materials). We provide the arguments for choosing the null distribution of *Z*_pena_ as *N* (0, 1) in the supplementary materials and simulation results next.

Xue et al. [12] proposed an adjusted TWAS approach based on individual-level data. They considered an adjusted variance of the estimated TWAS effect size to tune for the uncertainty of predicted expression. We consider a modified version of the adjusted variance approach described in the supplementary materials.

## 3 Simulation study

We performed extensive simulations to evaluate PenaTWAS compared to the unadjusted and other adjusted approaches. We compared the type I error rates and powers estimated by the different methods. We contrasted the accuracy in estimating the effect sizes. For brevity, we describe an outline of the simulation setup and analysis pipeline in the following. More details can be found in the supplementary materials. We refer to different versions of the PenaTWAS approach as: PenaTWAS0, PenaTWAS5, PenaTWAS10, PENAtaylor (Taylor’s expanded version of PenaTWAS). We also compare these approaches with a benchmark version of TWAS, which we refer to as Gold TWAS. It is a two-step TWAS method. In its second stage, while predicting the genetic component of expression for GWAS individuals, we use the unknown true local SNPs’ effects on gene expression. That is, we use the true genetic component of expression to perform the TWAS. Hence, this is the best possible TWAS approach; no other method can match its performance.

### 3.1 Simulation setting

We implemented the *hapgen* [22] software to simulate genotype data in the transcriptome and GWAS data. Gusev et al. [4] analyzed the transcriptome data from the Young Finish Sequencing (YFS) study to identify all the locally heritable genes in the whole blood tissue. We selected a locally heritable gene, DGCR8, on chromosome 22 from the YFS data analysis for simplicity in computation. To run *hapgen*, we considered a Finnish (European) population’s LD structure for the genomic region neighboring DGCR8. We chose the transcriptome data sample size of *n*_1_ = 500 and the GWAS data sample size of *n* = 10, 000. After generating the genotype data, we simulate the gene expressions of DGCR8 in the transcriptome data based on the local SNPs’ genotypes for the *n*_1_ individuals. We next simulate the outcome in GWAS data for *n* individuals. We selected ten choices of *β*_1_, the effect of the genetic component of gene expression on the GWAS phenotype, as 0.02, 0.04, …, 0.18, 0.2 to study power and estimation of effect sizes, and *β*_1_ = 0 for evaluating the type I error rate. To simulate a case-control phenotype, we threshold the continuous phenotype simulated. We provide the remaining details of the simulation setting in the supplementary materials.

### 3.2 Analysis pipeline

We implemented the GCTA software [23] to compute the p-value of testing the null hypothesis that the local heritability of the gene is zero versus the alternative hypothesis that it is positive. We applied the BH FDR controlling procedure to the p-values obtained from 6,000 iterations under a given simulation scenario. In a simulation scenario, we considered an iteration for downstream analysis if the gene was heritable in that iteration. For unadjusted TWAS, we implemented penalized regression using the Lasso, adaptive Lasso, and Elastic net penalty to estimate the prediction model for the genetically regulated component of expression based on the transcriptome data. We obtained the adjustment factor for PenaTWAS by applying the adaptive Lasso and residual bootstrapping. To assess power and type I error rate, we considered the standard choice of significance level as 0.05. We examined 0%, 5%, 10% trimmed means of the bootstrapped estimates while estimating the denominator of the adjustment factor in PenaTWAS. We denote the PenaTWAS that utilizes 0%, 5%, 10% trimmed means as PenaTWAS0, PenaTWAS5, and PenaTWAS10, respectively. To explore the normality of the null distribution of PenaTWAS statistics, we constructed the QQ and empirical density plots for the test statistic values under the null hypothesis of no association. We compared PenaTWAS with CoMM, PMR-Egger, and HMAT for individual-level TWAS and PMR-Egger, Otters, and HMAT for summary-TWAS. More details can be found in the supplementary materials.

### 3.3 Results

We first discuss the results for a TWAS based on individual-level data. Different PenaTWAS approaches and the adjusted variance approach control type I error rates adequately (supplementary Figure S1). While PENAtaylor produced the most conservative estimate of 0.02, the adjusted variance approach also led to a lower estimate of 0.029. On the other hand, PenaTWAS0, PenaTWAS5, and PenaTWAS10 resulted in an estimate of 0.035, 0.046, and 0.05, respectively (Figure S1). Thus, PenaTWAS5 and PenaTWAS10 produced a desirable type I error rate. In this simulation setting, the gene expression heritability was 10%. Regarding power, PENAtaylor yielded a lower power than the other versions of PenaTWAS and the adjusted variance approach (Figure S1). The adjusted variance approach results in a marginally higher power than PenaTWAS0, except for higher choices of *β* = 0.16, 0.18, 0.2, where PenaTWAS0 produces marginally higher power (Figure S1). We find that PenaTWAS5 and PenaTWAS10 consistently offer marginally higher power than the adjusted variance approach. For example, PenaTWAS5 produced a 1 − 4% power gain over the adjusted variance approach (Figure S1). PenaTWAS10 results in a slight power gain of 1% for most choices of *β* than PenaTWAS5. However, we chose PenaTWAS5 as the representative PenaTWAS approach. PenaTWAS10 inflated the type I error rate marginally in a simulation scenario for summary-level TWAS using out-of-sample LD (described later). In the following, for individual-level TWAS comparison, we mainly discuss the results obtained by PenaTWAS5, which we refer to as PenaTWAS in the subsequent sections.

Next, we compare the performances of the unadjusted TWAS approaches and PenaTWAS and contrast them with those of Gold TWAS. We refer to the unadjusted TWAS approaches as Lasso, ALasso (adaptive Lasso), and Enet (elastic net) TWAS, respectively, according to the type of penalty function used. The unadjusted TWASs produced a marginally inflated type I error rate (Figure 2). The Lasso, ALasso, and Enet TWAS led to a type I error rate estimate of 0.057, 0.056, and 0.058, respectively. The Gold TWAS produced a type I error rate of 0.052, which is lower than that of the unadjusted TWASs (Figure 2). The type I error rate estimated by PenaTWAS was 0.046. Thus, the unadjusted TWASs produced marginally inflated type I error rates, whereas PenaTWAS adequately controls them, a little lower than the Gold TWAS. The three choices of the unadjusted TWASs performed comparably in terms of controlling the false positive rate. The 95% confidence interval (CI) of the type I error rate for Lasso and Enet TWAS also marginally misses the true level of significance 0.05: [0.051, 0.063] and [0.052, 0.064] respectively. The CI for Gold TWAS includes the true level: [0.046, 0.058], which we observe in other simulation settings consistently. On the other hand, the CI for PenaTWAS includes the true level: [0.04, 0.052].

**Figure 2:**
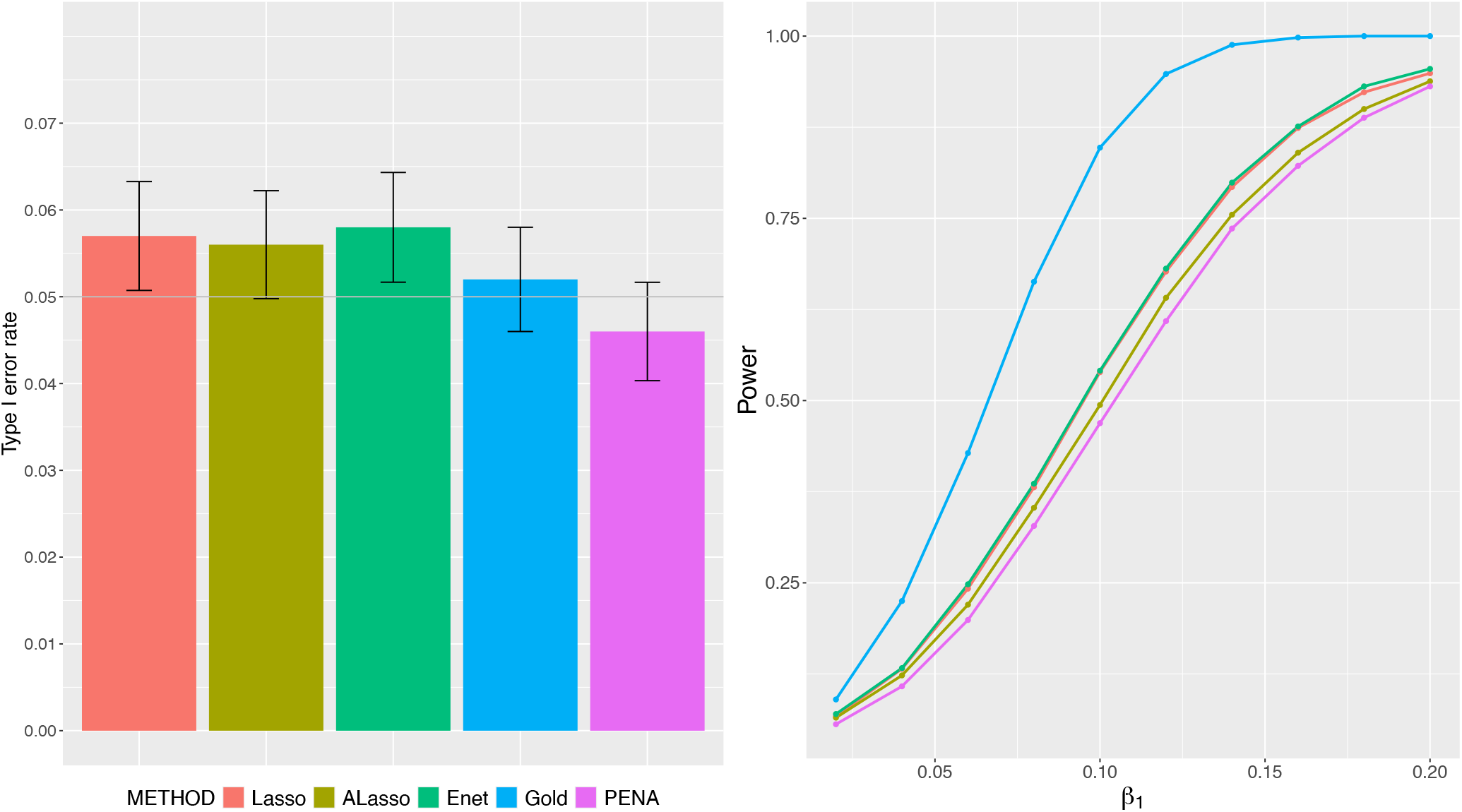
Comparison of type I error rate and power between the unadjusted TWAS approaches and the adjusted TWAS approach, PenaTWAS, based on individual-level data. The result for the gold standard TWAS, in which true eQTL effects are plugged in, is also presented. The unadjusted TWAS approaches are Lasso, ALasso (adaptive Lasso), and Enet (elastic net) TWAS, depending on the penalty function used in the penalized regression for transcriptome data. The first choice of true eQTL effects with gene expression heritability of 10% was considered. 95% CI of the type I error rate at the nominal level is also presented.

As expected, the Gold TWAS produced the maximum power, which no method could achieve (Figure 2). The Lasso and Enet TWAS yielded almost the same power, resulting in a marginal power increase of 1 − 5% than the ALasso TWAS. The nearly comparable results of these three types of unadjusted TWAS justify the development of PenaTWAS based on the adaptive Lasso framework. The unadjusted TWASs offered limited higher power than PenaTWAS at the expense of marginally inflated type I error rates (Figure 2). For simplicity, we pick the Enet TWAS to compare with PenaTWAS. Enet TWAS led to a limited higher power of 1 − 7% than PenaTWAS at the expense of a marginally inflated false positive rate, 0.046 versus 0.058. Given its calibrated type I error rate, the bearable extent of power loss by PenaTWAS is preferable.

We now describe the results for a summary-TWAS. PenaTWAS5 provides a tuned estimate of the type I error rate as 0.046 (Figure S2), whereas PENAtaylor, PenaTWAS0, PenaTWAS10 yielded estimates of 0.023, 0.042, 0.05, respectively. Regarding power, PenaTWAS10 offered a slight increase of 1% than PenaTWAS5 (Figure S2). The unadjusted summary-TWASs yielded marginally inflated type I error rates (Figure 3). The Lasso, ALasso, and Enet TWAS estimated a type I error rate of 0.056, 0.056, and 0.057, respectively. The Gold TWAS led to a type I error rate of 0.051, lower than the unadjusted TWASs. The type I error rate estimated by PenaTWAS was 0.046 (Figure 3), a little lower than the Gold TWAS. The CIs for Pena and Gold TWAS include the true level. Thus, the unadjusted summary-TWASs produced marginally inflated type I error rates again, whereas the summary PenaTWAS adequately controls it. The Gold TWAS, based on summary-level data, produced the maximum power, as expected (Figure 3). Lasso, ALasso, and Enet TWAS offered higher powers than PenaTWAS at the expense of marginally inflated type I error rates. The Lasso and Enet TWAS produced almost the same power (Figure 3). They offered a marginal power gain of 1 − 5% than ALasso TWAS. Enet TWAS led to a limited power gain of 1 − 7% than PenaTWAS at the expense of a marginally inflated false positive rate. Therefore, given its tuned type I error rate, PenaTWAS is preferable for a summary-TWAS despite the tolerable extent of power loss.

**Figure 3:**
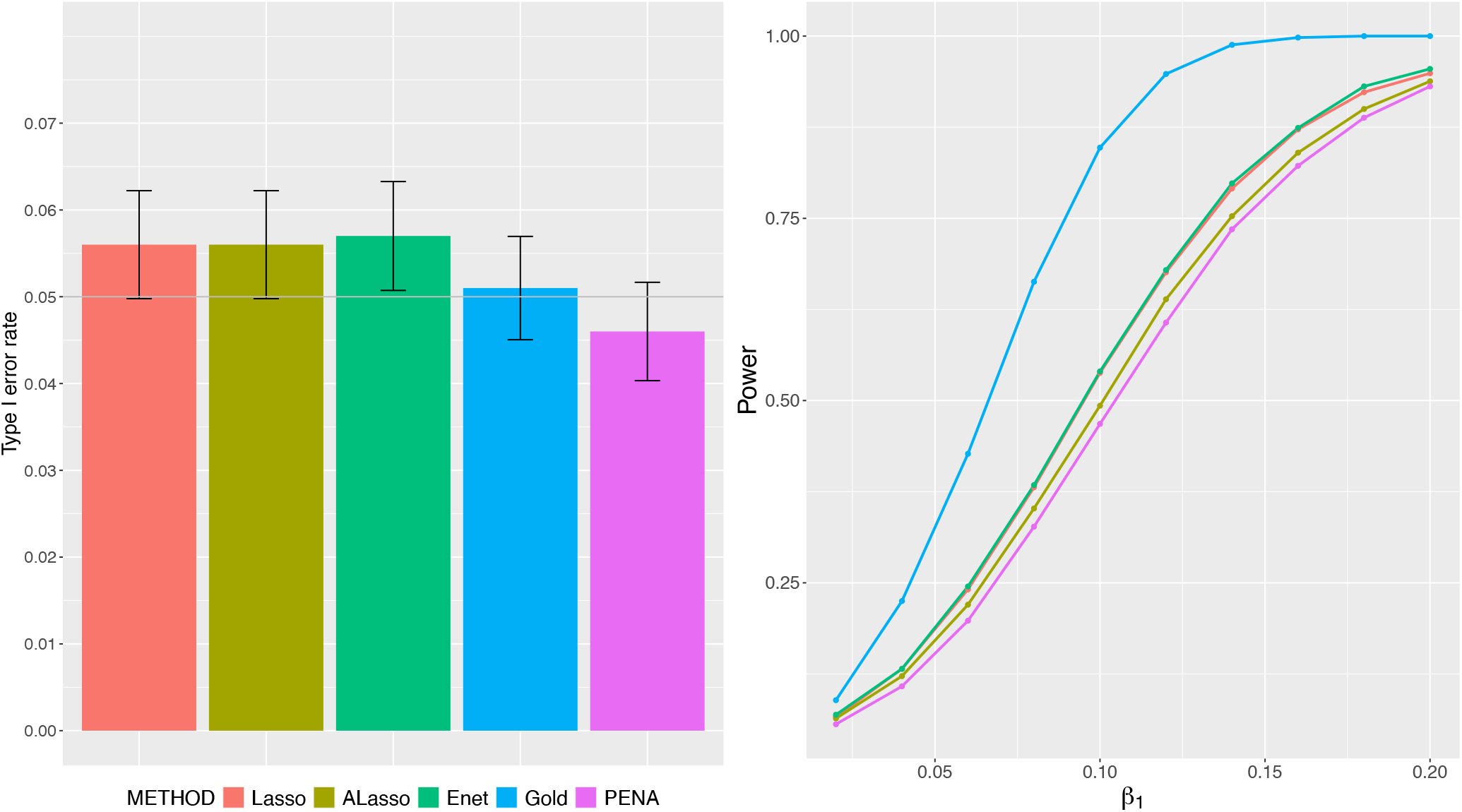
Comparison of type I error rate and power between the unadjusted TWAS approaches and the adjusted TWAS approach, PenaTWAS, based on summary-level data. Results for the gold standard TWAS, in which true eQTL effects are plugged in, are also presented. The unadjusted TWAS approaches are Lasso, ALasso (adaptive Lasso), and Enet (elastic net) TWAS, depending on the penalty function used. The first choice of true eQTL effects with gene expression heritability of 10% was considered. 95% CI of the type I error rate at the nominal level is also presented.

Next, we describe the accuracy of individual-level PenaTWAS and the unadjusted TWAS approaches while estimating the effect size of the genetic component of expression on the outcome. PenaTWAS performs well overall in estimating the effect size (Table 1). The mean of the estimated effects is the same as the true effect when *β*_1_ = 0.02, 0.04, 0.06. For the higher choices of *β*_1_, the mean falls very close to the true value, 0.01 larger than the actual value (Table 1). We utilize the relative bias, defined as 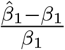, and the square root of the mean squared error (RMSE) to compare the estimation accuracy of the different approaches. ALasso TWAS produced a downward bias with a mean relative bias of −32% (Table S1). Lasso and Enet TWAS resulted in an upward bias with the mean relative bias in the range of 14%−17% and 17−23%, respectively (Table S1). PenaTWAS yielded the lowest upward bias with 6 − 7% mean relative bias. The s.d. of the relative bias is the lowest for ALasso TWAS, whereas it is comparable for the other approaches. The mean bias for a given method, excluding ALasso, decreases marginally for increasing values of *β*_1_. However, the s.d. of the relative bias decreases rapidly for all methods as the actual value of *β*_1_ increases (Table S1). Thus, PenaTWAS offered the most accurate estimates of the effect sizes concerning the relative bias. However, ALasso TWAS provided the smallest RMSE of the *β*_1_ estimates, and the other approaches yielded comparable RMSEs (Table S2).

**Table 1:**
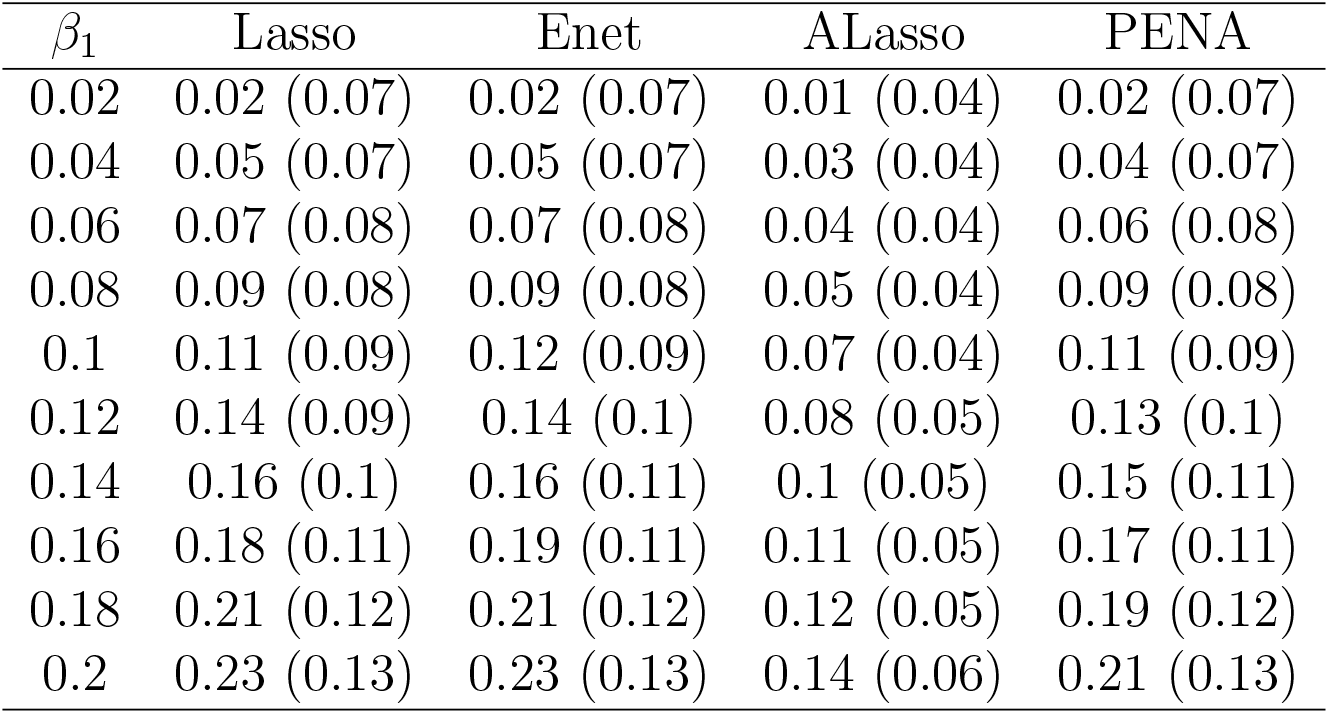
Summary of estimates of the effect size of the genetic component of expression on the outcome. The unadjusted TWAS approaches are Lasso, ALasso, and Enet TWAS, depending on the penalty function used. PENA denotes the adjusted TWAS approach, PenaTWAS. The first choice of true eQTL effects with gene expression heritability of 10% was considered. The mean and s.d. of the estimates obtained across simulation iterations are provided.

We regarded another choice of the eQTL effects with expression heritability as 10%. We repeated the above simulations to evaluate type I error rates, powers, and accuracy in effect size estimation. First, we summarize the results for the TWAS using individual-level data. We find that PenaTWAS5 performs well among the different versions of PenaTWAS concerning calibrated type I error rate and power (Figure S4). In this setting, the adjusted variance approach produces a calibrated type I error rate, and offers a power increase of 1−7% than PenaTWAS. Among the unadjusted approaches (Figure S5), Enet TWAS marginally inflated the type I error rate to 0.055; Lasso TWAS marginally overestimated it as 0.055; ALasso TWAS estimated it as 0.058. On the other hand, PenaTWAS adequately controlled the error rate as 0.045 below the desired level; the Gold TWAS correctly estimated it as 0.05 (Figure S5). Regarding power, Enet TWAS offered a 1 − 7% power increase over PenaTWAS at the expense of a marginally higher type I error rate (Figure S5). For summary-TWAS, PenaTWAS5 performs well compared to its other versions (Figure S6). We again observe the same pattern, i.e., PenaTWAS provides better control of the false positive rate at the cost of 2 − 4% loss in power compared to the unadjusted approaches (Figure S7). For the effect size estimation, PenaTWAS produced a very small upward relative bias and performed substantially better than the unadjusted approaches (Table S4). The mean estimates are the same as the true effect sizes except for a few larger choices (Table S3). PenaTWAS yielded a comparable RMSE in the estimates to the Lasso and Enet TWAS but larger than the ALasso TWAS (Table S5). Overall, PenaTWAS provides more accurate estimates of the effect sizes.

We next consider a higher choice of the expression heritability as 20% (Figure S8, S9, S10, S11). The adjusted variance approach yields a limited power gain of 1 − 7% than PenaTWAS for individual-level data-based analyses (Figure S8). For brevity, we briefly discuss the accuracy of effect size estimation using individual-level TWAS approaches and the results obtained by summary-TWAS approaches. In general, PenaTWAS controlled the type I error rate better than the unadjusted TWASs and yielded a marginal loss in power (Figure S11). PenaTWAS estimated the effect sizes more accurately than the unadjusted approaches (Table 2, S6, S7). We then chose a lower expression heritability of 5%. We observe a similar pattern, i.e., PenaTWAS better controls the type I error rate. It loses a limited power of 1 − 7% than Enet TWAS. Interestingly, in this case, the individual-level PenaTWAS yields a substantially higher power than the adjusted variance approach in the range of 3 − 17%.

**Table 2:** Summary of estimates of the effect size of the genetic component of expression on the outcome. The unadjusted TWAS approaches are Lasso, ALasso, and Enet TWAS, depending on the type of penalty function used. PENA denotes the adjusted TWAS approach, PenaTWAS. The heritability of gene expression is 20%. The mean and s.d. of the estimates obtained across simulation iterations are provided.

| $\beta_1$ | Lasso | Enet | ALasso | PENA |
| --- | --- | --- | --- | --- |
| 0.02 | 0.02 (0.03) | 0.02 (0.03) | 0.02 (0.03) | 0.02 (0.03) |
| 0.04 | 0.04 (0.03) | 0.04 (0.03) | 0.03 (0.03) | 0.04 (0.03) |
| 0.06 | 0.07 (0.04) | 0.07 (0.04) | 0.05 (0.03) | 0.06 (0.04) |
| 0.08 | 0.09 (0.04) | 0.09 (0.04) | 0.07 (0.03) | 0.08 (0.04) |
| 0.1 | 0.11 (0.04) | 0.11 (0.04) | 0.08 (0.03) | 0.1 (0.04) |
| 0.12 | 0.13 (0.04) | 0.13 (0.04) | 0.1 (0.03) | 0.12 (0.04) |
| 0.14 | 0.15 (0.04) | 0.16 (0.05) | 0.12 (0.03) | 0.14 (0.05) |
| 0.16 | 0.18 (0.05) | 0.18 (0.05) | 0.13 (0.03) | 0.16 (0.05) |
| 0.18 | 0.2 (0.05) | 0.2 (0.05) | 0.15 (0.04) | 0.18 (0.05) |
| 0.2 | 0.22 (0.05) | 0.22 (0.06) | 0.17 (0.04) | 0.2 (0.06) |

We explored whether out-of-sample LD and GWAS summary statistics used in summary-TWAS affect the type I error rate and power compared to the TWAS based on individual-level data. Interestingly, we found no consistent pattern regarding the type I error rate. For the first choice of the true eQTL effects with expression heritability 10%, the summary PenaTWAS yielded a correct type I error rate of 0.05 compared to 0.046 by its individual-level data-based version (Figure S1). Here, summary PenaTWAS10 produced a marginally inflated rate of 0.054. Conversely, for the second choice of the eQTL effects above, the estimated rate was 0.045 for both the summary-level and individual-level PenaTWAS (Figure S4), respectively. Thus, using out-of-sample LD did not affect type I error control of PenaTWAS. Regarding power, both strategies performed comparably.

For a binary case-control phenotype, the unadjusted TWASs controlled the false-positive rate nearly adequately. The Lasso, Enet, and ALasso TWAS led to estimated type I error rates of 0.052, 0.054, and 0.054, respectively. PenaTWAS curbed it below the threshold with an estimate of 0.042. Consequently, it loses a limited power of 1 − 5% compared to the Enet TWAS. PenaTWAS estimated the effect sizes more accurately (Table 3). For smaller choices of *β*_1_, the mean of the estimates coincides with the true choice. For relatively larger *β*_1_s, the mean estimates are marginally larger than the true values. The ALasso TWAS produces a substantial downward bias, and Lasso and Enet TWAS lead to significant upward bias in the estimates (Table S8). PenaTWAS yields a limited upward bias in the estimates (Table S8). PenaTWAS estimates have marginally lower RMSEs than Enet TWAS for larger values of *β*_1_ and a bit higher RMSE than ALasso TWAS (Table S9). Thus, PenaTWAS offers more accurate effect size estimates for a binary case-control outcome.

**Table 3:**
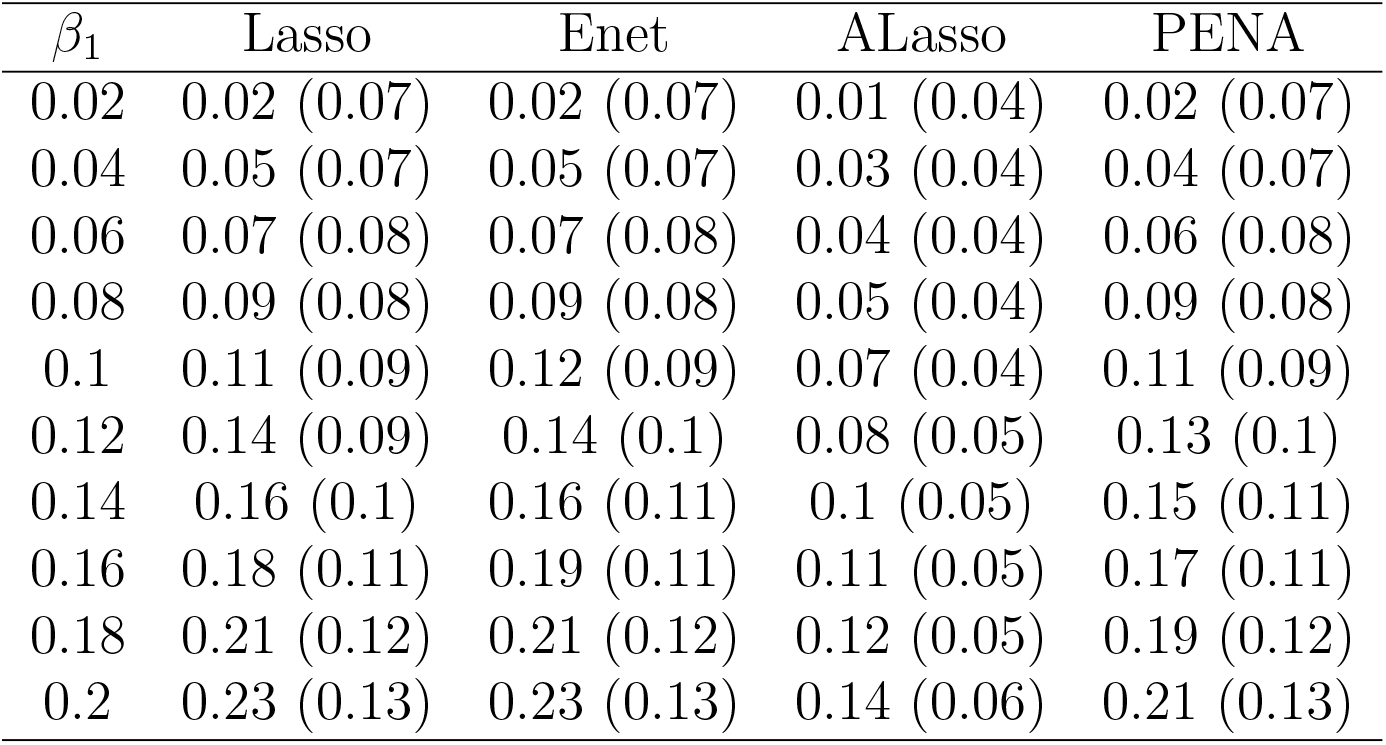
Summary of estimated effect sizes of the genetic component of expression on a binary case-control phenotype. The unadjusted TWAS approaches are Lasso, ALasso, and Enet TWAS, depending on the penalty function used. PENA denotes the adjusted TWAS approach, PenaTWAS. The heritability of gene expression is 10%. The mean and s.d. of the estimates obtained across simulation iterations are provided.

For simplicity in computation, we compared PenaTWAS, CoMM, PMR-Egger, Otters, and HMAT in the first simulation scenario for a continuous phenotype, where the gene expression heritability is 10%. Interestingly, CoMM, HMAT, and PMR-Egger, using individual-level data, control the type I error rate appropriately (Figure S3). However, for summary-TWAS, PMR-Egger, HMAT, and Otters inflate the type I error rate (Figure 4). Despite being an adjusted summary-TWAS approach, PMR-Egger inflates the false positive rate, comparable to Otters and HMAT, two unadjusted approaches (Figure 4). Thus, PenaTWAS is a more reliable alternative due to better control of the false positive rate. Consistent with the findings above, PenaTWAS loses some power compared to these approaches (Figure S3 and 4), due to its better grip on type I error control.

**Figure 4:**
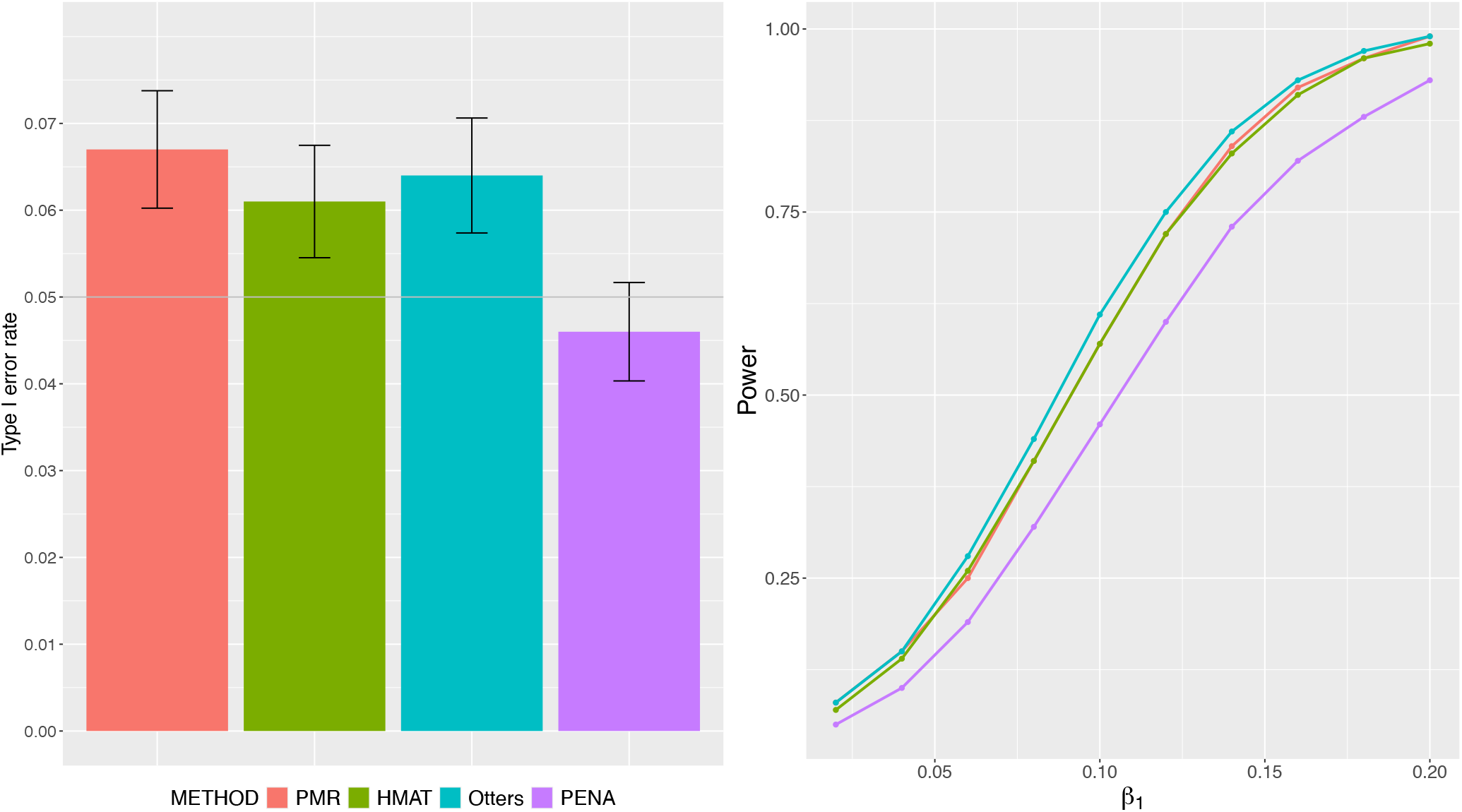
Comparison of type I error rate and power between summary-level PenaTWAS, PMR-Egger, HMAT, and Otters. PenaTWAS and PMR-Egger calibrate the uncertainty of predicted expressions. HMAT and Otters ignore the uncertainty in predicted expressions. The heritability of gene expression is 10%. 95% CI of the type I error rate at the nominal level is also presented.

For various methods, we provide QQ plots for null p-values with the 95% confidence bands (Figures 5, 6). We observe that for both Lasso and Enet TWAS, the QQ plots remain within the confidence band, with a marginal inflation near the tail of the null p-value distribution (Figure 5). The QQ plots move closer to the upper boundary on the right side. In comparison, the QQ plot for PenaTWAS demonstrates good control, albeit with a slight deflation. However, for PMR-Egger, Otters, and HMAT, the type I error inflation is substantial (Figure 6) when using summary-level data.

**Figure 5:**
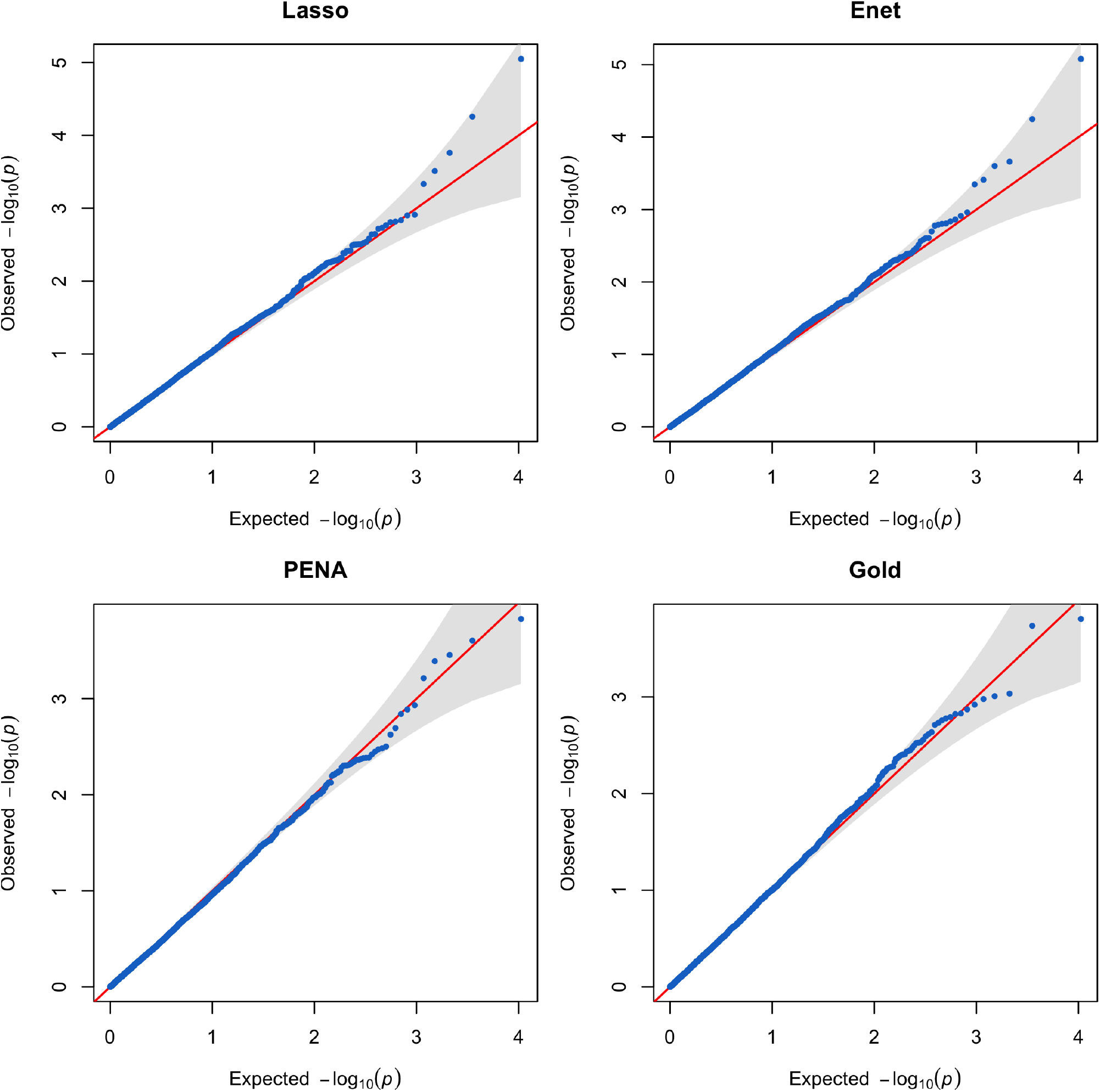
QQ plots for null p-values produced by the unadjusted TWAS approaches and the adjusted TWAS approach, PenaTWAS, based on summary-level data. Results for the gold standard TWAS, in which true eQTL effects are plugged in, are also presented. The unadjusted TWAS approaches are Lasso and Enet (elastic net) TWAS, depending on the penalty function used. The first choice of true eQTL effects with gene expression heritability of 10% was considered. 95% confidence band of the QQ plot is also presented.

**Figure 6:**
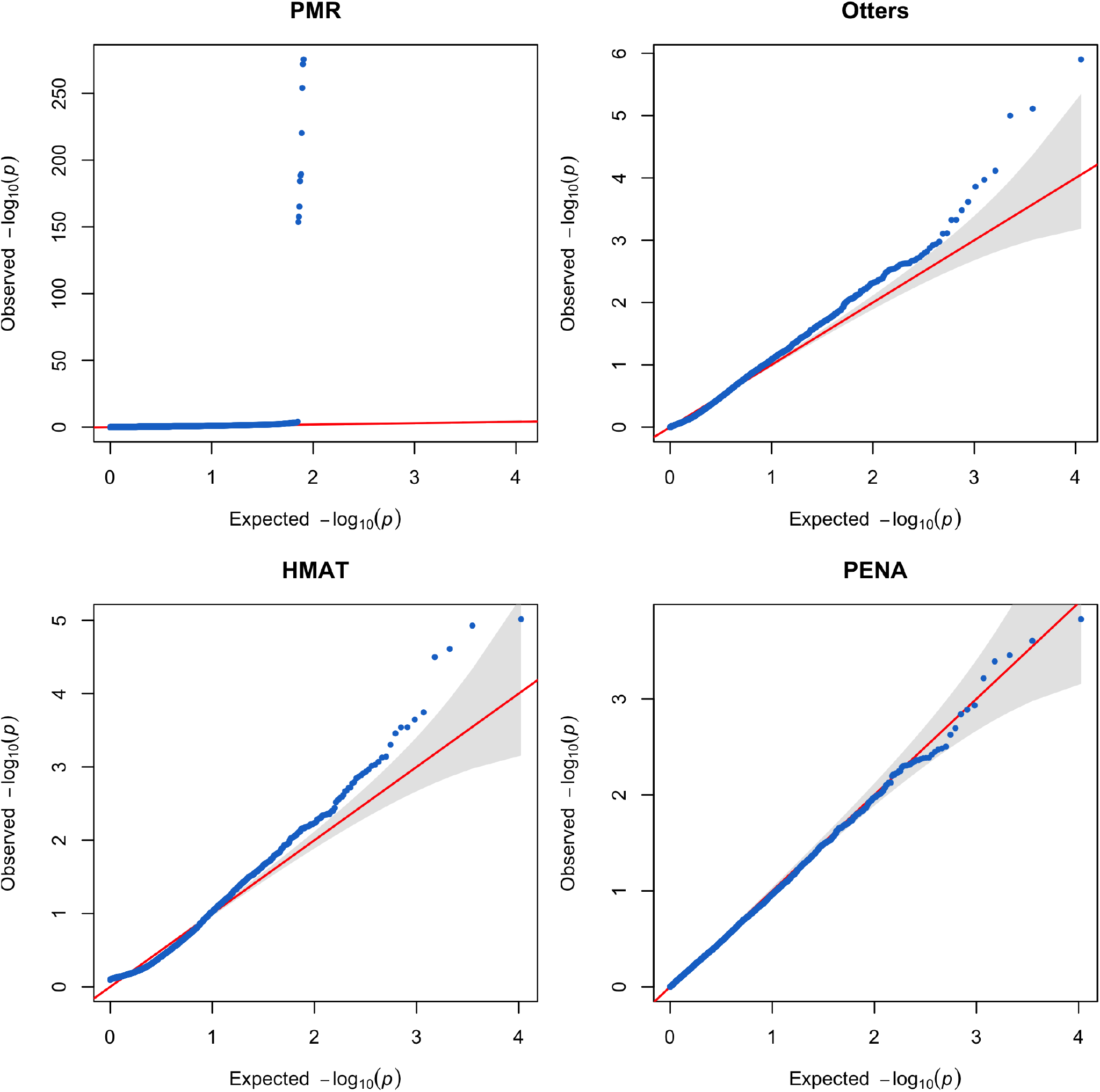
QQ plots for null p-values produced by the summary-level PMR-Egger, Otters, HMAT, and PenaTWAS. PenaTWAS and PMR-Egger calibrate the uncertainty of predicted expressions. HMAT and Otters ignore the uncertainty in predicted expressions. The heritability of gene expression is 10%. 95% confidence band of the QQ plot is also presented.

We find that the median of the adjustment factors to obtain the PenaTWAS test statistics from the unadjusted ALasso TWAS test statistics is 0.94, 0.95, and 0.94 in the three different simulation scenarios considered for a continuous outcome above (Figure S12). We note that the adjustment factors can sometimes be higher than one, which, however, happens less frequently, e.g., in 14%, 10%, and 2% of all iterations in the above three simulation scenarios. An adjustment factor marginally lower than one in most simulation iterations partly explains the better control of the type I error rates by PenaTWAS and the limited loss of power.

The empirical density plots of the individual-level PenaTWAS test statistics appear bell-shaped and symmetric around the mean zero under the null hypothesis (Figure S13). The Kolmogorov-Smirnov (KS) test for deviation from Normal distribution indicated an excellent match with normality, when the adjustment factor in PenaTWAS was calculated using 5% and 10% trimmed means of the bootstrap sample to estimate the denominator (Equation 10). The KS test p-values were 0.07 and 0.61, respectively. However, without trimming, the test statistics suggest deviation from normality with a p-value of 6.2 × 10^−5^ at a nominal level 0.05. We obtained a similar finding for summary-TWAS (Figure S14). The p-values from the KS test were 0.27, 0.34, and 0.01, for PenaTWAS10, PenaTWAS5, PenaTWAS0, respectively. Thus, PenaTWAS5 with 5% trimming appears robust, as its test statistics closely approximate *N* (0, 1) under the null. QQ plots demonstrate a perfect match between the null distribution and *N* (0, 1) (Figure S15, S16), more for 5% and 10% trimmed means.

## 4 Real data application

We applied the summary-level adjusted and unadjusted TWAS approaches to an anthropometric phenotype, height, and three lipid phenotypes: LDL cholesterol, HDL cholesterol, and triglycerides. We integrated the Geuvadis transcriptome data [24] and the UK Biobank (UKB) GWAS data [25]. We included the European individuals in Geuvadis and the white-British European individuals in the UKB data.

In Geuvadis data comprising 462 individuals from the 1000 Genomes Project, RNA sequencing was conducted for the lymphoblastoid cell line in peripheral blood tissue. Gene expression levels were measured for individuals who belonged to five different populations: CEPH (CEU), Finns (FIN), British (GBR), Toscani (TSI), and Yoruba (YRI). The individuals from CEU, FIN, GBR, and TSI were combined to form a sample from broader European (EUR) ancestry with a size of 358. Even though peripheral blood is one of the most studied tissue types, we acknowledge that it may not be the most relevant tissue type for the outcomes. For example, liver tissue, instead of blood tissue, is one of the most pertinent tissue types for lipids. We implemented the following QC steps for the genotype data. We filtered out SNPs with MAF *<* 0.05. We tested for the Hardy-Weinberg equilibrium (HWE) and removed the SNPs with p-value *<* 5 × 10^−8^.

We analyzed 14, 123 protein-coding genes. We adjusted the expression of each gene for relevant covariates, e.g., sex and the top 10 principal components (PCs) of genetic ancestry. The genetic PCs were included to adjust for the effect of population stratification on gene expression. For each gene, we regarded the SNPs located in one megabase region from the transcription start and end sites of the gene as the local SNPs. We implemented the GCTA software [23] to determine if a gene is locally heritable. Using GCTA, we obtained the p-value for each gene, testing the non-null hypothesis that its local heritability is significantly positive. We applied the BH FDR controlling procedure [26] to the p-values based on 5% FDR level to select the locally heritable genes. If a gene is locally heritable, we analyze it further. We found 1515 locally heritable genes; a limited number of genes were detected mainly due to the small sample size of the Geuvadis data. The same QC steps above were implemented to filter the genotype data in UKB. The MAF threshold was considered as 0.01 since the UKB data had an enormous sample size. After filtering for relatedness, 290,641 individuals were included in the UKB. We obtained the summary-level GWAS association data by adjusting for relevant covariates, such as age, sex, and the top 20 PCs for genetic ancestry. We employed the penalized regression based on the Elastic net penalty for the unadjusted TWAS, i.e., Enet TWAS. We utilized 300 residual bootstrap samples for PenaTWAS. We implemented another unadjusted summary-TWAS approach, Otters. PMR-Egger, using summary-level association data, yielded an error message on matrix decomposition and could not be implemented here. Analogous to the subset testing approach [27, 28], we apply the Bonferroni-corrected threshold for TWAS p-values in the second step regression for the 1515 genes selected in the first step, i.e., 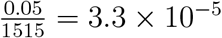.

First, we discuss the number of genes identified by PenaTWAS and Enet TWAS for the phenotypes. PenaTWAS discovered 120 genes for height, whereas Enet TWAS detected 142 genes. The two approaches identified a common subset of 102 genes (Table S17), leaving 18 genes discovered exclusively by PenaTWAS (Figure 7, Table S18) and 40 genes found exclusively by Enet TWAS (Figure 7, Table S18). PenaTWAS discovered 33 genes for HDL cholesterol, and Enet TWAS found 40 genes (Table S11, S12). Both approaches identified a common subset of 26 genes (Figure 7, Table S11). PenaTWAS spotted 25 genes for LDL cholesterol, and Enet TWAS found 26 genes. Both approaches detected a common set of 18 genes (Table S13, Figure 7). For triglycerides, PenaTWAS detected 32 genes, and Enet TWAS found 30 genes. Both methods identified a common set of 20 genes (Table S15), Figure 7). Thus, except for triglycerides, Enet TWAS found a few more genes than PenaTWAS, which resembles a limited higher power of unadjusted TWAS at the expense of inflated false positive rates observed in simulations. For each phenotype, nearly half or more of the discovered genes were identified by both approaches. Otters detected a few more genes than Enet TWAS for the phenotypes analyzed: 156 genes for height, 56 for HDL cholesterol, 38 genes for LDL cholesterol, and 40 for triglycerides. Some of these additional signals can be true; however, more false-positive findings can be present. Simulation studies showed a higher inflation of the type I error rate by Otters than by Enet TWAS and PenaTWAS. The latter and Otters identified 83 common genes for height, 17 overlapping genes for LDL, 20 common genes for HDL, and 17 overlapping genes for triglycerides, respectively.

**Figure 7:**
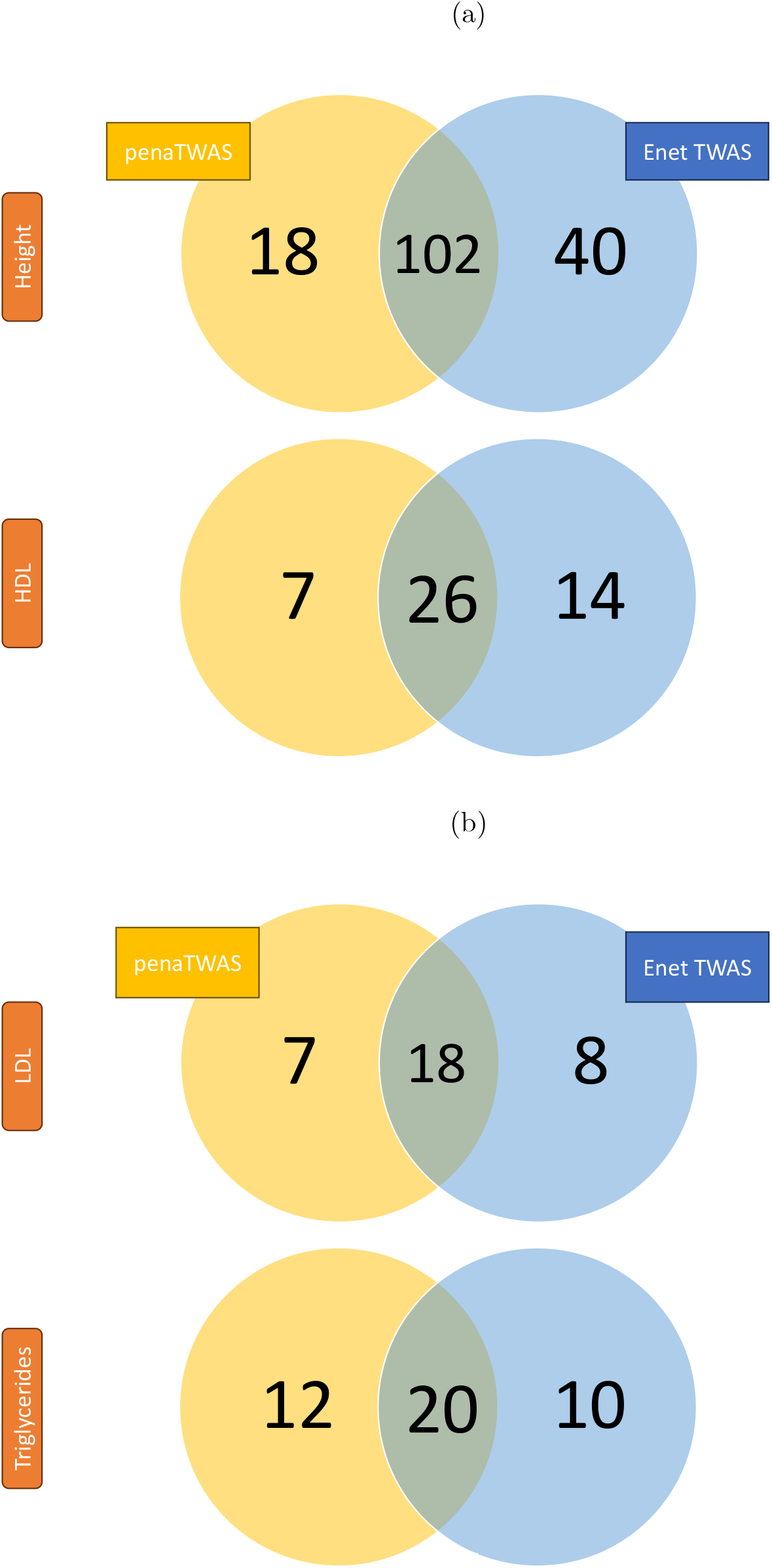
Number of genes associated with a phenotype identified by both PenaTWAS and Enet TWAS, only by PenaTWAS, and only by Enet TWAS. The subfigure (a) presents the results for height and HDL cholesterol, and the subfigure (b) presents the results for LDL cholesterol and triglycerides. PenaTWAS represents our adjusted TWAS approach, and Enet TWAS represents the unadjusted TWAS approach.

It is very challenging to demonstrate inflation in the false positive rate of the competing approaches based on real data analyses. For a heuristic exploration, we calculated the genomic inflation factor (GIF) based on the test statistics calculated by PenaTWAS and Enet TWAS for 1515 genes found locally heritable by GCTA. The GIF for PenaTWAS and Enet TWAS were 3.81 and 4 for height, 1.92 and 2.29 for HDL, 1.68 and 1.77 for LDL, 1.7 and 1.9 for triglycerides. The GIFs are substantially higher than one, mainly due to the estimation of these based on a limited number of locally heritable genes. Ideally, the GIF should be calculated based on all genes, which is at least ten-fold higher than the locally heritable genes in our analysis. However, relatively higher inflation for Enet TWAS than PenaTWAS is consistently observed across phenotypes. Interestingly, PenaTWAS found a few more genes associated with triglycerides despite a lower GIF than Enet TWAS. As expected, height appeared most polygenic. It is challenging to calculate the GIF for Otters as the method combines the p-values across different TWAS models using Cauchy aggregation.

Among the genes uniquely identified by Enet TWAS, many genes were missed by PenaTWAS with nominally significant association p-values. However, for some genes, PenaTWAS p-values are entirely insignificant. For height, the SLC25A16 gene located in the 10q21.3 region on chromosome 10 had a p-value of 2.6 × 10^−10^ by Enet TWAS, but a PenaTWAS p-value of 0.7 (Table S18). For HDL, the EEF2 gene located in the 19p13.3 region was detected by Enet TWAS but excluded by PenaTWAS with a p-value of 0.5 (Table S12). We observe similar patterns for LDL (Table S14) and triglycerides (Table S16). We assessed how many uniquely detected lipid phenotype genes are reported in the NHGRI-EBI GWAS catalog. For HDL, 5 out of 14 genes exclusively found by Enet TWAS and 1 out of 7 genes spotted by PenaTWAS are reported (Table S12). 4 out of 8 genes for Enet TWAS, and 2 out of 7 genes for PenaTWAS are reported for LDL (Table S14). For triglycerides, 2 out of 10 genes for Enet TWAS and 5 out of 12 genes for PenaTWAS are reported (Table S16). These exclusively identified genes that are not reported for lipids were communicated to be associated with many other biomedically important phenotypes in the catalog (Table S12, S14, S16). Interestingly, the gene TMEM120A on chromosome 7 was uniquely identified by PenaTWAS for all three lipid phenotypes, which suggests that it is a pleiotropic gene. Similarly, PenaTWAS detected the SLC25A16 gene on chromosome 10 to be significantly associated with height, LDL, and triglycerides; hence, it is likely to be a pleiotropic gene.

We performed pathway enrichment analysis for the PenaTWAS genes associated with lipids, implementing the *reactome* method [29]. We found the Glutathione conjugation pathway enriched for the HDL genes at 5% FDR level. The Glutathione conjugation pathway has been reported to regulate lipid levels in mice [30]. The ‘Rhesus blood group biosynthesis’ pathway was found for LDL genes. This pathway is not communicated to be relevant for LDL or other lipids. The rhesus blood group system is an important blood group system. For the triglycerides genes, *reactome* detected the ‘RUNX1 regulates transcription of genes involved in BCR signaling’ pathway to be enriched. The RUNX1 gene has been reported to be involved in the lipid metabolism of skin epithelial cells [31]. Thus, PenaTWAS performed well in discovering many gene-phenotype associations, including known and novel signals.

We also explored whether the genes discarded by PenaTWAS for lipid phenotypes are truly false positives. For this exploration, we took the genes associated with hypercholesterolemia from the OMIM database. The latter provides a list of biologically validated genes. These were APOA2, GHR, GSBS, EPHX2, and LDLR. However, we found that none of these genes were identified by any of the PenaTWAS, Enet TWAS, or Otters. One possible reason is that the TWAS methods applied to these real-data applications used whole-blood tissue-specific gene expression measurements in the Geuvadis transcriptome data rather than the directly relevant tissue type, e.g., liver tissue for lipid phenotypes.

We applied the Bonferroni corrected threshold using the number of locally heritable genes found by GCTA (see supplementary materials). However, another common practice is to employ a more stringent Bonferroni threshold using 20,000 as the total number of genes, i.e., 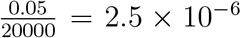. While the methods detected a slightly smaller number of genes based on this threshold (Table S10), the overall summary on the comparative performance of the different approaches remains similar to the above. Here, PenaTWAS and Otters detected 24 and 25 genes for triglycerides, respectively, higher than Enet TWAS, 18 genes (Table S10).

## 5 Discussion

TWAS-identified association signals need to be reliable so that follow-up functional studies can validate them successfully. An unadjusted TWAS approach suffers from the inherent methodological limitation that the uncertainty or variability of predicted expression is disregarded in the second step of regression of the outcome phenotype in GWAS data. The variance of the least squares estimator of the effect size for a standard TWAS is not calibrated for the uncertainty in predicted expression. Under the null hypothesis 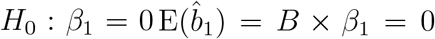. However, an unadjusted TWAS uses 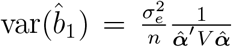. It ignores the expectation in the variance term calibrated for the uncertainty in the prediction model for expression: 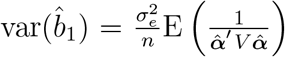. This shortcoming can lead to inference inaccuracies while identifying the gene-phenotype associations.

A standard TWAS is valid conditioned on the estimated local SNP effects on gene expression, hence the predicted expression. However, in the unconditioned form, we need to take into account the uncertainty of the estimated effects/predicted expression. Conditioning on the predicted expression, the expectation of the TWAS *Z* statistic is zero under the null, and the variance term is correctly defined. Unconditionally, i.e., while considering the uncertainty in the predicted expression, the expectation remains zero. However, the variance term does not remain the same as in standard TWAS (above). We can view the standard TWAS in two ways: 1) a burden-type test in GWAS data [32], where the relative weights of the SNPs to be aggregated come from training the transcriptome data in the form of cis-eQTL effects, and 2) evaluating the effect of predicted gene expression on the outcome phenotype in GWAS. The biological interpretation of the latter is more attractive, since we are not aggregating any set of SNPs but rather local SNPs surrounding a gene that explain a substantial proportion of the variation in gene expression. Xue et al. [33] show that the effect size estimator asymptotically converges to the actual effect size in TWAS. For a limited sample size of transcriptome data, asymptotic convergence may not hold well. The sample size may be smaller for several tissue types, e.g., brain tissue in the GTEx data.

We propose a novel approach adjusted for the uncertainty of predicted expression in TWAS. Adapting techniques from measurement error theory, we derive the adjustment factor. It is multiplied by the unadjusted TWAS test statistic obtained by using adaptive Lasso in transcriptome data. Then, we implement bootstrapping for adaptive Lasso to devise an estimation procedure for the adjustment factor. The proposed method, PenaTWAS, can be implemented for individual and summary-level GWAS data. Simulation studies show that PenaTWAS performs reliably in controlling the type I error rate for both continuous and binary traits. It consistently estimates the type I error rate slightly conservatively compared to the Gold TWAS. An unadjusted TWAS inflates the type I error rate, more so when using summary-level GWAS data. We also found simulation scenarios where an unadjusted TWAS adequately controls the type I error rate. However, for real-life data, assessing whether an unadjusted approach would maintain the rate of type I errors at a desired level is infeasible. On the contrary, PenaTWAS calibrates the uncertainty of predicted expression and is a more reliable alternative.

While estimating the adjustment factor in PenaTWAS, we considered three different choices of trimmed means: no trimming, 5% trimming, and 10% trimming of the bootstrap samples. More trimming would result in the exclusion of more bootstrap samples. We need an adequate number of final bootstrap samples to estimate the adjustment factor. Hence, more trimming will increase computing costs. Therefore, we restricted the trimming to a maximum of 10%. PenaTWAS0 is substantially more conservative. Both PenaTWAS5 and PenaTWAS10 demonstrate adequate control of type I errors and yield comparable powers. The null distribution of PenaTWAS5 test statistics fits *N* (0, 1) well with a Kolmogorov-Smirnov (KS) test p-value of 0.61. However, the KS test for evaluating the distributional fit of the PenaTWAS10 test statistics under the null hypothesis produced a p-value of 0.07, which is nearly significant at the nominal level of 0.05. Hence, the null distribution of the PenaTWAS10 test statistics may deviate marginally from *N* (0, 1). This is the primary empirical evidence from simulations, which led us to accept PenaTWAS5 as the preferred approach. In addition, summary-level PenaTWAS10 marginally inflated the type I error rate when using out-of-sample LD. A data-adaptive choice of the trimming percentage would be ideal; however, it requires novel strategies that we plan to explore in future studies.

In the simulation study, we allowed a gene in the downstream analysis if the GCTA p-value, for testing whether the local heritability of the gene’s expression is positive, passed the FDR correction. However, if the more stringent Bonferroni correction were applied instead to control FWER, the inflation of the type I error rates would be higher for an unadjusted TWAS [8]. The estimation of effect sizes is substantially more accurate for PenaTWAS than for an unadjusted TWAS. We observe it for both continuous and case-control phenotypes. The more precise estimation of effect sizes should improve the polygenic transcriptomic risk score (PTRS) calculation while predicting a quantitative or disease phenotype [34].

Leeuw et al. [8] argued that the TWAS framework tests for a non-zero local genetic correlation between the gene expression and the outcome. However, by ignoring the uncertainty of predicted expression, the null hypothesis in standard TWAS reduces to the local genetic correlation being constant, which need not be zero [8]. Following their derivation, it is straightforward to show that our null hypothesis *H*_0_ : *β*_1_ = 0 based on the ideal regression is equivalent to the correct null hypothesis of the local genetic correlation being equal to zero. We note that *β*_1_ is proportional to the covariance between the outcome and the true genetic component of expression based on the ideal regression (more details provided in the supplementary materials).

We modified an adjusted-variance approach [12], which also controls the false positive rate. For a lower choice of gene expression heritability, PenaTWAS yielded a substantially higher power than the adjusted variance approach, and the latter produced limited higher power for a higher choice of expression heritability. The formula for adjusted variance of the TWAS effect size estimator in the adjusted variance approach (provided in the supplementary material) shows that the corrected variance decreases with the heritability of gene expression. Because the error variance in the 1^*st*^ stage regression of gene expression decreases as expression heritability increases. As the standard error of the effect size estimate decreases, power should increase compared to settings with lower expression heritability. We note that many genes may not have a higher local heritability. The adjusted variance approach requires individual-level GWAS data.

CoMM and PMR-Egger adjust for uncertainty in expression prediction via joint modeling of the transcriptome and GWAS data. Simulations show that these approaches are not immune to the inflation of false positive rates. Both CoMM and PMR-Egger assume that all local SNPs around a gene have non-zero effects on the gene expression, i.e., all local SNPs are eQTLs for a gene. This assumption is debatable. The heritability of gene expression due to local SNPs is limited. Most eQTL studies discovered a small fraction of local SNPs as eQTLs. Many eQTLs play the role of various regulatory elements for gene expression, such as enhancers, promoters, etc. The phenomenon that all local SNPs have non-zero effects on expression will imply a large number of regulatory variants for a gene, which can be challenging to interpret biologically. However, the limited sample size of existing eQTL studies can also contribute to discovering fewer eQTLs, and the polygenicity of eQTL effects can be more extensive. PenaTWAS explicitly models sparsity in the eQTL effects. The key methodological difference between PenaTWAS and the above methods is that PenaTWAS considers adaptive Lasso while modeling the relationship between gene expression and the gene’s local SNPs. It allows for the shrinkage of minor effects and variable selection. The above approaches assume normality for the effect sizes of all local SNPs, which does not allow for explicit variable selection. For example, the model underlying PMR-Egger considers independent normal priors for the regression coefficients of the linear model in the transcriptome data, which is analogous to a penalty function considered in ridge regression for shrinkage. It does not perform differential shrinkage and variable selection like the adaptive Lasso.

We conducted summary-level TWAS for height and lipid phenotypes while integrating the Geuvadis and UK Biobank data. PenaTWAS performed well; the number of genes discovered was a little smaller than the unadjusted TWAS approach, Enet TWAS, for height, HDL, and LDL cholesterols, falling in line with the simulation results. However, the former detected a few more genes for triglycerides. PenaTWAS found some genes overlooked by the unadjusted TWAS and vice versa. However, for each phenotype, nearly half or more genes were detected by both. Many of the identified gene-phenotype associations have already been communicated in the literature. Evaluating the accuracy of the adjusted versus unadjusted approaches based on reported associations is challenging. Appropriate replication studies are crucial in this regard. PenaTWAS exclusively identified some genes for each phenotype, which the unadjusted TWAS missed. This can happen because our simulation studies demonstrated that the adjustment factor in PenaTWAS is not bounded above by one (Equation 10). Relatively few genes were identified exclusively by PenaTWAS. In simulations, the adjustment factor was more than unity for a limited percentage of all simulation iterations. Another unadjusted approach, Otters, identified more genes for the phenotypes. However, it inflated the false positive rate more in simulations.

While comparing with other adjusted approaches for TWAS based on individual-level data, we observe that PMR-Egger also controls the type I error rate adequately. It produces a limited power gain compared to PenaTWAS. But for the summary-level TWAS, PMR-Egger inflates the type I error rate. Hence, we recommend PenaTWAS to perform summary-level TWAS when a study aims to identify reliable gene-trait associations with fewer false positives. However, if a summary-level TWAS aims to discover more associated genes and the inflation of the false positive rate to some extent is acceptable, we recommend PMR-Egger and Otters instead.

The elastic net penalty is more suited for covariates with correlation, e.g., SNPs in LD. However, our extensive simulation study finds that the adaptive Lasso TWAS performs robustly even in the presence of LD among the local SNPs. The unadjusted TWAS based on adaptive Lasso leads to a limited power loss compared to the unadjusted TWAS based on Elastic net and Lasso. Adjusting for the uncertainty of imputed expression based on the adaptive Lasso reduces the power a little more, i.e., PenaTWAS loses a limited power compared to ALasso TWAS. We explored the residual bootstrap for the elastic net, but found that the type I error rate estimation is over-conservative and the power loss is severe (around 40-50%). The residual bootstrap estimates for the elastic net may not converge in distribution.

We acknowledge that there are other novel TWAS methods in the literature that we did not compare PenaTWAS to. For example, one such method is PTWAS [35], which analyzes causality using a novel strategy after an initial scan to detect marginally associated genes. An important future direction of work will be to develop computationally feasible bootstrapping algorithms for elastic net and Lasso regressions. In future work, we aim to apply the method to more phenotypes while integrating transcriptome data with larger sample sizes in various tissues or cell types. We plan to extend this approach to incorporate multiple ancestries. An important direction is upgrading the method to combine multiple tissue types available in transcriptome data. We explored the utility of PenaTWAS for a case-control outcome, even though we developed the method for a continuous outcome. We plan to explicitly extend the approach for a case-control phenotype.

In summary, we propose a TWAS approach that calibrates the predicted expression’s uncertainty and is a statistically valid test for gene-phenotype association. The method is technically sound. It can be implemented for complex phenotypes using publicly available GWAS summary statistics data to obtain novel insights into their genetic architecture.

## 6 Data availability

The datasets that we have analyzed or used in our analyses in this work are available (either openly or via applications) from the following websites:

Geuvadis data: https://www.internationalgenome.org/data-portal/data-collection/geuvadis.

UK Biobank: https://www.ukbiobank.ac.uk/.

Fusion: http://gusevlab.org/projects/fusion/

## 7 Acknowledgement

The authors thank Dr. Arindam Chatterjee and Dr. Prajamitra Bhuyan for helpful discussions related to this work. This research used the UK Biobank resource under application 77327. Arunabha Majumdar was supported by the Anusandhan National Research Foundation (ANRF) advanced research grant with an identification number ANRF/ARG/2025/005607/MS.

