## Supplementary text, tables and figures for "A TWAS method calibrated for the uncertainty of predicted expression"

#### 1 Introduction: an example of TWAS

Consider the lipid phenotype, HDL cholesterol, the outcome of interest, and liver tissue as the relevant tissue type. As an example of the reference transcriptome data, the GTEx data [36] has measurements of gene expression in various tissue types and genotypes for genome-wide SNPs. In the first step of TWAS, we consider a gene and its expression in the liver tissue and a set of local SNPs surrounding the gene, for example, within one megabase region from the gene's transcription start and end sites. We fit a regression model of expression on the local SNPs' genotypes in the GTEx data to learn a prediction model for the expression. In the second step, we have a separate GWAS dataset. For example, the UK Biobank has data for HDL and genotypes for genome-wide SNPs, but expression data is unavailable. We implement the prediction model of expression obtained in the GTEx data to predict the expression for the GWAS individuals in the UK Biobank. Finally, we regress HDL on the predicted genetically regulated component of expression to test for an association between HDL and the gene.

#### 2 Materials and methods

We use the following main notations to describe the details of our method.

#### 2.1 Various notations used for the transcriptome data

$n_1$: number of individuals.

$\mathbf{E}$: vector of expression values for a gene for $n_1$ individuals in the specific tissue of interest.

$E_i$: expression for $i^{th}$ individual, $i = 1, \ldots, n_1$.

$p$: number of local SNPs for the gene.

$G_{\text{ref}}$: $n_1 \times p$ genotype matrix at local SNPs for individuals in reference transcriptome data.

$\mathbf{g}_{\text{ref},i}$: genotype vector at local SNPs for $i^{th}$ individual in transcriptome data, $i = 1, \ldots, n_1$.

$\boldsymbol{\alpha}$: effect size vector for $p$ local SNPs on the expression of the gene.

#### 2.2 Various notations used for the GWAS data

$n$: number of individuals.

$\mathbf{y}$: vector of outcome phenotype values for $n$ individuals in GWAS.

$G$: genotype matrix of order $n \times p$ for GWAS individuals at $p$ local SNPs.

$x_i$: true genetic component of expression for $i^{th}$ individual in GWAS, $i = 1, \ldots, n$.

$X_i$: predicted genetic component of expression for $i^{th}$ individual in GWAS, $i = 1, \ldots, n$.

$\beta_1$: effect of the actual genetic component of expression on the outcome in GWAS.

$b_1$: effect of the predicted genetic component of expression on the outcome.

$\mathbf{z}_{\text{gwas}}$: vector of $Z$ (Wald) statistics for marginal SNP-phenotype association at the $p$ local SNPs in GWAS.

$Z_{\text{unadj}}$: $Z$-statistic for the unadjusted TWAS.

$Z_{\text{pena}}$: $Z$-statistic for PenaTWAS.

$V$: linkage disequilibrium (LD) matrix for the $p$ local SNPs in GWAS.

#### 2.3 Regression of gene expression on local SNPs

The main model for the regression of gene expression on the genotypes of local SNPs in reference transcriptome data has been described in the main text (Equation 1). Since the number of local SNPs can be larger than the sample size of the reference data, a penalized regression is implemented in practice to estimate $\boldsymbol{\alpha}$, e.g., Lasso [19] and Elastic net [20] are two commonly used methods.

#### 2.4 Regression of phenotype on genetic component of expression

##### 2.4.1 Ideal and working regressions

The definitions of the ideal and working regressions are provided in the main text (Equations 5 and 6). Based on the ideal regression, $\mathrm{E}(y_i) = \beta_0 + \beta_1 x_i$ and $\mathrm{var}(y_i) = \mathrm{var}(\epsilon_i) = \sigma_\epsilon^2$. Since $\boldsymbol{\alpha}$ is an unknown constant vector, $x_i = \mathbf{g}_i^{'}\boldsymbol{\alpha}$ is non-random.

##### 2.4.2 An adjusted estimator and related test

Our main goal is to obtain an estimator of $\beta_1$ in the ideal regression and test for $H_0 : \beta_1 = 0$ vs. $H_1 : \beta_1 \neq 0$. We obtain an estimator of $\beta_1$ based on $\hat{b}_1$:

$$\hat{b}_1 = \frac{S_{Xy}}{S_{XX}} = \frac{\sum_{i=1}^{n}(X_i - \bar{X})(y_i - \bar{y})}{\sum_{j=1}^{n}(X_j - \bar{X})^2}. \tag{15}$$

$\hat{b}_1$ is the LSE of $b_1$, hence it is an unbiased estimator of $b_1$ based on the working regression (Equation 6). We note that:

$$\sum_{i=1}^{n} X_i = \sum_{i=1}^{n} \mathbf{g}_i^{'}\hat{\boldsymbol{\alpha}} = (\sum_{i=1}^{n} \mathbf{g}_i)^{'}\hat{\boldsymbol{\alpha}} = 0, \text{ since } \sum_{i=1}^{n} \mathbf{g}_i = \mathbf{0}. \tag{16}$$

Because each local SNP's genotype data is normalized to have a mean of zero and a variance of one. Thus, $\bar{X} = 0$. The randomness of $X_i$ is due to $\hat{\boldsymbol{\alpha}}$, and $y_i$ is random in the ideal regression due to $\epsilon_i$, $i = 1, \ldots, n$. Denote $\boldsymbol{\epsilon} = (\epsilon_1, \ldots, \epsilon_n)'$. We obtain that:

$$\mathrm{E}(\hat{b}_1) = \mathrm{E}_{\hat{\boldsymbol{\alpha}}}\mathrm{E}_{\boldsymbol{\epsilon}|\hat{\boldsymbol{\alpha}}}(\hat{b}_1) = \mathrm{E}_{\hat{\boldsymbol{\alpha}}}\mathrm{E}_{\boldsymbol{\epsilon}|\hat{\boldsymbol{\alpha}}}\left(\frac{\sum_{i=1}^n X_i(y_i - \bar{y})}{\sum_{j=1}^n X_j^2}\right) = \mathrm{E}_{\hat{\boldsymbol{\alpha}}}\left(\sum_{i=1}^n \frac{X_i}{\sum_{j=1}^n X_j^2}\mathrm{E}_{\boldsymbol{\epsilon}|\hat{\boldsymbol{\alpha}}}(y_i - \bar{y})\right). \tag{17}$$

Since $\boldsymbol{\epsilon}$ in the ideal regression, based on the GWAS data, does not depend on $\hat{\boldsymbol{\alpha}}$, which is based on the transcriptome data, the conditional expectation $\mathrm{E}_{\boldsymbol{\epsilon}|\hat{\boldsymbol{\alpha}}}$ is the same as the marginal expectation $\mathrm{E}_{\boldsymbol{\epsilon}}$. Now, based on the ideal regression in Equation 5,

$$\mathrm{E}_{\boldsymbol{\epsilon}}(y_i) = \beta_0 + \beta_1 x_i,\ \mathrm{E}_{\boldsymbol{\epsilon}}(\bar{y}) = \frac{1}{n}\sum_{i=1}^n E(y_i) = \frac{1}{n}\sum_{i=1}^n(\beta_0 + \beta_1 x_i) = \beta_0 + \frac{\beta_1}{n}\sum_{i=1}^n x_i. \tag{18}$$

Because the genotype vector is centered at the mean, we note that:

$$\sum_{i=1}^n \mathbf{g}_i = \mathbf{0}, \sum_{i=1}^n x_i = \sum_{i=1}^n \mathbf{g}_i^{'}\boldsymbol{\alpha} = (\sum_{i=1}^n \mathbf{g}_i)^{'}\boldsymbol{\alpha} = 0, E_{\boldsymbol{\epsilon}}(y_i - \bar{y}) = \beta_1 x_i. \tag{19}$$

Then we obtain:

$$\sum_{j=1}^n X_j^2 = \sum_{j=1}^n X_j^{'}X_j = \sum_{j=1}^n \hat{\boldsymbol{\alpha}}^{'}\mathbf{g}_j\mathbf{g}_j^{'}\hat{\boldsymbol{\alpha}} = \hat{\boldsymbol{\alpha}}^{'}\left(\sum_{j=1}^n \mathbf{g}_j\mathbf{g}_j^{'}\right)\hat{\boldsymbol{\alpha}} = n\hat{\boldsymbol{\alpha}}^{'}V\hat{\boldsymbol{\alpha}}, V = \frac{1}{n}\sum_{j=1}^n \mathbf{g}_j\mathbf{g}_j^{'}.$$

Here, $V$ denotes the linkage disequilibrium (LD) matrix of the $p$ local SNPs based on the GWAS data. Thus,

$$\begin{aligned}\mathrm{E}(\hat{b}_1) &= \mathrm{E}_{\hat{\boldsymbol{\alpha}}}\left(\sum_{i=1}^n \frac{X_i}{n\hat{\boldsymbol{\alpha}}'V\hat{\boldsymbol{\alpha}}}\beta_1 x_i\right) = \frac{\beta_1}{n}\mathrm{E}_{\hat{\boldsymbol{\alpha}}}\left(\sum_{i=1}^n x_i\frac{\mathbf{g}_i^{'}\hat{\boldsymbol{\alpha}}}{n\hat{\boldsymbol{\alpha}}'V\hat{\boldsymbol{\alpha}}}\right) = \frac{\beta_1}{n}\sum_{i=1}^n x_i\mathbf{g}_i^{'}\mathrm{E}_{\hat{\boldsymbol{\alpha}}}\left(\frac{\hat{\boldsymbol{\alpha}}}{\hat{\boldsymbol{\alpha}}'V\hat{\boldsymbol{\alpha}}}\right)\\ &= \frac{\beta_1}{n}\sum_{i=1}^n(\boldsymbol{\alpha}^{'}\mathbf{g}_i)\mathbf{g}_i^{'}\mathrm{E}\left(\frac{\hat{\boldsymbol{\alpha}}}{\hat{\boldsymbol{\alpha}}'V\hat{\boldsymbol{\alpha}}}\right) = \beta_1\boldsymbol{\alpha}^{'}(\frac{1}{n}\sum_{i=1}^n \mathbf{g}_i\mathbf{g}_i^{'})\mathrm{E}\left(\frac{\hat{\boldsymbol{\alpha}}}{\hat{\boldsymbol{\alpha}}'V\hat{\boldsymbol{\alpha}}}\right) = \beta_1\boldsymbol{\alpha}^{'}V\mathrm{E}\left(\frac{\hat{\boldsymbol{\alpha}}}{\hat{\boldsymbol{\alpha}}'V\hat{\boldsymbol{\alpha}}}\right).\end{aligned} \tag{20}$$

The term multiplied by $\beta_1$ is an unknown constant. Therefore,

$$\mathrm{E}(\hat{b}_1) = B \times \beta_1, \text{ where } B = \boldsymbol{\alpha}' V \mathrm{E}\left(\frac{\hat{\boldsymbol{\alpha}}}{\hat{\boldsymbol{\alpha}}' V \hat{\boldsymbol{\alpha}}}\right). \text{ Hence, } \mathrm{E}\left(\frac{\hat{b}_1}{B}\right) = \beta_1. \tag{21}$$

We regard $\frac{\hat{b}_1}{\hat{B}}$ as an estimator of $\beta_1$, where $\hat{b}_1$ is an LSE of $b_1$ in the working regression. Next, we obtain the variance of $\hat{b}_1$. Since $X_i$ is $\mathbf{g}_i' \hat{\boldsymbol{\alpha}}$, the randomness of $X_i$ comes through $\hat{\boldsymbol{\alpha}}$. Hence, the $\mathbf{X}$ vector is fixed for a given $\hat{\boldsymbol{\alpha}}$. By the law of total variance,

$$\mathrm{var}(\hat{b}_1) = \mathrm{E}(\mathrm{var}(\hat{b}_1|\mathbf{X})) + \mathrm{var}(\mathrm{E}(\hat{b}_1|\mathbf{X})), \text{ where } \hat{b}_1 = \frac{\sum_{i=1}^n (X_i - \bar{X})(y_i - \bar{y})}{\sum_{j=1}^n (X_j - \bar{X})^2}. \tag{22}$$

$$\begin{aligned}
\mathrm{var}(\hat{b}_1|\mathbf{X}) &= \frac{1}{\{\sum_{j=1}^n (X_j - \bar{X})^2\}^2} \mathrm{var}\{\sum_{i=1}^n (X_i - \bar{X})(y_i - \bar{y})|\mathbf{X}\} \\
&= \frac{1}{\{\sum_{j=1}^n (X_j - \bar{X})^2\}^2} \mathrm{var}\{\sum_{i=1}^n (X_i - \bar{X}) y_i|\mathbf{X}\} \text{ [since } \sum_{i=1}^n (X_i - \bar{X})|\mathbf{X} = 0] \\
&= \frac{1}{\{\sum_{j=1}^n (X_j - \bar{X})^2\}^2} \sum_{i=1}^n (X_i - \bar{X})^2 \mathrm{var}(y_i|X_i) \text{ [since covariances are zero]} \\
&= \frac{\sigma_e^2}{\sum_{j=1}^n (X_j - \bar{X})^2} = \frac{\sigma_e^2}{\sum_{j=1}^n X_j^2} \text{ [since } \bar{X} = 0, \text{ conditioned on } \mathbf{X}] \\
&= \frac{\sigma_e^2}{n \hat{\boldsymbol{\alpha}}' V \hat{\boldsymbol{\alpha}}} \\
&\implies \mathrm{E}(\mathrm{var}(\hat{b}_1|\mathbf{X})) = \frac{\sigma_e^2}{n} \mathrm{E}\left(\frac{1}{\hat{\boldsymbol{\alpha}}' V \hat{\boldsymbol{\alpha}}}\right).
\end{aligned} \tag{23}$$

Based on the working regression model (Equation 6), $y_i|X_i$, $i = 1, \ldots, n$, are independently distributed with $\mathrm{var}(y_i|X_i) = \sigma_e^2$; hence, the covariance terms in the above variance become zero. The final expectation term concerns $\hat{\boldsymbol{\alpha}}$ based on the transcriptome data. The other part of $\mathrm{var}(\hat{b}_1)$ is $\mathrm{var}(\mathrm{E}(\hat{b}_1|\mathbf{X}))$.

$$\bar{X}|\mathbf{X}=0 \implies \mathrm{E}(\hat{b}_1|\mathbf{X})=\mathrm{E}\left(\frac{\sum_{i=1}^{n}X_i(y_i-\bar{y})}{\sum_{j=1}^{n}X_j^2}\middle|\mathbf{X}\right)=\sum_{i=1}^{n}\frac{X_i}{\sum_{j=1}^{n}X_j^2}\mathrm{E}\left((y_i-\bar{y})|\mathbf{X}\right),$$

$$\mathrm{E}(y_i|X_i)=b_0+b_1X_i, \mathrm{E}(\bar{y}|\mathbf{X})=b_0, \mathrm{E}\left((y_i-\bar{y})|\mathbf{X}\right)=b_1X_i \text{ [based on Equation (6)]}$$

$$\implies \mathrm{E}(\hat{b}_1|\mathbf{X})=b_1 \implies \mathrm{var}(\mathrm{E}(\hat{b}_1|\mathbf{X}))=0.$$

Conditioned on $\mathbf{X}$, $\hat{b}_1$ is an unbiased estimator of $b_1$ based on the working regression (Equation 6). Thus, putting together all parts of the derivations:

$$\mathrm{var}(\hat{b}_1)=\frac{\sigma_e^2}{n}\mathrm{E}\left(\frac{1}{\hat{\boldsymbol{\alpha}}'V\hat{\boldsymbol{\alpha}}}\right). \quad (24)$$

If $B$ was known in the Equation 21, an unbiased estimator of $\beta_1$ would be $\frac{\hat{b}_1}{B}$. However, $B$ is unknown. To test for $H_0:\beta_1=0$ versus $H_1:\beta_1\neq 0$, denote the $Z$ (Wald) statistic in PenaTWAS as $Z_{\text{pena}}$. If we have $\hat{B},\hat{\sigma}_e$, and $\hat{\mathrm{E}}\left(\frac{1}{\hat{\boldsymbol{\alpha}}'V\hat{\boldsymbol{\alpha}}}\right)$ as the estimates of $B,\sigma_e$, and $\mathrm{E}\left(\frac{1}{\hat{\boldsymbol{\alpha}}'V\hat{\boldsymbol{\alpha}}}\right)$, respectively,

$$Z_{\text{pena}}=\frac{\frac{\hat{b}_1}{\hat{B}}}{\text{s.e.}\left(\frac{\hat{b}_1}{\hat{B}}\right)}=\frac{\frac{\hat{b}_1}{\hat{B}}}{\frac{\hat{\sigma}_e}{\sqrt{n}\hat{B}}\sqrt{\hat{\mathrm{E}}\left(\frac{1}{\hat{\boldsymbol{\alpha}}'V\hat{\boldsymbol{\alpha}}}\right)}}=\frac{\sqrt{n}\hat{b}_1}{\hat{\sigma}_e\sqrt{\hat{\mathrm{E}}\left(\frac{1}{\hat{\boldsymbol{\alpha}}'V\hat{\boldsymbol{\alpha}}}\right)}}.$$

The $Z$ statistic for the unadjusted TWAS based on the working regression model (Equation 6) is obtained as:

$$Z_{\text{unadj}}=\frac{\hat{b}_1}{\text{s.e.}(\hat{b}_1)}=\frac{\sqrt{\hat{\boldsymbol{\alpha}}'V\hat{\boldsymbol{\alpha}}}\sqrt{n}\hat{b}_1}{\hat{\sigma}_e}, \text{ where } \hat{b}_1=\frac{\sum_{i=1}^{n}X_i(y_i-\bar{y})}{\sum_{j=1}^{n}X_j^2} \text{ and } \mathrm{var}(\hat{b}_1|\mathbf{X})=\frac{\sigma_e^2}{n\hat{\boldsymbol{\alpha}}'V\hat{\boldsymbol{\alpha}}}.$$

Thus, we obtain the relation between $Z_{\text{pena}}$ and $Z_{\text{unadj}}$ as:

$$Z_{\text{pena}} = \hat{A} \times Z_{\text{unadj}}, \text{ where } A = \frac{\frac{1}{\sqrt{\hat{\boldsymbol{\alpha}}' V \hat{\boldsymbol{\alpha}}}}}{\sqrt{\mathrm{E}\left(\frac{1}{\hat{\boldsymbol{\alpha}}' V \hat{\boldsymbol{\alpha}}}\right)}}. \tag{25}$$

To adjust for the uncertainty of the predicted expression, we need to multiply the $Z$ statistic used in the unadjusted TWAS by an estimate of the adjustment factor $A$. In the following, we discuss an approach to estimating the adjustment factor $A$. To obtain an estimate of $\beta_1$, we use Equation 21:

$$\hat{\beta}_1 = \frac{\hat{b}_1}{\hat{B}}, \text{ where } B = \boldsymbol{\alpha}' V \mathrm{E}\left(\frac{\hat{\boldsymbol{\alpha}}}{\hat{\boldsymbol{\alpha}}' V \hat{\boldsymbol{\alpha}}}\right) \text{ and } \hat{B} = \hat{\boldsymbol{\alpha}}' V \hat{\mathrm{E}}\left(\frac{\hat{\boldsymbol{\alpha}}}{\hat{\boldsymbol{\alpha}}' V \hat{\boldsymbol{\alpha}}}\right). \tag{26}$$

We plug in the estimate of $\boldsymbol{\alpha}$ based on complete data, $\hat{\boldsymbol{\alpha}}$. $\hat{\mathrm{E}}\left(\frac{\hat{\boldsymbol{\alpha}}}{\hat{\boldsymbol{\alpha}}' V \hat{\boldsymbol{\alpha}}}\right)$ is an estimate of $\mathrm{E}\left(\frac{\hat{\boldsymbol{\alpha}}}{\hat{\boldsymbol{\alpha}}' V \hat{\boldsymbol{\alpha}}}\right)$.

#### 2.5 Taylor's expanded version of PenaTWAS: PENAtaylor

We devise another approach to estimate $\mathrm{E}\left(\frac{1}{\hat{\boldsymbol{\alpha}}' V \hat{\boldsymbol{\alpha}}}\right)$ based on the bootstrap sample. Using Taylor's theorem, we can approximate the following multi-variable function:

$$f(\hat{\boldsymbol{\alpha}}) = \frac{1}{\hat{\boldsymbol{\alpha}}' V \hat{\boldsymbol{\alpha}}}, f(\hat{\boldsymbol{\alpha}}) \approx f(\hat{\boldsymbol{\alpha}}_{\text{obs}}) + (\hat{\boldsymbol{\alpha}} - \hat{\boldsymbol{\alpha}}_{\text{obs}})' Df(\hat{\boldsymbol{\alpha}}_{\text{obs}}) + \frac{1}{2}(\hat{\boldsymbol{\alpha}} - \hat{\boldsymbol{\alpha}}_{\text{obs}})' Hf(\hat{\boldsymbol{\alpha}}_{\text{obs}})(\hat{\boldsymbol{\alpha}} - \hat{\boldsymbol{\alpha}}_{\text{obs}}).$$

Here, $Df(\hat{\boldsymbol{\alpha}}_{\text{obs}})$ and $Hf(\hat{\boldsymbol{\alpha}}_{\text{obs}})$, i.e., $D^2 f(\hat{\boldsymbol{\alpha}}_{\text{obs}})$, denote the gradient vector and the Hessian matrix of $f$ evaluated at $\hat{\boldsymbol{\alpha}}_{\text{obs}}$, which is the original estimate of $\boldsymbol{\alpha}$ obtained by the adaptive

Lasso based on the complete data. Taking an expectation of both sides, we get:

$$\mathrm{E}(f(\hat{\boldsymbol{\alpha}})) \approx f(\hat{\boldsymbol{\alpha}}_{\text{obs}}) + \{\mathrm{E}(\hat{\boldsymbol{\alpha}} - \hat{\boldsymbol{\alpha}}_{\text{obs}})\}' Df(\hat{\boldsymbol{\alpha}}_{\text{obs}}) + \frac{1}{2}\mathrm{E}\{(\hat{\boldsymbol{\alpha}} - \hat{\boldsymbol{\alpha}}_{\text{obs}})' Hf(\hat{\boldsymbol{\alpha}}_{\text{obs}})(\hat{\boldsymbol{\alpha}} - \hat{\boldsymbol{\alpha}}_{\text{obs}})\},$$
$$Df(\hat{\boldsymbol{\alpha}}) = -\frac{2}{(\hat{\boldsymbol{\alpha}}' V \hat{\boldsymbol{\alpha}})^2} V\hat{\boldsymbol{\alpha}},\ Hf(\hat{\boldsymbol{\alpha}}) = -\frac{2}{(\hat{\boldsymbol{\alpha}}' V \hat{\boldsymbol{\alpha}})^2} V + \frac{8}{(\hat{\boldsymbol{\alpha}}' V \hat{\boldsymbol{\alpha}})^3} V\hat{\boldsymbol{\alpha}}\hat{\boldsymbol{\alpha}}' V.$$

Both $Df(\hat{\boldsymbol{\alpha}})$ and $Hf(\hat{\boldsymbol{\alpha}})$ are evaluated at $\hat{\boldsymbol{\alpha}} = \hat{\boldsymbol{\alpha}}_{\text{obs}}$. We use the bootstrapped estimates of $\boldsymbol{\alpha}$ to estimate the bias term and the expectation of the quadratic form as:

$$\hat{\mathrm{E}}(\hat{\boldsymbol{\alpha}} - \hat{\boldsymbol{\alpha}}_{\text{obs}}) = \frac{1}{T}\sum_{j=1}^{T}(\tilde{\boldsymbol{\alpha}}_j - \hat{\boldsymbol{\alpha}}_{\text{obs}}),\ \text{where } \tilde{\boldsymbol{\alpha}}_j \text{ is } j^{\text{th}} \text{ bootstrap sample of } \boldsymbol{\alpha}, j = 1, \ldots, T,$$

$$\hat{\mathrm{E}}\{(\hat{\boldsymbol{\alpha}} - \hat{\boldsymbol{\alpha}}_{\text{obs}})' Hf(\hat{\boldsymbol{\alpha}}_{\text{obs}})(\hat{\boldsymbol{\alpha}} - \hat{\boldsymbol{\alpha}}_{\text{obs}})\} = \frac{1}{T}\sum_{j=1}^{T}\{(\tilde{\boldsymbol{\alpha}}_j - \hat{\boldsymbol{\alpha}}_{\text{obs}})' Hf(\hat{\boldsymbol{\alpha}}_{\text{obs}})(\tilde{\boldsymbol{\alpha}}_j - \hat{\boldsymbol{\alpha}}_{\text{obs}})\}.$$

##### 2.5.1 Estimation of the TWAS effect size

For estimating the effect size $\beta_1$, we require an estimate of $B$ (Equation 26) which includes $\mathrm{E}\left(\frac{\hat{\boldsymbol{\alpha}}'}{\hat{\boldsymbol{\alpha}}' V \hat{\boldsymbol{\alpha}}}\right)$. We estimate this term based on the bootstrap sample as $\frac{1}{T}\sum_{j=1}^{T}\frac{1}{\tilde{\boldsymbol{\alpha}}_j' V \tilde{\boldsymbol{\alpha}}_j}\tilde{\boldsymbol{\alpha}}_j$. $B$ also involves $\boldsymbol{\alpha}$, which we replace by $\hat{\boldsymbol{\alpha}}$, the ALasso estimate based on original complete data. The ALasso estimator $\hat{\boldsymbol{\alpha}}$ is a consistent estimator of $\boldsymbol{\alpha}$. Altogether, $\hat{\beta}_1$, i.e., $\frac{\hat{b}_1}{\hat{B}}$ is a sensible estimator of $\beta_1$ (Equation 26).

#### 2.6 Null distribution of $Z_{\text{pena}}$

To test for $H_0 : \beta_1 = 0$ vs $H_1 : \beta_1 \neq 0$ in the ideal regression (Equation 5), we require the expectation and variance of $Z_{\text{pena}}$ under $H_0$.

##### 2.6.1 Expectation and variance of $Z_{\text{pena}}$ under null hypothesis

To test for $H_0 : \beta_1 = 0$ vs $H_1 : \beta_1 \neq 0$, we require the expectation and variance of $Z_{\text{pena}}$ under $H_0$. For the summary-TWAS, $Z_{\text{pena}} = \hat{A} \times Z_{\text{unadj}}$, where $A = \frac{\frac{1}{\sqrt{\hat{\boldsymbol{\alpha}}' V \hat{\boldsymbol{\alpha}}}}}{\sqrt{\mathrm{E}\left(\frac{1}{\hat{\boldsymbol{\alpha}}' V \hat{\boldsymbol{\alpha}}}\right)}}$ and $Z_{\text{unadj}} = \frac{\hat{\boldsymbol{\alpha}}' \mathbf{z}_{\text{gwas}}}{\sqrt{\hat{\boldsymbol{\alpha}}' V \hat{\boldsymbol{\alpha}}}}$. Since the transcriptome and GWAS data have separate groups of unrelated individuals, $\hat{\boldsymbol{\alpha}}$ and $\mathbf{z}_{\text{gwas}}$ are independent. Also, $V$ is a fixed LD matrix based on the GWAS or external data. Thus, $\mathrm{E}_{H_0}(Z_{\text{pena}}) = \mathrm{E}(\frac{\hat{A}}{\sqrt{\hat{\boldsymbol{\alpha}}' V \hat{\boldsymbol{\alpha}}}} \hat{\boldsymbol{\alpha}}')\mathrm{E}_{H_0}(\mathbf{z}_{\text{gwas}})$. Since the genotypes for each SNP and the phenotype data are normalized to have mean zero and variance one, $\mathbf{z}_{\text{gwas}}$ can be approximated as $\frac{1}{\sqrt{n}} G' \mathbf{y}$. Here, it is also assumed that the heritability of $Y$ due to a single SNP is negligible. Under the ideal regression model (Equation 5), $\mathrm{E}(y_i) = \beta_1 x_i$; $\beta_0 = 0$, because $y$s are mean-centered. Thus, $\mathrm{E}_{H_0}(\mathbf{z}_{\text{gwas}}) = \frac{\beta_1}{\sqrt{n}} G' \mathbf{x} = \mathbf{0}$, since $\beta_1 = 0$ under $H_0$. Hence, $\mathrm{E}_{H_0}(Z_{\text{pena}}) = 0$.

Next, we obtain the variance term under $H_0$. $Z_{\text{pena}} = \frac{1}{\sqrt{\hat{C}}} \frac{\hat{\boldsymbol{\alpha}}' \mathbf{z}_{\text{gwas}}}{\hat{\boldsymbol{\alpha}}' V \hat{\boldsymbol{\alpha}}}$, where $C = \mathrm{E}\left(\frac{1}{\hat{\boldsymbol{\alpha}}' V \hat{\boldsymbol{\alpha}}}\right)$. $\mathrm{var}(Z_{\text{pena}}) = \mathrm{var}(\frac{\hat{\boldsymbol{\alpha}}' \mathbf{z}_{\text{gwas}}}{\sqrt{\hat{C} \hat{\boldsymbol{\alpha}}' V \hat{\boldsymbol{\alpha}}}})$. Denote $\mathbf{u} = \frac{1}{\sqrt{\hat{C} \hat{\boldsymbol{\alpha}}' V \hat{\boldsymbol{\alpha}}}} \hat{\boldsymbol{\alpha}}$ and $\mathbf{w} = \mathbf{z}_{\text{gwas}}$. Thus, $\mathrm{var}(Z_{\text{pena}}) = \mathrm{var}(\mathbf{u}'\mathbf{w})$. Note that $\mathrm{var}(\mathbf{u}'\mathbf{w}) = \mathrm{E}((\mathbf{u}'\mathbf{w})^2) - \mathrm{E}^2(\mathbf{u}'\mathbf{w})$. Since $\mathbf{u}$ and $\mathbf{w}$ are independent, $\mathrm{E}(\mathbf{u}'\mathbf{w}) = \mathrm{E}(\mathbf{u}')\mathrm{E}(\mathbf{w})$. Again, $\mathrm{E}(\mathbf{w}) = \mathbf{0}$ under $H_0$. Thus, $\mathrm{var}(\mathbf{u}'\mathbf{w}) = \mathrm{E}((\mathbf{u}'\mathbf{w})^2)$ $= \mathrm{E}((\mathbf{u}'\mathbf{w}) \times (\mathbf{w}'\mathbf{u})) = \mathrm{tr}(\mathrm{E}(\mathbf{u}'\mathbf{w}\mathbf{w}'\mathbf{u})) = \mathrm{E}(\mathrm{tr}(\mathbf{u}'\mathbf{w}\mathbf{w}'\mathbf{u})) = \mathrm{E}(\mathrm{tr}(\mathbf{u}\mathbf{u}'\mathbf{w}\mathbf{w}')) = \mathrm{tr}(\mathrm{E}(\mathbf{u}\mathbf{u}'\mathbf{w}\mathbf{w}'))$ $= \mathrm{tr}(\mathrm{E}(\mathbf{u}\mathbf{u}')\mathrm{E}(\mathbf{w}\mathbf{w}'))$. Note that $\mathbf{w} = \mathbf{z}_{\text{gwas}} = \frac{1}{\sqrt{n}} G' \mathbf{y}$, where $G$ is the genotype matrix for the local SNPs in GWAS. The genotype and phenotype data are normalized to have a mean of zero and a variance of one. $\mathrm{cov}(\mathbf{z}_{\text{gwas}}) = \mathrm{cov}(\frac{1}{\sqrt{n}} G' \mathbf{y}) = \frac{1}{n} G' \mathrm{cov}(\mathbf{y}) G = \frac{\sigma_\epsilon^2}{n} G' G = V$. Under the assumption that a single gene has a negligible heritability for the phenotype, which is normalized to have variance one, $\mathrm{cov}(\mathbf{z}_{\text{gwas}}) = \mathrm{cov}(\mathbf{w}) = V$. Thus, $\mathrm{var}(Z_{\text{pena}})$ $= \mathrm{var}(\mathbf{u}'\mathbf{w}) = \mathrm{tr}(\mathrm{E}(\mathbf{u}\mathbf{u}')V) = \mathrm{tr}(V\mathrm{E}(\mathbf{u}\mathbf{u}')) = \mathrm{tr}(\mathrm{E}(V\mathbf{u}\mathbf{u}')) = \mathrm{E}(\mathrm{tr}(V\mathbf{u}\mathbf{u}')) = \mathrm{E}(\mathrm{tr}(\mathbf{u}'V\mathbf{u}))$ $= \mathrm{tr}(\mathrm{E}(\mathbf{u}'V\mathbf{u})) = \mathrm{E}(\mathbf{u}'V\mathbf{u}) = \mathrm{E}(\frac{1}{\hat{C}\hat{\boldsymbol{\alpha}}' V \hat{\boldsymbol{\alpha}}})$. Thus, $\mathrm{var}(Z_{\text{pena}}) = 1$ under $H_0$.

The LSE of $b_1$, $\frac{\sum_{i=1}^{n}(X_i-\bar{X})(y_i-\bar{y})}{\sum_{i=1}^{n}(X_i-\bar{X})^2}$, follows normal under the ideal regression model (Equation 5), since $y_i$ follows normal, $i = 1, \ldots, n$. For large sample sizes of contemporary GWAS, the unadjusted TWAS test statistic $Z_{\text{unadj}}$ follows normal under $H_0$. Based on Slutsky's theorem, if the adjustment factor $\hat{A}$ converges in probability to a constant, the adjusted test statistic $Z_{\text{pena}}$ should converge to a normal random variable in distribution under $H_0$. Since it is very challenging to show this condition analytically, we explore the null distribution using simulations. We assume the null distribution of $Z_{\text{pena}}$ is $N(0, 1)$ since the expectation and variance are shown to be zero and one under $H_0$. Using empirical density plots and QQ plots, we demonstrate that the null distribution matches excellently with $N(0, 1)$ (results described in the simulation study section in the main text).

#### 2.7 An adjusted variance approach

Xue et al. [12] proposed an adjusted TWAS approach based on individual-level data. To tune for the uncertainty of predicted expression, they considered an adjusted variance of the estimated TWAS effect size [12]. Let $\hat{b}$ stand for the OLS estimate of $b$ based on the working regression (Equation 6). Suppose $\text{var}(\hat{b})$ denotes the variance of $\hat{b}$ based on the unadjusted TWAS. The calibrated variance [12] takes the following form: $\text{var}_c(\hat{b}) = \text{var}(\hat{b})\big(1 + \hat{b}^2 \frac{n}{n_1} \frac{\sigma_\eta^2}{\sigma_e^2}\big)$. The adjusted variance $\text{var}_c(\hat{b})$ is inflated than the initial variance $\text{var}(\hat{b})$. Here, $n_1$ and $n$ are the transcriptome and GWAS data sample sizes; $\sigma_\eta^2$ and $\sigma_e^2$ are the variances of the residual terms in the linear models (Equations 1 and 6) based on the transcriptome and GWAS data, respectively. Xue et al. [12] considered a less realistic scenario in the first stage of TWAS, where the number of local SNPs for a gene is much smaller than the sample size, allowing the simple multiple linear regression to predict the expression. On the contrary, the contemporary TWAS approaches consider penalized regression in the first stage of TWAS to incorporate many SNPs. However, the proposed adjusted variance can also be implemented for a penalized regression in the first stage. To modify the adjusted variance approach proposed by Xue et al., we consider the Lasso while regressing the expression on the genotypes of local SNPs (Equations 1). We follow Reid et al. [37] to estimate the error variance in a penalized regression. We devise the estimator of $\sigma_\eta^2$ as $\frac{1}{n_1 - p_1}\sum_{i=1}^{n_1}\hat{\eta}_i^2$, where $p_1$ is the number of local SNPs selected in the Lasso variable selection, i.e., the number of non-zero estimated coefficients in $\hat{\boldsymbol{\alpha}}$ obtained by Lasso based on the complete transcriptome data. Here, $\hat{\eta}_i$, $i = 1, \ldots, n_1$, denote the estimated residuals from Lasso. $\sigma_e^2$ is estimated from the linear working regression in Equation 6. We note that the adjusted variance approach requires individual-level GWAS data for implementation.

#### 2.8 Connection to testing for genetic correlation between outcome phenotype and gene expression

De Leeuw et al. [8] showed that the genetic relationship between gene expression and the GWAS phenotype can be viewed as the genetic correlation between the pair of traits. Their main purpose was to show that the standard TWAS does not test whether the genetic correlation is zero under the null. It rather tests $H_0$: genetic correlation $= -\hat{C}$, which can take any value (including non-zero values). The main reason behind that is that the standard TWAS ignores the uncertainty in the predicted expression. We show that our framework explicitly tests whether the genetic correlation is zero, not an arbitrary value under the null. The main reason is that we test for the regression coefficient in the ideal regression, but not in the working regression. The former denotes the effect of the true genetic component of gene expression (GReX) on the GWAS outcome.

In the reference transcriptome data: $E = \boldsymbol{g}_{ref}'\boldsymbol{\alpha} + \eta$, where $E$ denotes the gene expression, $\boldsymbol{g}_{ref}$ denotes the cis SNPs' genotype vector, $\boldsymbol{\alpha}$ denotes the cis SNPs' effects. For a GWAS individual, $x = \boldsymbol{g}'\boldsymbol{\alpha}$ denotes the true GReX, where $\boldsymbol{g}$ denotes the individual's genotypes for the cis SNPs. Instead of the true GReX, we have the observed predicted GReX: $X = \boldsymbol{g}'\hat{\boldsymbol{\alpha}}$, where $\hat{\boldsymbol{\alpha}}$ is the estimated $\boldsymbol{\alpha}$ in the transcriptome data. According to working regression (dropping the intercept), $y = Xb + e$, where $y$ is the GWAS outcome phenotype. From this linear regression: $b = \frac{cov(X,y)}{V(X)}$ [also shown in De Leeuw et al.].

Thus, in standard TWAS, testing $H_0 : b = 0$ vs $H_1 : b \neq 0$ is equivalent to testing $H_0 : cov(X, y) = 0$ vs $H_1 : cov(X, y) \neq 0$. In the ideal regression framework: $y = x\beta + \epsilon$. Hence, $\beta = \frac{cov(x,y)}{V(x)}$. Thus, in the adjusted TWAS based on ideal regression, testing $H_0 : \beta = 0$ vs $H_1 : \beta \neq 0$ is equivalent to testing $H_0 : cov(x, y) = 0$ vs $H_1 : cov(x, y) \neq 0$. In the next steps, De Leeuw et al. considered a decomposition of the GWAS outcome into two parts: one contribution due to the same cis-SNPs marginally, and the other part is the unexplained part. Therefore, they consider: $y = \boldsymbol{g}'\boldsymbol{\alpha}_y + \psi = y_g + \psi$. Here, $y_g$ is the genetic component of the GWAS outcome due to the cis-SNPs with marginal effect $\boldsymbol{\alpha}_y$. In the subsequent steps, the authors show that $cov(X, y) = cov(X, y_g)$. Following the same steps, it can be shown that $cov(x, y) = cov(x, y_g)$ in our setup, where $x$ is true GReX. Thus, $cov(x, y)$ can be interpreted as the genetic covariance between the gene expression and the GWAS outcome phenotype. Hence, testing $H_0 : \beta = 0$ vs $H_1 : \beta \neq 0$ is equivalent to testing $H_0 : cov(x, y_g) = 0$. That is, equivalently, whether the genetic correlation between expression and outcome is zero. De Leeuw et al. (PLoS Genetics, 2023) derived that this is the correct hypothesis-testing framework for TWAS, specifically when interpreting the TWAS signal as a genetic relationship between gene expression and the GWAS outcome phenotype. In standard TWAS, $H_0 : cov(X, y) = cov(X, y_g) = 0$ becomes equivalent to $H_0 : cov(x, y_g) = -\hat{C}$, where $\hat{C}$ can take an arbitrary value, including non-zero values. The interpretation of $\hat{C}$ is provided in the paper, which we skip for brevity. Therefore, in standard TWAS, we test for the actual genetic correlation being equal to $-\hat{C}$, not necessarily zero under the null.

Therefore, the null hypothesis, which we tested in our proposed PenaTWAS based on the ideal regression, is indeed the correct null hypothesis with respect to the local genetic correlation between gene expression and the GWAS phenotype.

### 3 Simulation setting

We implemented the *hapgen* [22] software to simulate genotype data in the transcriptome and GWAS data. We used simulation models to generate the gene expression in the transcriptome data and the outcome phenotype in the GWAS data. Gusev et al. [4] analyzed the transcriptome data from the Young Finish Sequencing (YFS) study to identify all the locally heritable genes in the whole blood tissue. They provided the results from the analysis in the software package Fusion. Since we test one gene at a time in traditional TWAS, we selected a locally heritable gene, DGCR8, on chromosome 22 from the YFS data analysis for simplicity in computation. To run *hapgen*, we considered a Finnish (European) population's LD structure for the genomic region neighboring DGCR8. We considered 300 local SNPs, with MAF $> 0.05$, randomly selected from the 1MB local region surrounding DGCR8. To mimic the TWAS setup, we generate the transcriptome and GWAS data independently with non-overlapping individuals from the same population. We chose the transcriptome data sample size of $n_1 = 500$ and the GWAS data sample size of $n = 10,000$. *Hapgen* is a widely used method to generate genotype data mimicking real-life populations. Using *hapgen* allows the creation of non-overlapping and independent individuals' genotype data across the iterations in a given simulation scenario.

After generating the genotype data, we simulate the gene expressions of DGCR8 in the transcriptome data based on the local SNPs' genotypes for the $n_1$ individuals. Genotype values for each local SNP are normalized to have a mean of zero and a variance of one.

We consider a linear model to simulate the expression: $E = \mathbf{g}_{ref}^{'}\boldsymbol{\alpha} + \eta$. Here, $E$ denotes the expression, $\mathbf{g}_{ref}$ denotes the genotype values for the local SNPs, and $\boldsymbol{\alpha}$ denotes the effect size vector for the local SNPs on expression. Assuming that $\text{var}(E) = 1$, we consider $\eta \sim N(0, 1 - h_E^2)$, where $h_E^2$ is the local heritability of the expression due to the SNPs. We assume that 5% of the local SNPs affect the expression. If $m_c$ denotes the number of such SNPs, each element of $\boldsymbol{\alpha}$ follows $N(0, \frac{h_E^2}{m_c})$. We chose $h_E^2 = 10\%, 20\%$. We simulated a few different choices of $\boldsymbol{\alpha}$. We next simulate the outcome in GWAS data using the linear model: $y = \beta_1 x + \epsilon$. Here, $y$ denotes the phenotype, $x = \mathbf{g}^{'}\boldsymbol{\alpha}$ denotes the true genetic component of expression, where $\mathbf{g}$ is the genotype vector for the local SNPs for a GWAS individual, and $\boldsymbol{\alpha}$ is the same as in the transcriptome data. The genetic component of expression for one gene should not explain a significant proportion of the variance of a complex phenotype. Hence, we assume that $\text{var}(y) = \text{var}(\epsilon) = 1$. We selected ten choices of $\beta_1$ as $0.02, 0.04, \ldots, 0.18, 0.2$ to study power and estimation of effect sizes, and $\beta_1 = 0$ for evaluating the type I error rate. To simulate a case-control phenotype, we threshold the continuous phenotype simulated. We use the threshold such that 25% of the individuals with higher values of the continuous phenotype become cases, and the remaining 75% of the individuals become controls.

#### 3.1 Analysis pipeline

We implemented the GCTA software [23] to compute the p-value of testing the null hypothesis that the local heritability of the gene is zero versus the alternative hypothesis that it is positive. We applied the BH FDR controlling procedure to the p-values obtained from 6,000 iterations under a given simulation scenario. In a simulation scenario, we regarded an iteration for downstream analysis if the gene was heritable in the iteration based on the FDR correction with an FDR threshold of 0.05. In the simulation settings considered, we always found more than 1800 iterations to pass the correction, based on which we evaluated the various measures for the inference. For unadjusted TWAS, we implemented penalized regression using the Lasso, adaptive Lasso, and Elastic net penalty to estimate the prediction model for the genetically regulated component of expression based on the transcriptome data. To perform a TWAS based on individual-level data, we imputed the expression using the prediction model in the GWAS data. Then, we tested for an association between the predicted expression and the outcome using linear or logistic regression. We obtained the adjustment factor for PenaTWAS by applying the adaptive Lasso and residual bootstrapping. Next, we computed the summary-level association data for the local SNPs in GWAS and calculated the test statistic for an unadjusted TWAS based on summary-level data. In an iteration, we utilized the LD matrix of local SNPs estimated from the transcriptome data in the next iteration. This strategy imitates the TWAS procedure based on summary-level data, where LD structure is calculated from external reference data, such as 1000 Genomes data. The latter has a sample size of 503 for the European population. To assess power and type I error rate, we considered the standard choice of significance level as 0.05. We examined $0\%, 5\%, 10\%$ trimmed means of the bootstrapped estimates while estimating the denominator of the adjustment factor in PenaTWAS. We denote the PenaTWAS that utilizes $0\%, 5\%, 10\%$ trimmed means as PenaTWAS0, PenaTWAS5, and PenaTWAS10, respectively. We used $T = 300$ residual bootstrap samples in PenaTWAS. To explore the normality of the null distribution of PenaTWAS statistics, we constructed the QQ and empirical density plots for the test statistic values under the null hypothesis of no association. To compare PenaTWAS with CoMM, PMR-Egger, Otters, and HMAT, we used the software packages available for these methods from the GitHub. While implementing HMAT, we combined three different prediction models for the gene expression: ridge regression, Lasso, and mixed-effects model as implemented in the GEMMA software. We compared PenaTWAS with CoMM, PMR-Egger, and HMAT for individual-level TWAS and PMR-Egger, Otters, and HMAT for summary-TWAS.

Figure S1: Comparison of type I error rate and power between the different versions of PenaTWAS and the adjusted variance approach based on individual-level data. Here, PenaTWAS0, PenaTWAS5, and PenaTWAS10 denote the PenaTWAS approaches that use 0%, 5%, 10% trimmed means based on the bootstrap sample while estimating the adjustment factor. PENAtaylor stands for the Taylor series expanded version of estimating the adjustment factor in PenaTWAS. AdjVar represents the modified adjusted variance approach. The heritability of gene expression is 10%. The first choice of true eQTL effects was considered.

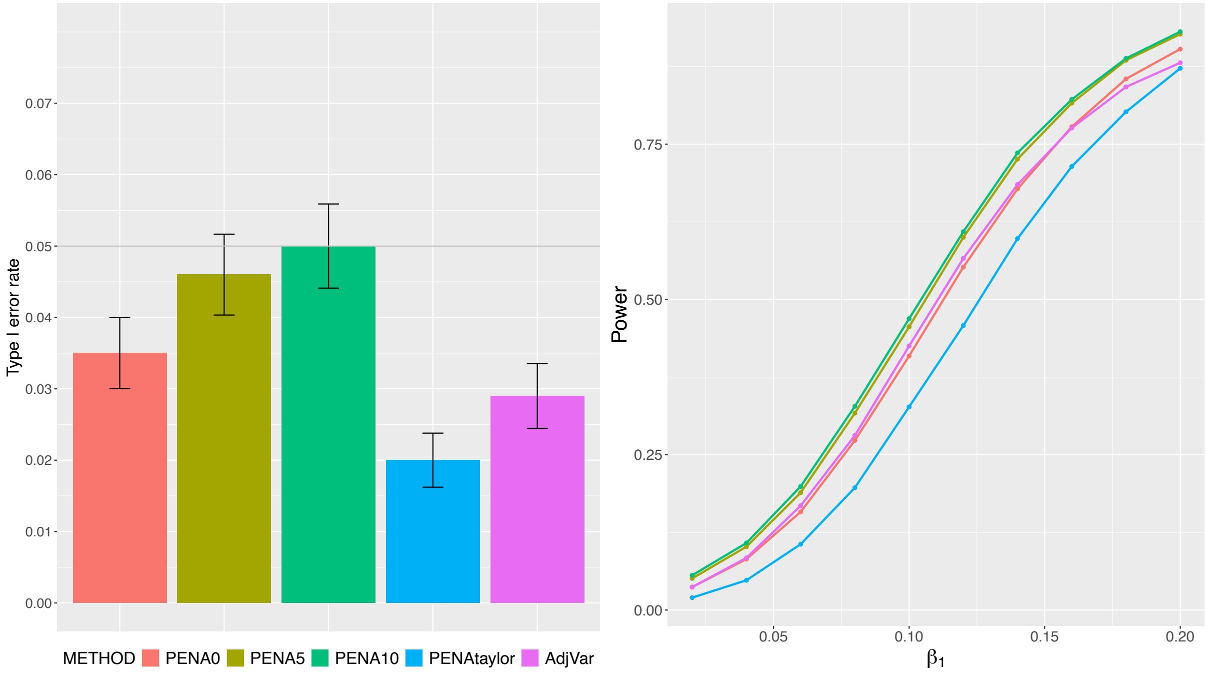

Figure S2: Comparison of type I error rate and power between the different PenaTWAS approaches based on summary-level data. Here, PenaTWAS0, PenaTWAS5, PenaTWAS10 denote the PenaTWAS approaches that use $0\%, 5\%, 10\%$ trimmed means based on the bootstrap sample while estimating the adjustment factor. PENAtaylor stands for the Taylor series expanded version of estimating the adjustment factor in PenaTWAS. The heritability of gene expression is 10%. The first choice of true eQTL effects was considered.

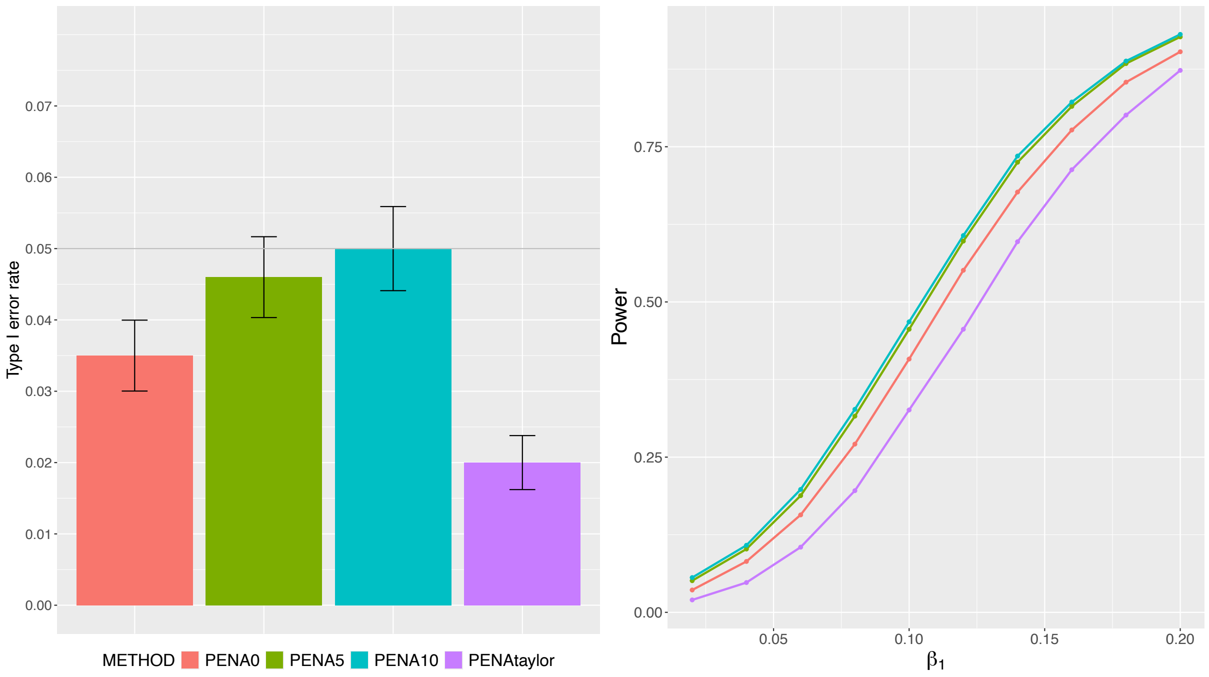

Figure S3: Comparison of type I error rate and power between individual-level PenaTWAS, CoMM, PMR-Egger, and HMAT. HMAT is an unadjusted TWAS approach, and the others are adjusted TWAS approaches. The heritability of gene expression is 10%.

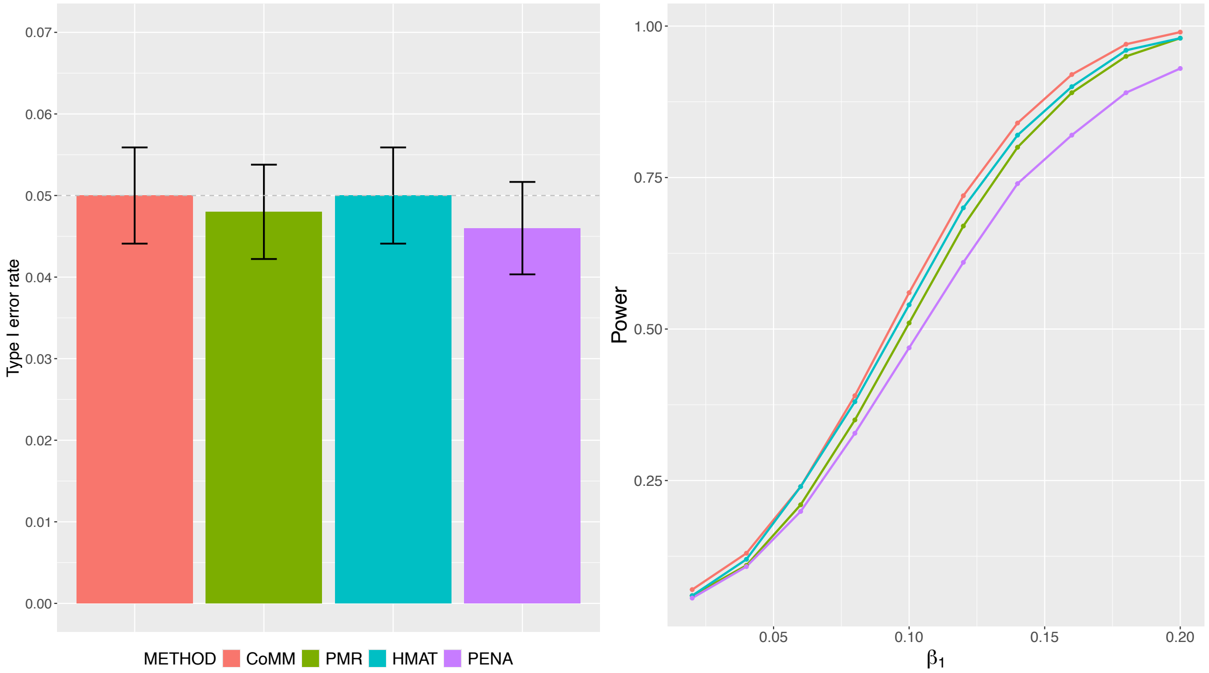

Figure S4: Comparison of type I error rate and power between the different versions of PenaTWAS and adjusted variance approach based on individual-level data. Here, PenaTWAS0, PenaTWAS5, PenaTWAS10 denote the PenaTWAS approaches that use 0%, 5%, 10% trimmed means based on bootstrap sample while estimating the adjustment factor. PENAtaylor stands for the Taylor series expanded version of estimating the adjustment factor in PenaTWAS. AdjVar represents the modified adjusted variance approach. The heritability of gene expression is 10%. The second choice of true eQTL effects was considered.

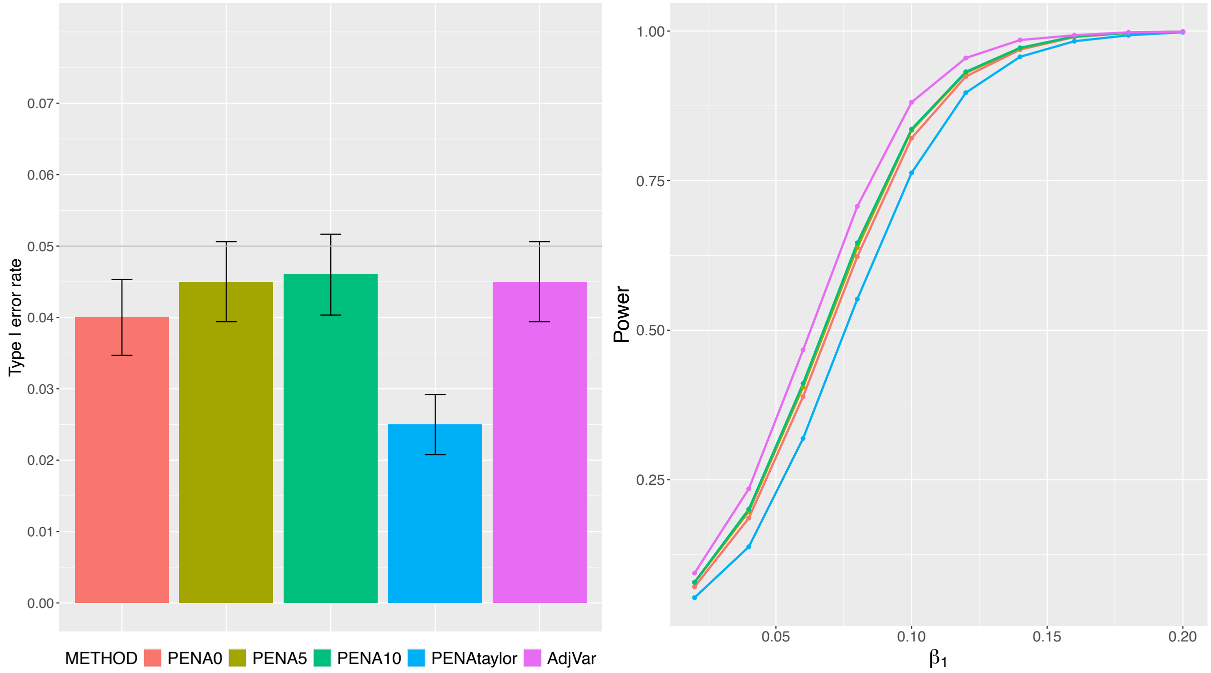

Figure S5: Comparison of type I error rate and power between the unadjusted TWAS approaches and the adjusted TWAS approach PenaTWAS based on individual-level data. Results for the gold standard TWAS, in which true eQTL effects are plugged in, are also presented. The unadjusted TWAS approaches are Lasso, ALasso, and Enet TWAS, depending on the type of penalty function used. The heritability of gene expression is 10%. The second choice of true eQTL effects was considered.

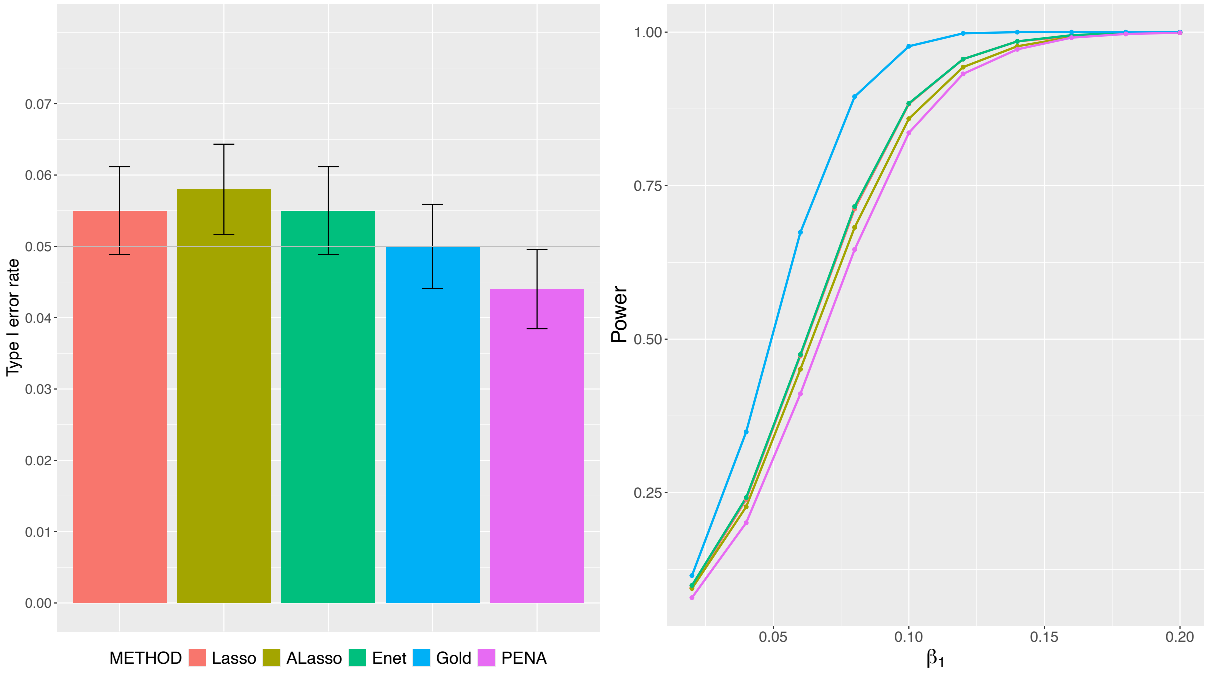

Figure S6: Comparison of type I error rate and power between the different versions of PenaTWAS based on summary-level data. Here, PenaTWAS0, PenaTWAS5, PenaTWAS10 denote the PenaTWAS approaches that use 0%, 5%, 10% trimmed means while estimating the adjustment factor. PENAtaylor stands for the Taylor series expanded version of estimating the adjustment factor in PenaTWAS. The heritability of gene expression is 10%. The second choice of true eQTL effects was considered.

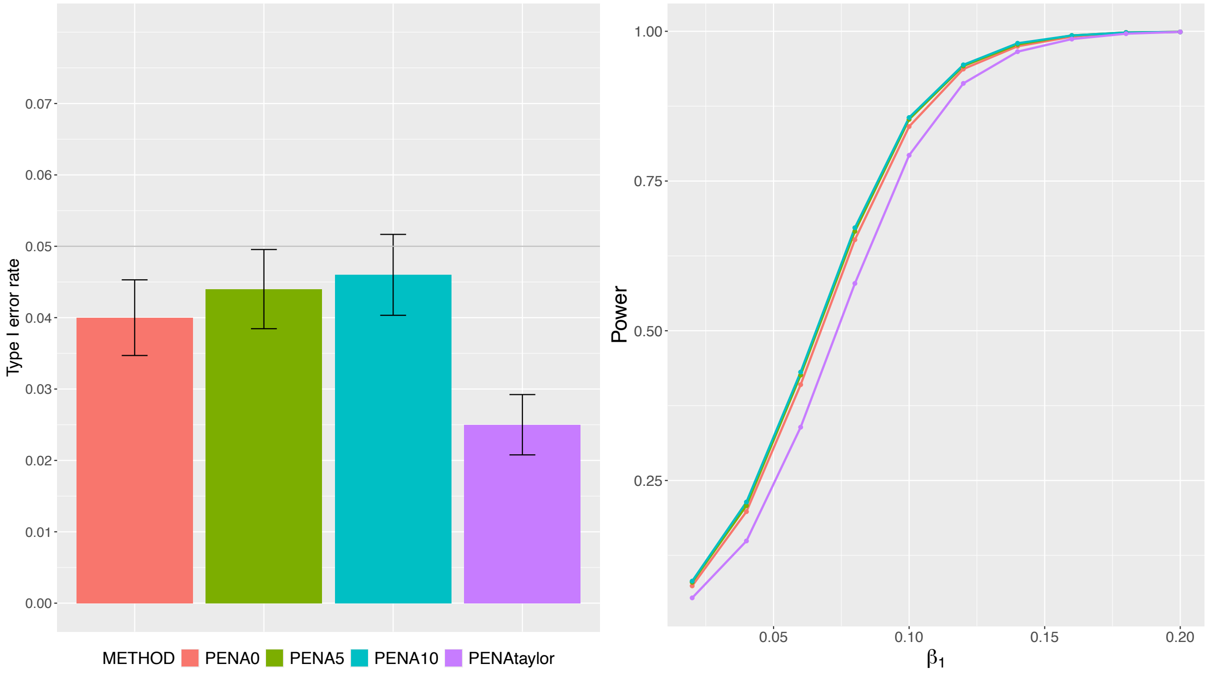

Figure S7: Comparison of type I error rate and power between the unadjusted TWAS approaches and the adjusted TWAS approach PenaTWAS based on summary-level data. Results for the gold standard TWAS, in which true eQTL effects are plugged in, are also presented. The unadjusted TWAS approaches are Lasso, ALasso, and Enet TWAS, depending on the type of penalty function used. The heritability of gene expression is 10%. The second choice of true eQTL effects was considered.

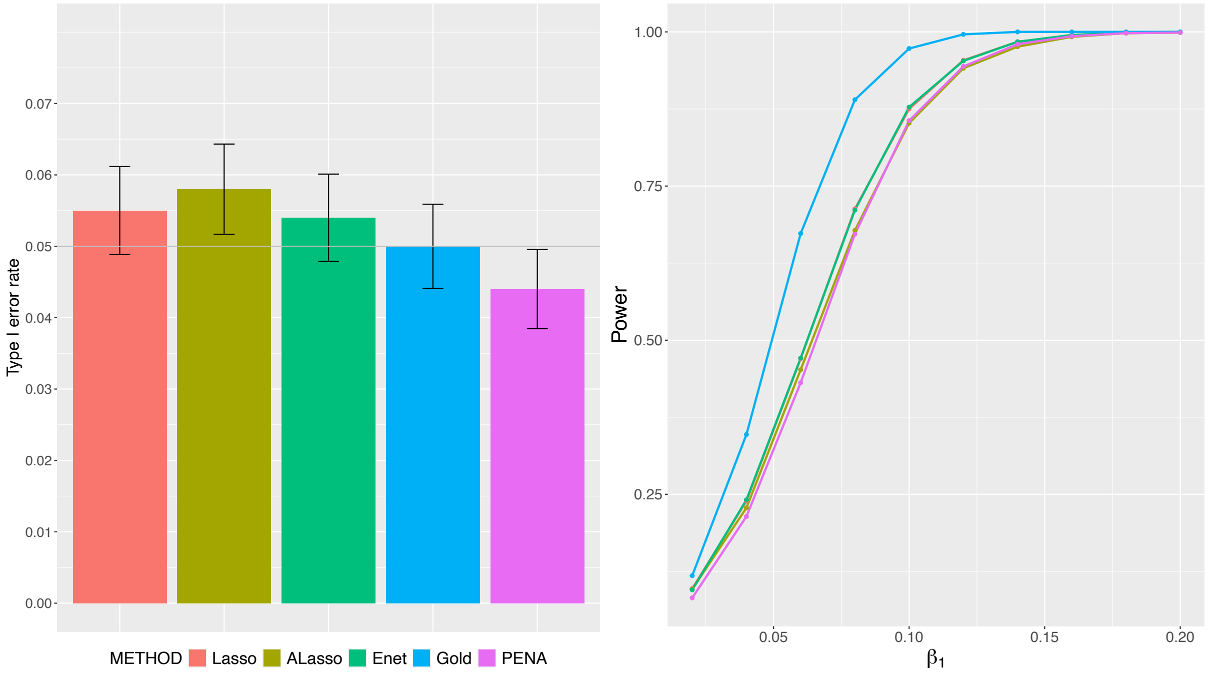

Figure S8: Comparison of type I error rate and power between the different versions of PenaTWAS and adjusted variance approach based on individual-level data. Here, PenaTWAS0, PenaTWAS5, PenaTWAS10 denote the PenaTWAS approaches that use 0%, 5%, 10% trimmed means based on the bootstrap sample while estimating the adjustment factor. PENAtaylor stands for the Taylor series expanded version of estimating the adjustment factor in PenaTWAS. AdjVar represents the modified adjusted variance approach. The heritability of gene expression is 20%.

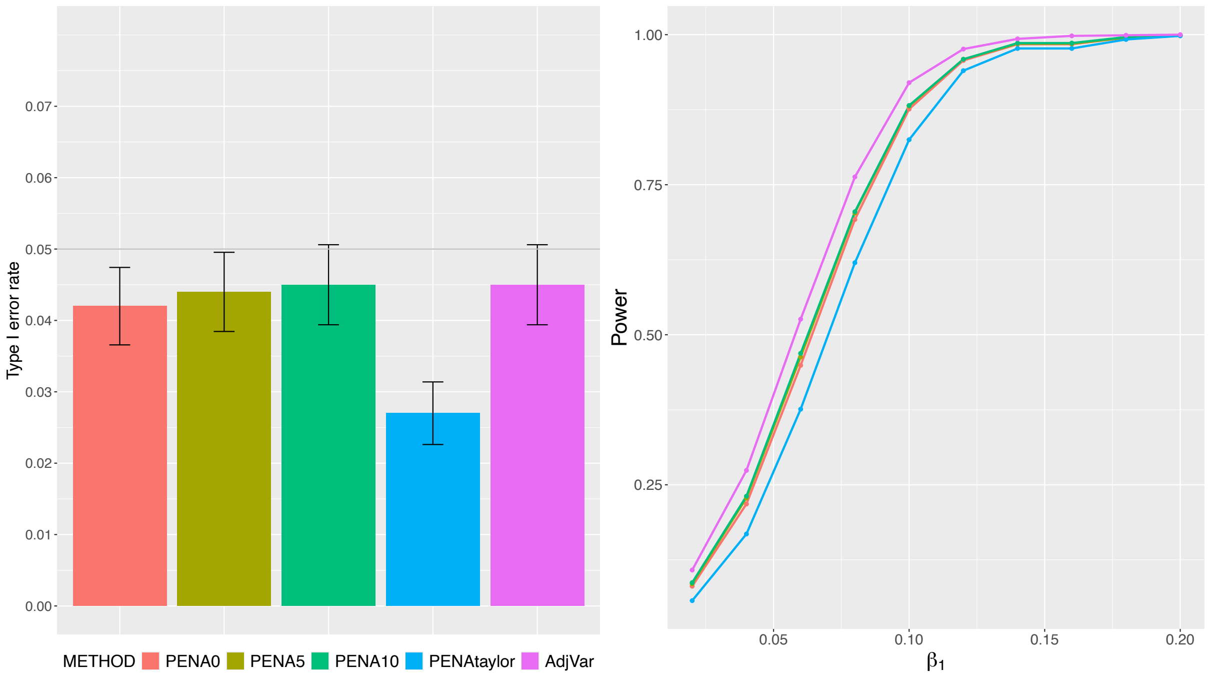

Figure S9: Comparison of type I error rate and power between the unadjusted TWAS approaches and the adjusted TWAS approach PenaTWAS based on individual-level data. Results for the gold standard TWAS, in which true eQTL effects are plugged in, are also presented. The unadjusted TWAS approaches are Lasso, ALasso, and Enet TWAS, depending on the type of penalty function used. The heritability of gene expression is 20%.

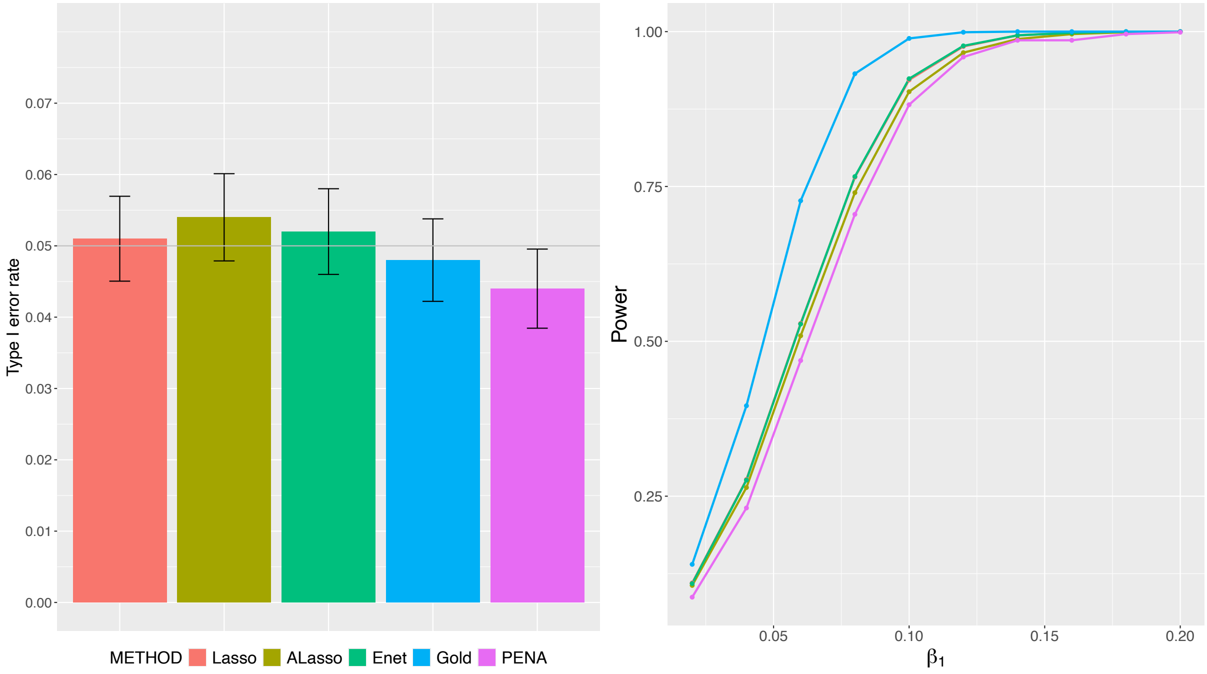

Figure S10: Comparison of type I error rate and power between the different versions of PenaTWAS approaches based on summary-level data. Here, PenaTWAS0, PenaTWAS5, PenaTWAS10 denote the PenaTWAS approaches that use $0\%, 5\%, 10\%$ trimmed means based on the bootstrap sample while estimating the adjustment factor. PENAtaylor stands for the Taylor series expanded version of estimating the adjustment factor in PenaTWAS. The heritability of gene expression is $20\%$.

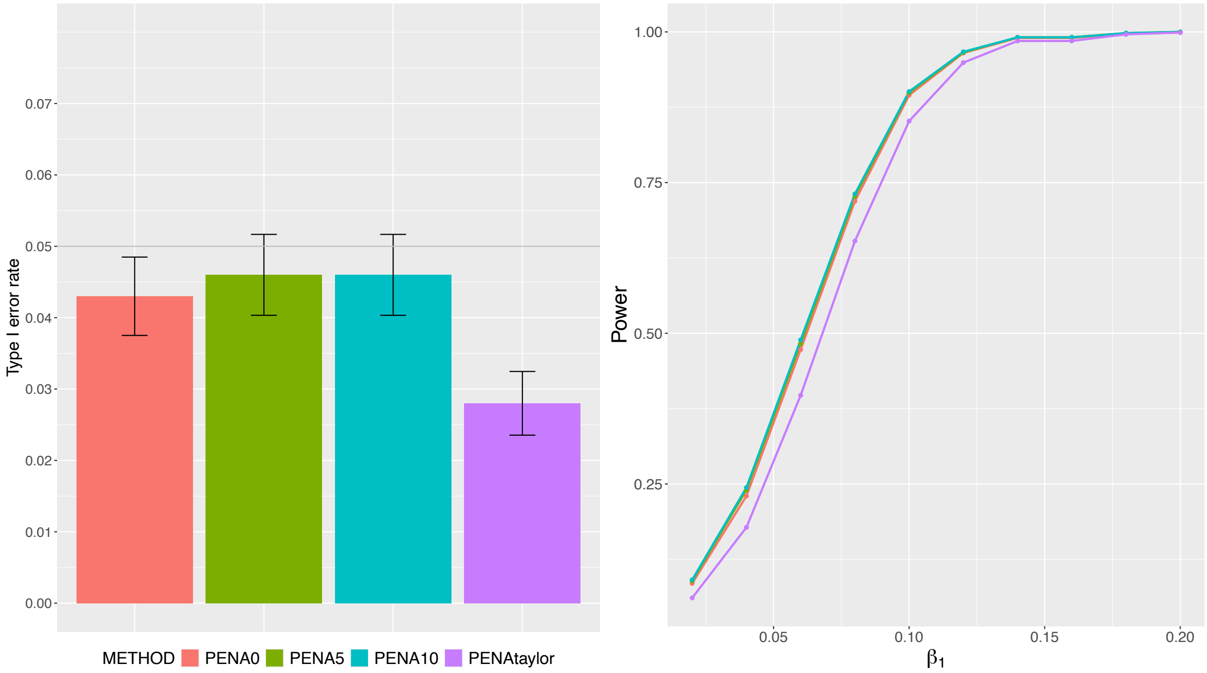

Figure S11: Comparison of type I error rate and power between the unadjusted TWAS approaches and the adjusted TWAS approach PenaTWAS based on summary-level data. Results for the gold standard TWAS, in which true eQTL effects are plugged in, are also presented. The unadjusted TWAS approaches are referred to as Lasso, ALasso, and Enet TWAS, depending on the type of penalty function used. The heritability of gene expression is 20%.

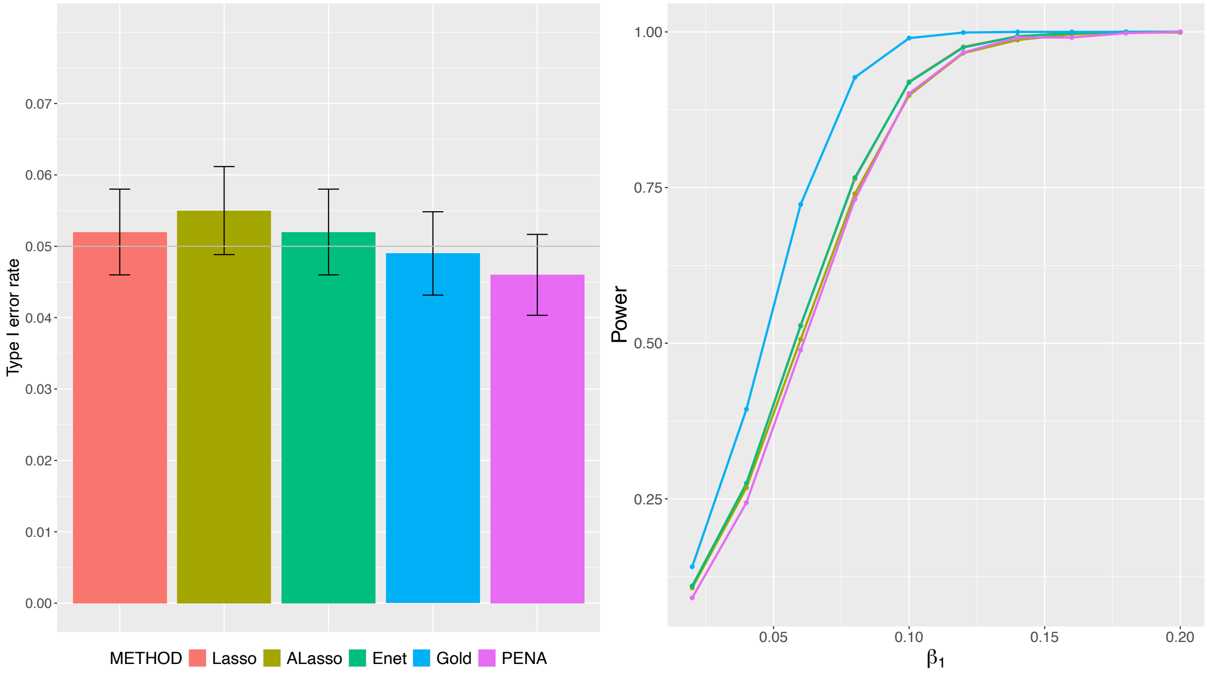

Figure S12: The adjustment factors to obtain the $Z$ statistics for PenaTWAS based on individual-level data in the three simulation scenarios considered for a continuous phenotype. In the first two scenarios, the gene expression local heritability is 10%, while in the third scenario, it is 20%.

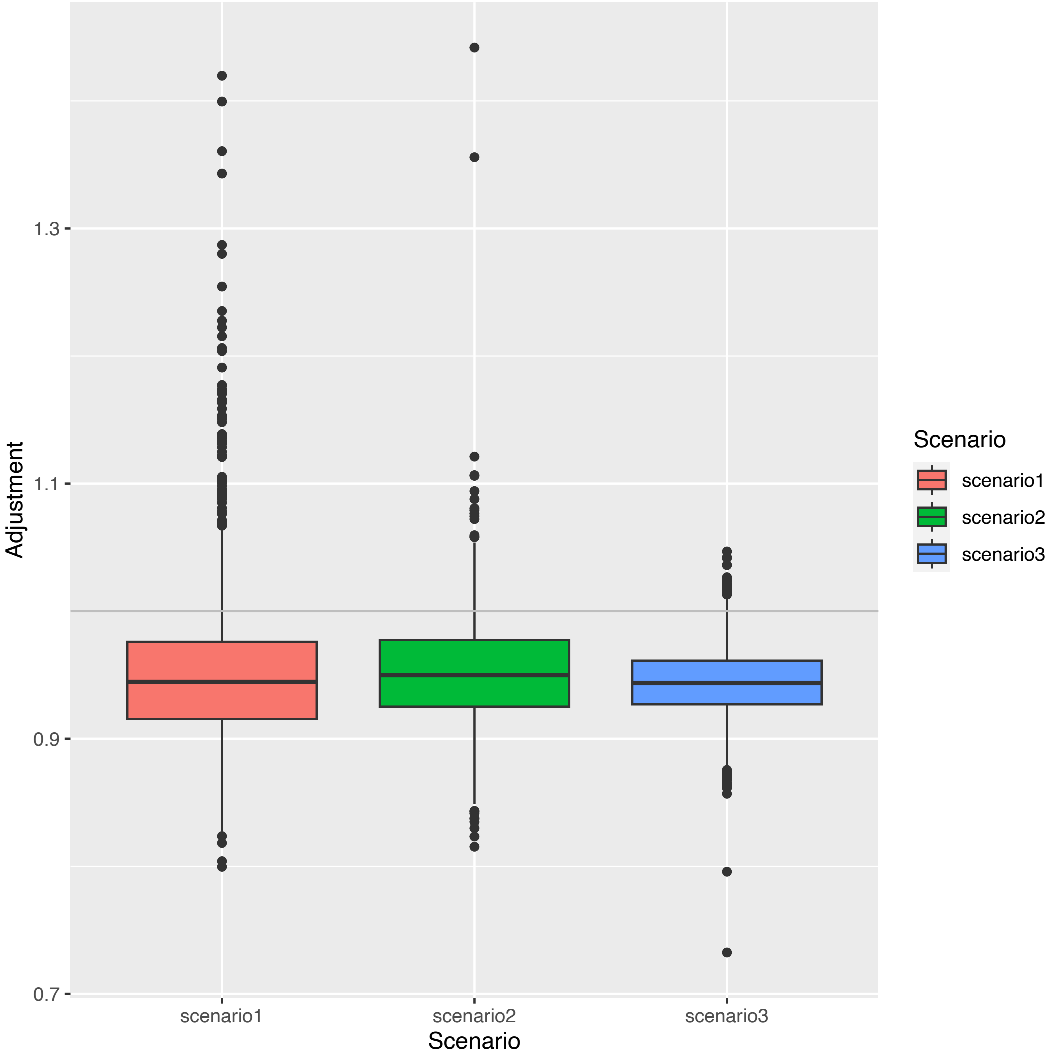

Figure S13: Empirical density plots for the PenaTWAS test statistics under the null hypothesis of no association obtained from the individual-level TWAS for a continuous phenotype. The gene expression heritability is 10% for the first choice of the true eQTL effects. In PenaTWAS test statistics, $0\%, 5\%, 10\%$ trimmed means based on the bootstrap sample were considered while estimating the adjustment factor.

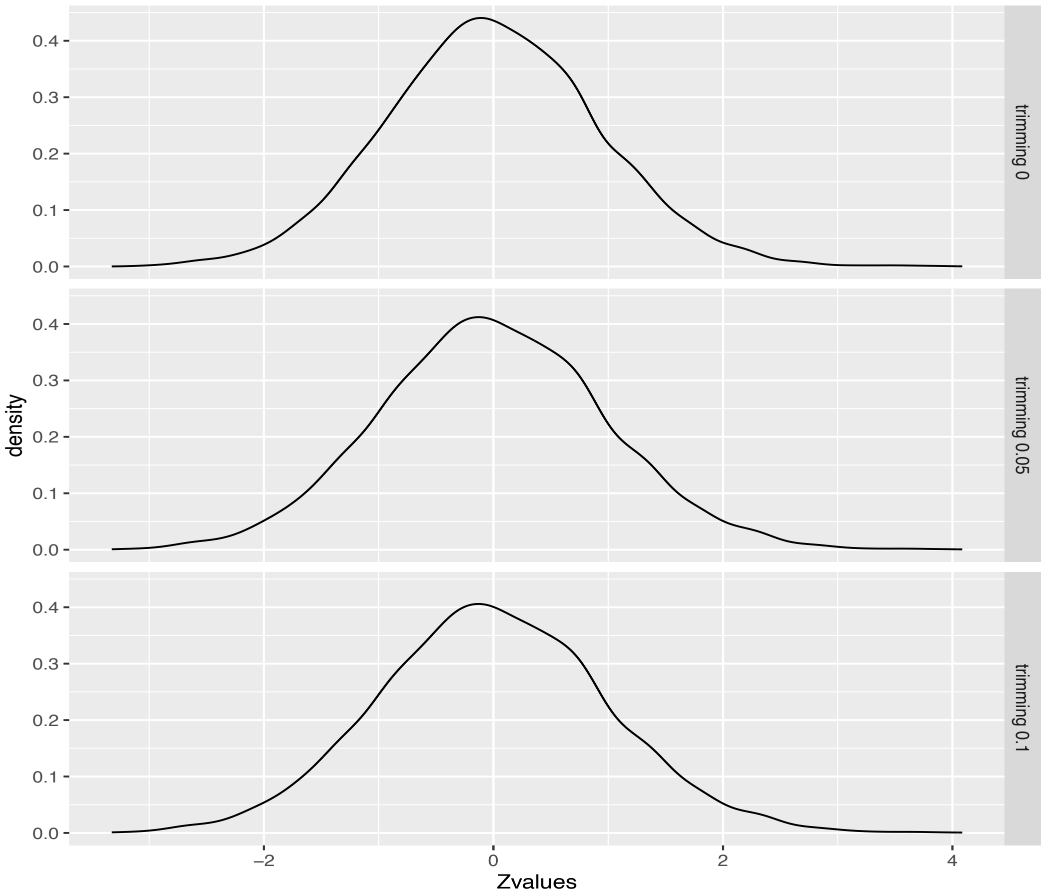

Figure S14: Empirical density plots for the PenaTWAS test statistics under the null hypothesis of no association obtained from the summary-TWAS for a continuous phenotype. The gene expression heritability is 10% for the first choice of the true eQTL effects. In PenaTWAS test statistics, $0\%, 5\%, 10\%$ trimmed means based on the bootstrap sample were considered while estimating the adjustment factor.

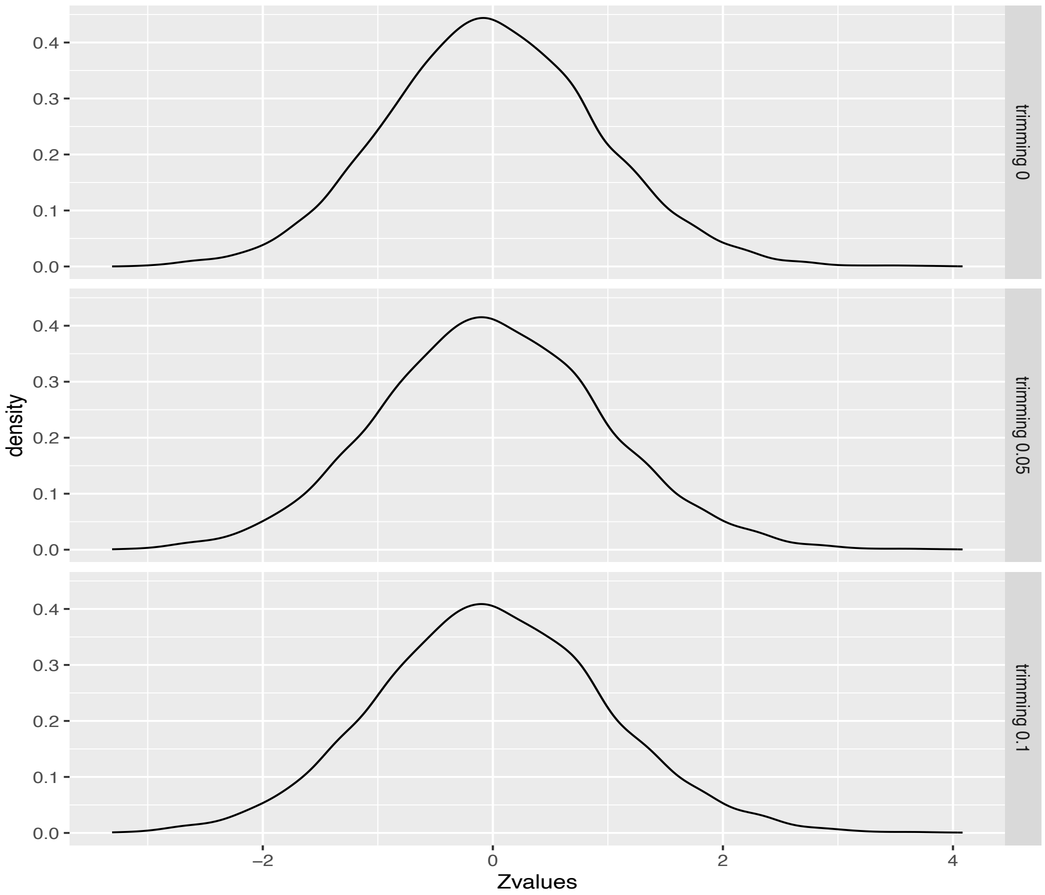

Figure S15: QQ plots for the PenaTWAS test statistics under the null hypothesis of no association obtained from the individual-level TWAS for a continuous phenotype. The gene expression heritability is 10% for the first choice of the true eQTL effects. In PenaTWAS test statistics, $0\%, 5\%, 10\%$ trimmed means based on the bootstrap sample were considered while estimating the adjustment factor.

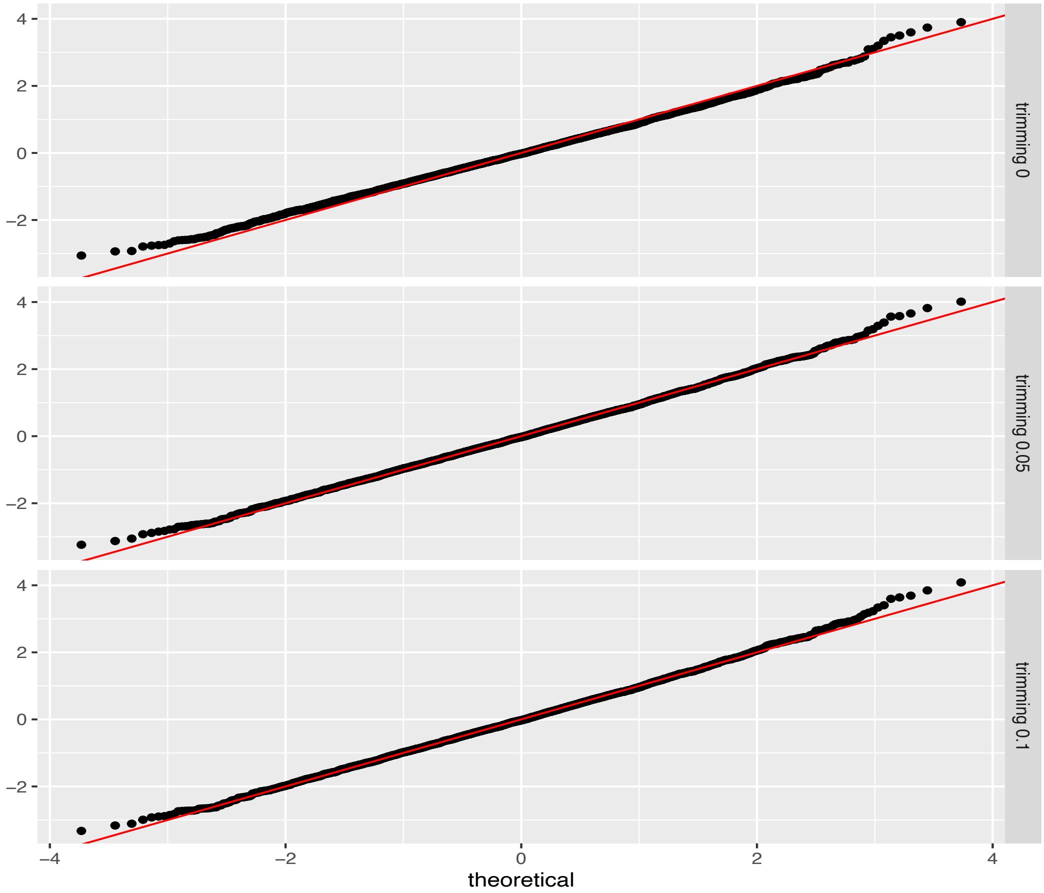

Figure S16: QQ plots for the PenaTWAS test statistics under the null hypothesis of no association obtained from the summary-TWAS for a continuous phenotype. The gene expression heritability is 10% for the first choice of the true eQTL effects. In PenaTWAS test statistics, $0\%, 5\%, 10\%$ trimmed means based on the bootstrap sample were considered while estimating the adjustment factor.

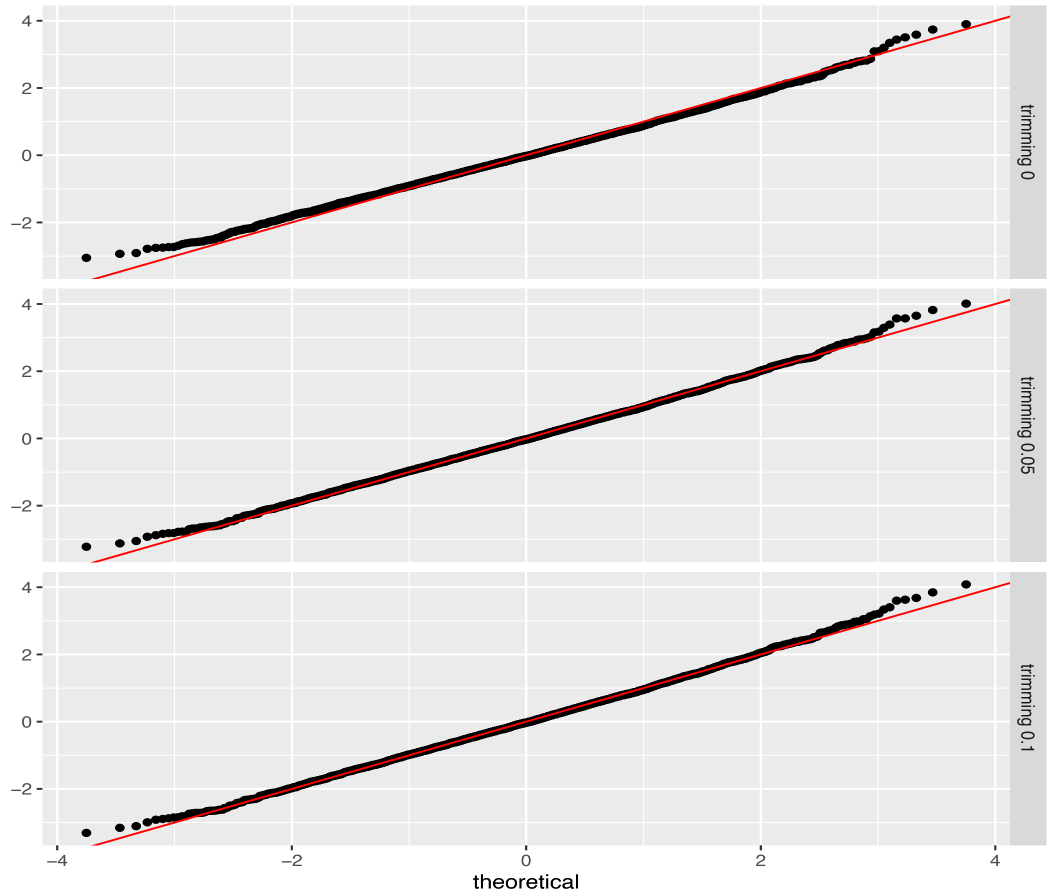

Table S1: Summary of relative bias in estimates of the effect size of the genetic component of expression on the outcome. The unadjusted TWAS approaches are Lasso, ALasso, and Enet TWAS, depending on the type of penalty function used. PENA denotes the adjusted TWAS approach PenaTWAS. The heritability of gene expression is 10%. The first choice of true eQTL effects was considered. The mean and s.d. of the relative bias obtained across simulation iterations are provided.

| $\beta_1$ | Lasso | Enet | ALasso | PENA |
|---|---|---|---|---|
| 0.02 | 17 (347) | 23 (352) | -32 (191) | 7 (356) |
| 0.04 | 15 (180) | 20 (185) | -32 (97) | 7 (185) |
| 0.06 | 15 (127) | 19 (131) | -32 (67) | 6 (130) |
| 0.08 | 14 (102) | 18 (105) | -32 (52) | 6 (105) |
| 0.1 | 14 (87) | 18 (91) | -32 (43) | 6 (90) |
| 0.12 | 14 (79) | 18 (82) | -32 (38) | 6 (81) |
| 0.14 | 14 (73) | 17 (76) | -32 (34) | 6 (76) |
| 0.16 | 14 (69) | 17 (72) | -32 (32) | 6 (71) |
| 0.18 | 14 (66) | 17 (69) | -32 (30) | 6 (69) |
| 0.2 | 14 (63) | 17 (66) | -32 (28) | 6 (66) |

Table S2: Square root of mean squared error (RMSE) of the estimates of the effect size of the genetic component of expression on the outcome. The unadjusted TWAS approaches are Lasso, ALasso, and Enet TWAS, depending on the type of penalty function used. PENA denotes the adjusted TWAS approach PenaTWAS. The heritability of gene expression is 10%. The first choice of true eQTL effects was considered.

| $\beta_1$ | Lasso | Enet | ALasso | PENA |
|---|---|---|---|---|
| 0.02 | 0.07 | 0.07 | 0.04 | 0.07 |
| 0.04 | 0.07 | 0.07 | 0.04 | 0.07 |
| 0.06 | 0.08 | 0.08 | 0.04 | 0.08 |
| 0.08 | 0.08 | 0.09 | 0.05 | 0.08 |
| 0.1 | 0.09 | 0.09 | 0.05 | 0.09 |
| 0.12 | 0.1 | 0.1 | 0.06 | 0.1 |
| 0.14 | 0.1 | 0.11 | 0.07 | 0.11 |
| 0.16 | 0.11 | 0.12 | 0.07 | 0.11 |
| 0.18 | 0.12 | 0.13 | 0.08 | 0.12 |
| 0.2 | 0.13 | 0.14 | 0.08 | 0.13 |

Table S3: Summary of estimated effect sizes of the genetic component of expression on the outcome. The unadjusted TWAS approaches are Lasso, ALasso, and Enet TWAS, depending on the penalty function used. PENA denotes the adjusted TWAS approach, PenaTWAS. The heritability of gene expression is 10%. The second choice of true eQTL effects was considered. The mean and s.d. of the estimates obtained across simulation iterations are provided.

| $\beta_1$ | Lasso | Enet | ALasso | PENA |
|---|---|---|---|---|
| 0.02 | 0.02 (0.04) | 0.02 (0.04) | 0.02 (0.03) | 0.02 (0.04) |
| 0.04 | 0.04 (0.04) | 0.05 (0.04) | 0.03 (0.03) | 0.04 (0.04) |
| 0.06 | 0.07 (0.04) | 0.07 (0.04) | 0.05 (0.03) | 0.06 (0.04) |
| 0.08 | 0.09 (0.04) | 0.09 (0.04) | 0.06 (0.03) | 0.08 (0.04) |
| 0.1 | 0.11 (0.05) | 0.12 (0.05) | 0.08 (0.03) | 0.1 (0.05) |
| 0.12 | 0.14 (0.05) | 0.14 (0.05) | 0.1 (0.03) | 0.12 (0.05) |
| 0.14 | 0.16 (0.05) | 0.16 (0.05) | 0.11 (0.04) | 0.14 (0.05) |
| 0.16 | 0.18 (0.06) | 0.19 (0.06) | 0.13 (0.04) | 0.17 (0.06) |
| 0.18 | 0.21 (0.06) | 0.21 (0.06) | 0.15 (0.04) | 0.19 (0.06) |
| 0.2 | 0.23 (0.07) | 0.23 (0.07) | 0.16 (0.04) | 0.21 (0.07) |

Table S4: Summary of relative bias in estimates of the effect size of the genetic component of expression on the outcome. The unadjusted TWAS approaches are Lasso, ALasso, and Enet TWAS, depending on the type of penalty function used. PENA denotes the adjusted TWAS approach PenaTWAS. The heritability of gene expression is 10%. The second choice of true eQTL effects was considered. The mean and s.d. of the relative bias obtained across simulation iterations are provided.

| $\beta_1$ | Lasso | Enet | ALasso | PENA |
|---|---|---|---|---|
| 0.02 | 8 (189) | 9 (192) | -23 (136) | -2 (179) |
| 0.04 | 11 (97) | 13 (99) | -21 (70) | 1 (92) |
| 0.06 | 12 (68) | 14 (69) | -20 (48) | 2 (65) |
| 0.08 | 13 (54) | 15 (55) | -19 (38) | 3 (52) |
| 0.1 | 13 (46) | 15 (47) | -19 (32) | 3 (45) |
| 0.12 | 14 (41) | 15 (42) | -19 (28) | 3 (41) |
| 0.14 | 14 (38) | 16 (39) | -19 (25) | 3 (38) |
| 0.16 | 14 (36) | 16 (36) | -19 (24) | 3 (36) |
| 0.18 | 14 (34) | 16 (35) | -19 (22) | 4 (35) |
| 0.2 | 14 (33) | 16 (34) | -19 (21) | 4 (34) |

Table S5: Square root of mean squared error (RMSE) of the estimates of the effect size of the genetic component of expression on the outcome. The unadjusted TWAS approaches are Lasso, ALasso, and Enet TWAS, depending on the type of penalty function used. PENA denotes the adjusted TWAS approach, PenaTWAS. The heritability of gene expression is 10%. The second choice of true eQTL effects was considered.

| $\beta_1$ | Lasso | Enet | ALasso | PENA |
|---|---|---|---|---|
| 0.02 | 0.04 | 0.04 | 0.03 | 0.04 |
| 0.04 | 0.04 | 0.04 | 0.03 | 0.04 |
| 0.06 | 0.04 | 0.04 | 0.03 | 0.04 |
| 0.08 | 0.04 | 0.05 | 0.03 | 0.04 |
| 0.1 | 0.05 | 0.05 | 0.04 | 0.05 |
| 0.12 | 0.05 | 0.05 | 0.04 | 0.05 |
| 0.14 | 0.06 | 0.06 | 0.04 | 0.05 |
| 0.16 | 0.06 | 0.06 | 0.05 | 0.06 |
| 0.18 | 0.07 | 0.07 | 0.05 | 0.06 |
| 0.2 | 0.07 | 0.07 | 0.06 | 0.07 |

Table S6: Summary of relative bias in estimates of the effect size of the genetic component of expression on the outcome. The unadjusted TWAS approaches are Lasso, ALasso, and Enet TWAS, depending on the type of penalty function used. PENA denotes the adjusted TWAS approach, PenaTWAS. The heritability of gene expression is 20%. The mean and s.d. of the relative bias obtained across simulation iterations are provided.

| $\beta_1$ | Lasso | Enet | ALasso | PENA |
|---|---|---|---|---|
| 0.02 | 8 (166) | 10 (169) | -19 (129) | 1 (164) |
| 0.04 | 9 (85) | 11 (87) | -18 (66) | 2 (85) |
| 0.06 | 9 (59) | 11 (60) | -18 (45) | 2 (59) |
| 0.08 | 9 (47) | 11 (47) | -18 (35) | 2 (47) |
| 0.1 | 9 (39) | 11 (40) | -18 (30) | 2 (40) |
| 0.12 | 9 (35) | 11 (36) | -18 (26) | 2 (36) |
| 0.14 | 9 (32) | 11 (33) | -18 (23) | 2 (33) |
| 0.16 | 9 (30) | 11 (30) | -18 (22) | 2 (31) |
| 0.18 | 9 (28) | 11 (29) | -17 (20) | 2 (30) |
| 0.2 | 10 (27) | 11 (28) | -17 (19) | 2 (29) |

Table S7: Square root of mean squared error (RMSE) of the estimates of the effect size of the genetic component of expression on the outcome. The unadjusted TWAS approaches are Lasso, ALasso, and Enet TWAS, depending on the type of penalty function used. PENA denotes the adjusted TWAS approach PenaTWAS. The heritability of gene expression is 20%.

| $\beta_1$ | Lasso | Enet | ALasso | PENA |
|---|---|---|---|---|
| 0.02 | 0.03 | 0.03 | 0.03 | 0.03 |
| 0.04 | 0.03 | 0.03 | 0.03 | 0.03 |
| 0.06 | 0.04 | 0.04 | 0.03 | 0.04 |
| 0.08 | 0.04 | 0.04 | 0.03 | 0.04 |
| 0.1 | 0.04 | 0.04 | 0.03 | 0.04 |
| 0.12 | 0.04 | 0.04 | 0.04 | 0.04 |
| 0.14 | 0.05 | 0.05 | 0.04 | 0.05 |
| 0.16 | 0.05 | 0.05 | 0.04 | 0.05 |
| 0.18 | 0.05 | 0.06 | 0.05 | 0.05 |
| 0.2 | 0.06 | 0.06 | 0.05 | 0.06 |

Table S8: Summary of relative bias in estimates of the effect size of genetic component of expression on a binary case-control outcome. The unadjusted TWAS approaches are Lasso, ALasso, and Enet TWAS, depending on the type of penalty function used. PENA denotes the adjusted TWAS approach PenaTWAS. The heritability of gene expression is 10%. The mean and s.d. of the relative bias obtained across simulation iterations are provided.

| $\beta_1$ | Lasso | Enet | ALasso | PENA |
|---|---|---|---|---|
| 0.02 | 17 (347) | 23 (352) | -32 (191) | 7 (356) |
| 0.04 | 15 (180) | 20 (185) | -32 (97) | 7 (185) |
| 0.06 | 15 (127) | 19 (131) | -32 (67) | 6 (130) |
| 0.08 | 14 (102) | 18 (105) | -32 (52) | 6 (105) |
| 0.1 | 14 (87) | 18 (91) | -32 (43) | 6 (90) |
| 0.12 | 14 (79) | 18 (82) | -32 (38) | 6 (81) |
| 0.14 | 14 (73) | 17 (76) | -32 (34) | 6 (76) |
| 0.16 | 14 (69) | 17 (72) | -32 (32) | 6 (71) |
| 0.18 | 14 (66) | 17 (69) | -32 (30) | 6 (69) |
| 0.2 | 14 (63) | 17 (66) | -32 (28) | 6 (66) |

Table S9: Square root of mean squared error (RMSE) in the estimates of the effect size of the genetic component of expression on a binary case-control outcome. The unadjusted TWAS approaches are Lasso, ALasso, and Enet TWAS, depending on the type of penalty function used. PENA denotes the adjusted TWAS approach PenaTWAS. The heritability of gene expression is 10%.

| $\beta_1$ | Lasso | Enet | ALasso | PENA |
|---|---|---|---|---|
| 0.02 | 0.07 | 0.07 | 0.04 | 0.07 |
| 0.04 | 0.07 | 0.07 | 0.04 | 0.07 |
| 0.06 | 0.08 | 0.08 | 0.04 | 0.08 |
| 0.08 | 0.08 | 0.09 | 0.05 | 0.08 |
| 0.1 | 0.09 | 0.09 | 0.05 | 0.09 |
| 0.12 | 0.1 | 0.1 | 0.06 | 0.1 |
| 0.14 | 0.1 | 0.11 | 0.07 | 0.11 |
| 0.16 | 0.11 | 0.12 | 0.07 | 0.11 |
| 0.18 | 0.12 | 0.13 | 0.08 | 0.12 |
| 0.2 | 0.13 | 0.14 | 0.08 | 0.13 |

Table S10: Number of genes significantly associated with the various phenotypes analyzed in the real data application based on a more stringent threshold of 0.05/20000. Here, PenaTWAS is an adjusted approach, and Enet TWAS and Otters are unadjusted approaches.

| | PenaTWAS | Enet TWAS | Otters |
|---|---|---|---|
| Height | 88 | 105 | 118 |
| LDL | 17 | 20 | 30 |
| HDL | 27 | 31 | 40 |
| Triglycerides | 24 | 18 | 25 |

Table S11: Genes associated with HDL cholesterol identified by both PenaTWAS and Enet TWAS. PenaTWAS is an adjusted TWAS approach, and Enet TWAS is an unadjusted TWAS approach.

| Chr | Gene | Chr | Gene |
|---|---|---|---|
| 1 | CTSS | 11 | NR1H3 |
| 1 | GSTM1 | 11 | SLC22A18 |
| 1 | GSTM2 | 11 | TAGLN |
| 1 | DR1 | 11 | ACP2 |
| 1 | NT5C1A | 12 | TCTN2 |
| 2 | IMMT | 15 | NDUFAF1 |
| 2 | TGOLN2 | 15 | PRKXP1 |
| 4 | DGKQ | 16 | SMPD3 |
| 6 | PEX6 | 17 | ORMDL3 |
| 7 | ZFAND2A | 17 | TRIM65 |
| 10 | ADRB1 | 17 | CCR10 |
| 11 | MRPL21 | 17 | CNTNAP1 |
| 11 | SNX32 | 17 | PLEKHH3 |

Table S12: Genes associated with HDL cholesterol identified by Enet TWAS but overlooked by PenaTWAS (black font) and genes identified by PenaTWAS but missed by Enet TWAS (blue font). Enet TWAS is an unadjusted TWAS approach, and PenaTWAS is an adjusted TWAS approach.

| Chr | Gene | Enet P | PENA P | Reported | Other important phenotypes |
|---|---|---|---|---|---|
| 3 | MKRN2 | 1.53E-05 | 0.07 | N | LDL cholesterol, Total cholesterol level, etc. |
| 6 | GNMT | 3.99E-11 | 0.02 | Y | |
| 6 | NT5DC1 | 1.22E-06 | 0.0001 | Y | |
| 7 | PILRB | 1.06E-05 | 0.02 | N | Dementia, hemoglobin level, etc. |
| 10 | OR13A1 | 2.46E-06 | 0.0005 | N | Beta-microseminoprotein levels, etc. |
| 11 | C1QTNF4 | 2.22E-13 | 6.28E-05 | N | BMI, height, Cardiovascular disease, etc. |
| 11 | IGHMBP2 | 4.06E-08 | 0.003 | Y | |
| 12 | MMAB | 8.82E-06 | 0.001 | Y | |
| 19 | ETHE1 | 1.63E-05 | 0.0004 | N | LYPD3 protein levels, etc. |
| 19 | EEF2 | 1.53E-06 | 0.5 | N | Height |
| 19 | QPCTL | 2.58E-07 | 0.0003 | N | BMI, LDL cholesterol levels, etc. |
| 19 | PNPLA6 | 2.06E-05 | 0.0002 | N | vaginal microbiome measures, RETN levels, etc. |
| 20 | GSS | 1.74E-19 | 0.02 | N | Schizophrenia, Serum urate levels, etc. |
| 20 | PLTP | 3.84E-14 | 0.02 | Y | |
| 3 | NT5DC2 | 0.001 | 3.34E-23 | N | Waist to hip ratio, Hemoglobin, Autism, etc. |
| 5 | ERAP2 | 0.88 | 1.62E-07 | N | Hypertension, Cardiovascular disease, etc. |
| 5 | SETD9 | 0.02 | 2.62E-06 | N | Triglyceride to HDL ratio, Breast cancer, etc. |
| 7 | TMEM120A | 0.25 | 1.30E-08 | N | Mean platelet volume, Sarcoidosis, etc. |
| 9 | CEL | 0.003 | 1.94E-05 | N | Serum levels of protein CEL, Type 1 diabetes. |
| 11 | VEGFB | 0.006 | 3.41E-08 | Y | |
| 19 | ZNF155 | 3.90E-05 | 1.62E-05 | N | protein measurement, PHF-tau measurement |

Table S13: Genes associated with LDL cholesterol identified by both PenaTWAS and Enet TWAS. PenaTWAS is an adjusted TWAS approach, and Enet TWAS is an unadjusted TWAS approach.

| Chr | Gene | Chr | Gene |
|---|---|---|---|
| 1 | GSTM4 | 17 | MRPL45P2 |
| 1 | SORT1 | 19 | ZNF155 |
| 1 | RHD | 19 | ZNF554 |
| 3 | WDR5B | 19 | ALDH16A1 |
| 11 | ZDHHC24 | 19 | SLC25A42 |
| 11 | TAGLN | 19 | SMARCA4 |
| 15 | ULK3 | 19 | KEAP1 |
| 16 | MARVELD3 | 19 | ZNF221 |
| 17 | TBKBP1 | 22 | DNAJB7 |

Table S14: Genes associated with LDL cholesterol identified by Enet TWAS but overlooked by PenaTWAS (black font) and genes identified by PenaTWAS but missed by Enet TWAS (blue font). Enet TWAS is an unadjusted TWAS approach, and PenaTWAS is an adjusted TWAS approach.

| Chr | Gene | Enet P | PENA P | Reported | Other important phenotypes |
|---|---|---|---|---|---|
| 2 | NRBP1 | 9.98E-07 | 4.05E-05 | Y | |
| 5 | ENC1 | 9.55E-08 | 7.34E-05 | N | Height, metabolite levels, etc. |
| 6 | NT5DC1 | 1.08E-05 | 0.0003 | Y | |
| 7 | UFSP1 | 2.10E-06 | 0.0003 | N | Height, resting heart rate, etc. |
| 9 | INPP5E | 6.36E-06 | 0.0002 | Y | |
| 15 | MPI | 1.22E-06 | 0.0002 | Y | |
| 19 | IL12RB1 | 1.57E-06 | 0.001 | N | Systemic sclerosis, Percent liver fat |
| 20 | ABHD12 | 8.30E-07 | 0.03 | N | Hemoglobin measurement, lung function, etc |
| 5 | ERAP2 | 0.66 | 7.31E-08 | N | Hypertension, cardiovascular disease, etc. |
| 5 | TMEM161B-AS1 | 0.11 | 6.38E-27 | N | BMI, blood pressure, etc. |
| 7 | TMEM120A | 0.56 | 4.47E-06 | N | Mean platelet volume, Sarcoidosis, etc. |
| 10 | SLC25A16 | 0.17 | 2.24E-07 | N | Drug dependence, Sarcoidosis, etc. |
| 15 | SPTBN5 | 0.003 | 1.42E-09 | N | BMI, cardiovascular disease, etc. |
| 17 | SKAP1 | 9.61E-05 | 1.24E-05 | Y | *Pleiotropy among LDL, HDL, triglycerides |
| 19 | SYMPK | 0.0002 | 9.77E-29 | Y | |

Table S15: Genes associated with triglycerides identified by both PenaTWAS and Enet TWAS. PenaTWAS is an adjusted TWAS approach, and Enet TWAS is an unadjusted TWAS approach.

| Chr | Gene | Chr | Gene |
|---|---|---|---|
| 2 | ATRAID | 12 | YEATS4 |
| 2 | NRBP1 | 15 | SERF2 |
| 2 | IFT172 | 15 | ELL3 |
| 3 | NISCH | 17 | SLFN5 |
| 6 | SNHG5 | 17 | MRPS7 |
| 8 | FAM167A | 17 | CCR10 |
| 10 | REEP3 | 17 | CNTNAP1 |
| 11 | MRPL21 | 19 | SLC25A42 |
| 11 | OVOL1 | 19 | PNPLA6 |
| 11 | TAGLN | 22 | TTC38 |

Table S16: Genes associated with triglycerides identified by Enet TWAS but overlooked by PenaTWAS (black font) and genes identified by PenaTWAS but missed by Enet TWAS (blue font). Enet TWAS is an unadjusted TWAS approach, and PenaTWAS is an adjusted TWAS approach.

| Chr | Gene | Enet P | PENA P | Reported | Other phenotypes |
|---|---|---|---|---|---|
| 1 | CTSS | 3.25E-06 | 0.0001 | N | Renal function |
| 1 | DR1 | 8.31E-06 | 0.003 | Y | |
| 2 | TLK1 | 9.78E-06 | 0.05 | N | Platelet count, autism, etc. |
| 4 | PDE5A | 1.60E-05 | 0.07 | N | Anemia, Biopolar disorder, etc. |
| 7 | CCDC126 | 1.17E-05 | 0.04 | N | FEV1, serum alkaline phosphatase levels, etc. |
| 13 | UPF3A | 9.19E-06 | 3.36E-05 | N | Blood pressure, cardiovascular disease, etc. |
| 16 | CLEC18A | 3.04E-05 | 0.0001 | N | BMI, Idiopathic knee osteoarthritis |
| 17 | ALKBH5 | 7.96E-06 | 0.0004 | N | LDL cholesterol, total cholesterol level, etc. |
| 17 | RNFT1 | 1.71E-05 | 0.0003 | N | Cystatin C production |
| 20 | PLTP | 2.52E-11 | 0.2 | Y | |
| 1 | WDR77 | 0.4 | 1.00E-08 | N | LDL cholesterol, total cholesterol, etc. |
| 3 | NT5DC2 | 0.001 | 4.05E-32 | N | Hemoglobin, brain measurement, etc. |
| 7 | TMEM120A | 0.007 | 6.19E-12 | N | Mean platelet volume, Sarcoidosis |
| 8 | BLK | 0.01 | 1.18E-09 | Y | |
| 8 | RPL8 | 0.1 | 2.25E-05 | N | Diamond-Blackfan Anemia, etc. |
| 8 | MFHAS1 | 0.2 | 6.83E-06 | Y | |
| 10 | BAMBI | 0.3 | 2.63E-08 | N | Height, blood protein measurement, etc. |
| 10 | SLC25A16 | 0.02 | 1.04E-09 | N | Cognitive function, brain morphology, etc. |
| 11 | VEGFB | 0.006 | 2.24E-06 | Y | |
| 11 | DSCAML1 | 0.75 | 2.60E-08 | Y | |
| 17 | GPATCH8 | 0.002 | 1.58E-19 | Y | |
| 19 | SYMPK | 0.004 | 1.44E-05 | N | LDL cholesterol, total cholesterol, etc. |

Table S17: Genes associated with height identified by both PenaTWAS and Enet TWAS. PenaTWAS is an adjusted TWAS approach, and Enet TWAS is an unadjusted TWAS approach.

| Chr | Gene | Chr | Gene | Chr | Gene | Chr | Gene |
|---|---|---|---|---|---|---|---|
| 1 | GBP3 | 5 | B4GALT7 | 11 | XRRA1 | 17 | KANSL1-AS1 |
| 1 | INPP5B | 5 | CDO1 | 11 | C1QTNF4 | 17 | TBKBP1 |
| 1 | FHL3 | 5 | CEP120 | 11 | LRFN4 | 17 | RAPGEFL1 |
| 1 | KIF1B | 6 | SMPD2 | 11 | ZDHHC24 | 17 | LRRC37A |
| 1 | PSMD4 | 6 | AK9 | 11 | FDX1 | 17 | CTC1 |
| 1 | B3GALNT2 | 6 | UHRF1BP1 | 11 | ACP2 | 17 | FN3KRP |
| 2 | LINC00471 | 6 | TCP11 | 11 | CCS | 17 | SKAP1 |
| 2 | ATP6V1E2 | 6 | EPB41L2 | 12 | YEATS4 | 17 | DDX42 |
| 2 | FAM136A | 6 | MAPK14 | 12 | GATC | 17 | LRRC37A4P |
| 2 | VAMP5 | 6 | NT5DC1 | 12 | TCTN2 | 17 | COPRS |
| 2 | CRIPT | 7 | GNA12 | 13 | CYSLTR2 | 17 | ADAM11 |
| 2 | CNPPD1 | 7 | SLC13A4 | 13 | CDC16 | 17 | USP36 |
| 2 | NRBP1 | 8 | ZFAT-AS1 | 13 | RCBTB1 | 17 | MNT |
| 2 | IFT172 | 8 | DSCC1 | 14 | RAB15 | 19 | ZNF561 |
| 2 | VAMP8 | 8 | RPS20 | 15 | NDUFAF1 | 19 | TULP2 |
| 3 | ANAPC13 | 8 | OTUD6B-AS1 | 15 | ISL2 | 20 | CPNE1 |
| 3 | KLHDC8B | 9 | LINC00476 | 16 | TBX6 | 20 | PROCR |
| 3 | SENP7 | 9 | HABP4 | 16 | SYNGR3 | 20 | GSS |
| 3 | TMA7 | 9 | ERCC6L2 | 16 | CLEC18A | 22 | PACSIN2 |
| 3 | KIAA1143 | 9 | SNHG7 | 16 | CDK10 | 22 | RIBC2 |
| 4 | GUSBP5 | 9 | MEGF9 | 16 | CDIP1 | 22 | SLC25A1 |
| 4 | MED28 | 9 | C9orf43 | 16 | MARVELD3 | 22 | TTC38 |
| 4 | BBS7 | 9 | INPP5E | 16 | CLEC18C | 22 | RNF215 |
| 4 | CCNA2 | 9 | C9orf64 | 16 | CPNE7 | 22 | CBY1 |
| 5 | SETD9 | 10 | STOX1 | 16 | NOXO1 | | |
| 5 | CAMLG | 10 | ACTR1A | 17 | GJC1 | | |

Table S18: Genes associated with height identified by Enet TWAS but overlooked by PenaTWAS (black font) and genes identified by PenaTWAS but missed by Enet TWAS (blue font). Enet TWAS is an unadjusted TWAS approach, and PenaTWAS is an adjusted TWAS approach.

| Chr | Gene | Enet P | pena P | Chr | Gene | Enet P | pena P |
|---|---|---|---|---|---|---|---|
| 1 | ACP6 | 4.77E-07 | 5.30E-05 | 15 | TICRR | 3.17E-06 | 0.0005 |
| 1 | AMIGO1 | 1.61E-05 | 6.56E-05 | 16 | THUMPD1 | 2.85E-08 | 0.001 |
| 1 | MEAF6 | 9.06E-06 | 0.0003 | 17 | SHMT1 | 3.66E-06 | 8.82E-05 |
| 1 | TSEN15 | 1.26E-17 | 3.74E-05 | 17 | MRPL45P2 | 1.35E-06 | 0.002 |
| 1 | ARV1 | 6.08E-08 | 0.007 | 17 | CCDC103 | 1.51E-06 | 6.14E-05 |
| 1 | CNKSR1 | 3.62E-07 | 0.03 | 17 | ALKBH5 | 8.77E-06 | 0.0003 |
| 2 | PFN4 | 8.15E-06 | 0.005 | 17 | CRLF3 | 4.56E-09 | 0.002 |
| 3 | NT5DC2 | 2.29E-05 | 0.009 | 19 | LRRC25 | 5.41E-06 | 0.0002 |
| 3 | TIPARP-AS1 | 1.25E-05 | 0.0001 | 19 | SLC25A42 | 2.07E-05 | 0.0009 |
| 4 | GRPEL1 | 1.36E-05 | 0.0009 | 19 | ETHE1 | 4.21E-07 | 0.0001 |
| 4 | SLC2A9 | 2.48E-06 | 0.007 | 22 | SYNGR1 | 1.73E-05 | 4.89E-05 |
| 5 | PCYOX1L | 1.69E-05 | 0.39 | 1 | LRRC8C | 0.016 | 5.62E-08 |
| 6 | KIAA1586 | 1.40E-05 | 0.0004 | 1 | WDR77 | 0.43 | 2.50E-07 |
| 7 | MRPS33 | 3.79E-08 | 8.09E-05 | 2 | TMBIM1 | 0.02 | 1.12E-06 |
| 8 | FAM167A | 2.97E-06 | 0.0004 | 2 | MAP3K2 | 0.007 | 3.20E-05 |
| 8 | SMIM18 | 6.77E-07 | 0.06 | 4 | FGFRL1 | 4.53E-05 | 3.67E-09 |
| 9 | DNLZ | 1.05E-05 | 0.002 | 4 | CISD2 | 0.6 | 1.31E-10 |
| 10 | WAC-AS1 | 8.30E-08 | 0.0003 | 5 | NNT | 0.2 | 1.81E-06 |
| 10 | SLC25A16 | 2.55E-10 | 0.7 | 5 | TMEM161B-AS1 | 0.6 | 3.03E-05 |
| 10 | AS3MT | 3.12E-06 | 5.04E-05 | 6 | TMEM200A | 0.0002 | 1.57E-05 |
| 11 | PGM2L1 | 6.19E-08 | 6.85E-05 | 9 | SUSD1 | 0.002 | 3.27E-05 |
| 11 | POLA2 | 9.62E-06 | 0.003 | 12 | CDK2AP1 | 0.003 | 1.07E-14 |
| 11 | NR1H3 | 2.60E-10 | 0.007 | 15 | PRKXP1 | 0.03 | 1.91E-06 |
| 11 | ZDHHC5 | 2.31E-08 | 4.80E-05 | 15 | NEO1 | 0.0007 | 4.17E-07 |
| 11 | MED19 | 6.16E-08 | 0.0003 | 17 | GPATCH8 | 9.42E-05 | 1.47E-30 |
| 13 | LINC00462 | 2.97E-06 | 0.0004 | 17 | CPD | 0.8 | 5.97E-13 |
| 13 | ITM2B | 6.13E-07 | 0.0003 | 17 | ELP5 | 0.001 | 1.33E-07 |
| 14 | FBLN5 | 8.53E-08 | 0.3 | 19 | PLA2G4C | 3.46E-05 | 3.25E-05 |
| 14 | ACYP1 | 1.15E-05 | 0.0006 | 22 | DNAJB7 | 3.78E-05 | 1.31E-06 |
